# From egg to adult: comprehensive (e)DNA metabarcoding monitoring of fish diversity in a temperate estuary

**DOI:** 10.1101/2025.10.12.681914

**Authors:** André O. Ferreira, Cláudia Machado, Olga M. Azevedo, Cristina Barroso, Sofia Duarte, Conceição Egas, A. Miguel Piecho-Santos, Filipe O. Costa

## Abstract

Monitoring fish species through ichthyoplankton surveys provides important information for fish stock assessment and management. To test the effectiveness of DNA metabarcoding to identify fish species and to capture seasonal variations in local ichthyofauna, monthly ichthyoplankton and 2 L water samples were collected over 13 months in the lower section of the Guadiana River Estuary in southeast Portugal. Both sample types underwent high-throughput sequencing for three genetic markers (COI, 12S, 16S), with morphological identification also performed for ichthyoplankton. Bulk and water samples identified a total of 131 fish species throughout the year. DNA metabarcoding demonstrated higher taxonomic resolution and diversity detection, with 115 species recovered, while morphology identified only 23 species. Ichthyoplankton metabarcoding also detected 40% more fish than water eDNA, recovering almost the double of species despite the fact that both approaches used the same metabarcoding primers. The integration of multiple molecular markers was crucial to maximize diversity detection in both DNA-based methods. In addition, DNA metabarcoding was able to identify ichthyofaunal spawning periods and captured significant seasonal variations in fish community, with higher diversity observed during the warmer months. With this strategy, around 66% of the historically recorded ichthyoplankton taxa in the region were identified, along with several new records. The findings demonstrated the capability of (e)DNA metabarcoding to uncover seasonal variations in the regional fish community, provided new insights on the ichthyofauna of the Guadiana Estuary, and revealed the need for more in-depth studies to improve the efficiency of multiple sampling methods for fish species identification.

## Introduction

The seasonal dynamics of ichthyoplankton in estuaries are unique and complex, so it is essential to understand how these communities change over time to implement effective measures for fisheries management and conservation (Zhang et al., 2022). These habitats are biologically rich, serving as critical spawning and nursery areas for many fish species, being at the same time very vulnerable ecosystems (Kennish, 2002). Fish stock assessments rely on the monitoring of ichthyoplankton (Botsford et al., 2009), as data collected on the distribution and abundance of fish eggs and larvae gives insights into reproductive success, community structure, and the overall health of fish stocks (Ré, 1999; Reuben, 1981; Santos et al., 2018). Additionally, ichthyoplankton variation throughout the year provides valuable information into spawning periods, recruitment, and species migration, which are directly influenced by changes in multiple environmental factors such as temperature, salinity fluctuations, and nutrient availability (Sloterdijk et al., 2017; Tackx et al., 2004; Zhang et al., 2016). Despite their ecological significance, many estuaries still lack long-term studies focused on ichthyoplankton communities, which compromises their conservation and management efforts (Zhang et al., 2022).

The Guadiana River Estuary, a natural border between Portugal and Spain, is one of these important ecosystems, representing a central transitional zone in the region that serves as a breeding and nursery area for a variety of fish species, both resident and migratory, due to the available habitats and shelter, as well as the abundant food resources essential for the early life stages (Beck et al., 2001). Several ichthyoplankton studies were carried out in the Guadiana Estuary, exploring multiple subjects including ichthyoplankton composition (Chícharo et al., 2000; Chícharo & Teodósio, 1991; Chícharo et al., 2006; Faria et al., 2006), larval abundance and nutritional conditions (Esteves et al., 2000), the hydro-ecological effects of the Alqueva dam on fish species (Chícharo et al., 2001), and the impacts of environmental and anthropogenic factors on the estuary (Miró et al., 2020). All these studies generated important data about the ecosystem, but in some cases, more than a decade has passed since they were conducted, and the research relied exclusively on the morphological inspection of the eggs and larvae. It is necessary to integrate new molecular techniques for the assessment of this ecosystem, as they can increase the taxonomic resolution and the precision of these investigations, address more efficiently the current ecological challenges and improve our knowledge about the ichthyoplankton dynamics in this estuarine system.

The traditional approach to monitoring ichthyoplankton relies on the morphological identification of the eggs and larvae, a valuable method still used today but characterized by several limitations. Usually, the entire inspection process is time-consuming and low-throughput, demanding specific taxonomic expertise that can only be acquired during years of experience (Ko et al., 2013; Nobile et al., 2019; Yu et al., 2012). Even with all this knowledge, precise identifications are a challenging achievement, since it is frequent for these early life stages to display lack of distinctive physical traits or to be physically damaged during sampling (Darling & Blum, 2007; Hulley et al., 2018; Reynalte-Tataje et al., 2012). DNA-based techniques, such as DNA metabarcoding and environmental DNA (eDNA) (Hajibabaei et al., 2011; Taberlet et al., 2012), have been implemented in this field in order to overcome the limitations of morphology, constituting a valuable complementary method or an alternative with innovative solutions (Ferreira et al., 2024; Shokralla et al., 2012). The high-throughput capacity of metabarcoding is revolutionary for species identification in ichthyoplankton samples, allowing more frequent and cost-effective sampling events that benefit studies conducted across broader temporal scales (Nobile et al., 2019; Porter & Hajibabaei, 2018; Stefanni et al., 2018). These molecular studies can employ multiple genetic markers and primer sets instead of a single-marker approach, creating a complementary effect that enhances the diversity recovery and provides a more detailed profile of the community structure (Duke & Burton, 2020; Ferreira et al., 2024; Hollatz et al., 2017; Stefanni et al., 2018; Zhang et al., 2018).

In the context of eDNA analysis, water collection represents a non-invasive and sensitive method for detecting the presence of fish species (Jackman et al., 2021; Keck et al., 2022; Peterson et al., 2022). Both methodologies can be used in parallel, each with its own advantages and limitations, and requiring specific sampling and sequencing strategies (Van Nynatten et al., 2023). Bulk ichthyoplankton metabarcoding naturally targets the reproducing species within the ecosystem, collecting fish eggs and larvae that provide information on seasonal and spatial variations in the communities while accounting for the unpredictable distribution of ichthyoplankton (Parrish et al., 1981; Pritt et al., 2015; Rodriguez, 2019). In contrast, eDNA has the potential to detect organisms across all life stages, from eggs to adults, transient species and a wide range of biodiversity that ichthyoplankton samples cannot capture (Gold et al., 2021; Pont et al., 2018). Simultaneously, this clear advantage can also represent a limitation for fish stock assessments, since the identified taxa cannot be linked to specific developmental stages, a drawback that also applies to ichthyoplankton metabarcoding. Another important difference that can influence species identification between these molecular techniques is the DNA quality and integrity retrieved by each method, with eDNA samples often containing degraded or low-quality DNA, in contrast to the higher-quality DNA obtained from bulk ichthyoplankton samples, the ideal template for molecular analyses (Harrison et al., 2019; van der Loos & Nijland, 2021). Despite these specificities, the combined use of both methods creates an advanced approach to achieve precise species identification in estuarine ecosystems, detect shifts in species distributions, and complement all the information generated by traditional morphological identification in previous studies.

The primary objective of this study is to achieve species-level identification of fish communities in the Guadiana Estuary over an annual cycle, employing a multi-component approach that combines morphological identification of fish eggs and larvae, ichthyoplankton DNA metabarcoding, and eDNA analysis from water samples. This research aims to identify local ichthyofauna and their respective spawning periods using ichthyoplankton data, and to uncover seasonal variations in the local fish community by assessing species occurrence obtained using both sampling methods. By comparing the efficiency of molecular approaches, the study also examines the strengths and limitations of DNA metabarcoding and eDNA in capturing fish diversity, using a multi-marker approach to minimize misidentifications, false positives, and false negatives. Finally, this study will enhance the baseline knowledge of fish diversity and seasonal dynamics within the Guadiana estuary, delivering essential data for conservation and fisheries management.

## Methods

### Study area

The region of interest in this study was the Guadiana River Estuary, a mesotidal watercourse located along the southern border of Portugal and Spain. It is a dynamic and important system in the Iberian Peninsula that connects the Guadiana River to the Atlantic Ocean, with approximately 70 km in length and an average tidal amplitude of 2 meters. This system can be subdivided into three distinct sections: the upper freshwater region, the middle salinity transition zone, and the lower marine-dominated section. This characterization is made based on the variations in salinity, vegetation, and sedimentology across this aquatic system (Chícharo et al., 2001; Morales González, 2017), and according to frequently employed methods for estuarine subdivision (Olausson & Cato, 1980). A clear salinity gradient characterizes the estuary, with the upper section maintaining salinity levels close to zero (<0.5), the middle one representing a transitional brackish zone (0.5-25), and the lower section, closer to the river mouth, exhibiting salinity levels comparable to seawater (>25) (Chícharo et al., 2006). The inflow of freshwater in the system exhibits seasonal and annual variability, with an irregular hydrological regime that includes severe drought and flooding periods. Moreover, the Guadiana Estuary is strongly regulated by the Alqueva Dam, which controls freshwater discharge and influences the estuarine dynamics. These conditions make the area a unique location for ecological studies, particularly under the influence of climatic and anthropogenic pressures.

### Sample collection

Between April 2022 and April 2023, monthly sampling was conducted in the lower section of the Guadiana River Estuary, with an additional month added to the year of sampling to gain preliminary insights into potential interannual differences in the fish community. Three points within this area were selected as replicates, representing this section of the estuary rather than distinct sampling sites (Figure 1; Table S.1). Ichthyoplankton samples were collected during flood tide, two hours before high tide, using a conical plankton net with a mesh size of 500 μm and a diameter of 60 cm. Consecutive horizontal tows were conducted in the middle of the river channel at a subsurface level for 5 minutes. Samples from each point were randomly fixed with 4% formalin for morphological inspection and 96% ethanol or higher for molecular analysis immediately after collection. The samples chosen for molecular analyses were stored at -20 °C until DNA extraction was performed. Additionally, the volume of water filtered was measured using a flowmeter placed at the mouth of the net. For the environmental DNA analysis, water samples totaling 12 L per month (two 2 L samples from each of the three sampling points) were collected at approximately 0.5 m depth and immediately stored under cooled conditions until filtration. Water samples were filtered in the laboratory as soon as possible to minimize DNA degradation, using 47 mm diameter, 0.45 μm pore-sized filters (mixed cellulose esters, Merck-Millipore, Burlington, MA, USA), and stored at -20°C until further processing. Negative controls, consisting of one 2 L flask, containing MilliQ water, were used throughout the workflow (sampling, storage, sample processing in the lab, DNA extraction and PCR amplification) and processed as the water samples. 0.45 μm pore-sized filters were also used to filter the preservative ethanol from the ichthyoplankton samples. All sampling procedures complied with ethical standards established by the former General Direction of Veterinary (DGV) under the former Ministry of Agriculture, Rural Development, and Fisheries in Lisbon, Portugal, for ichthyoplankton collection and ecological research.

**Figure 1.**
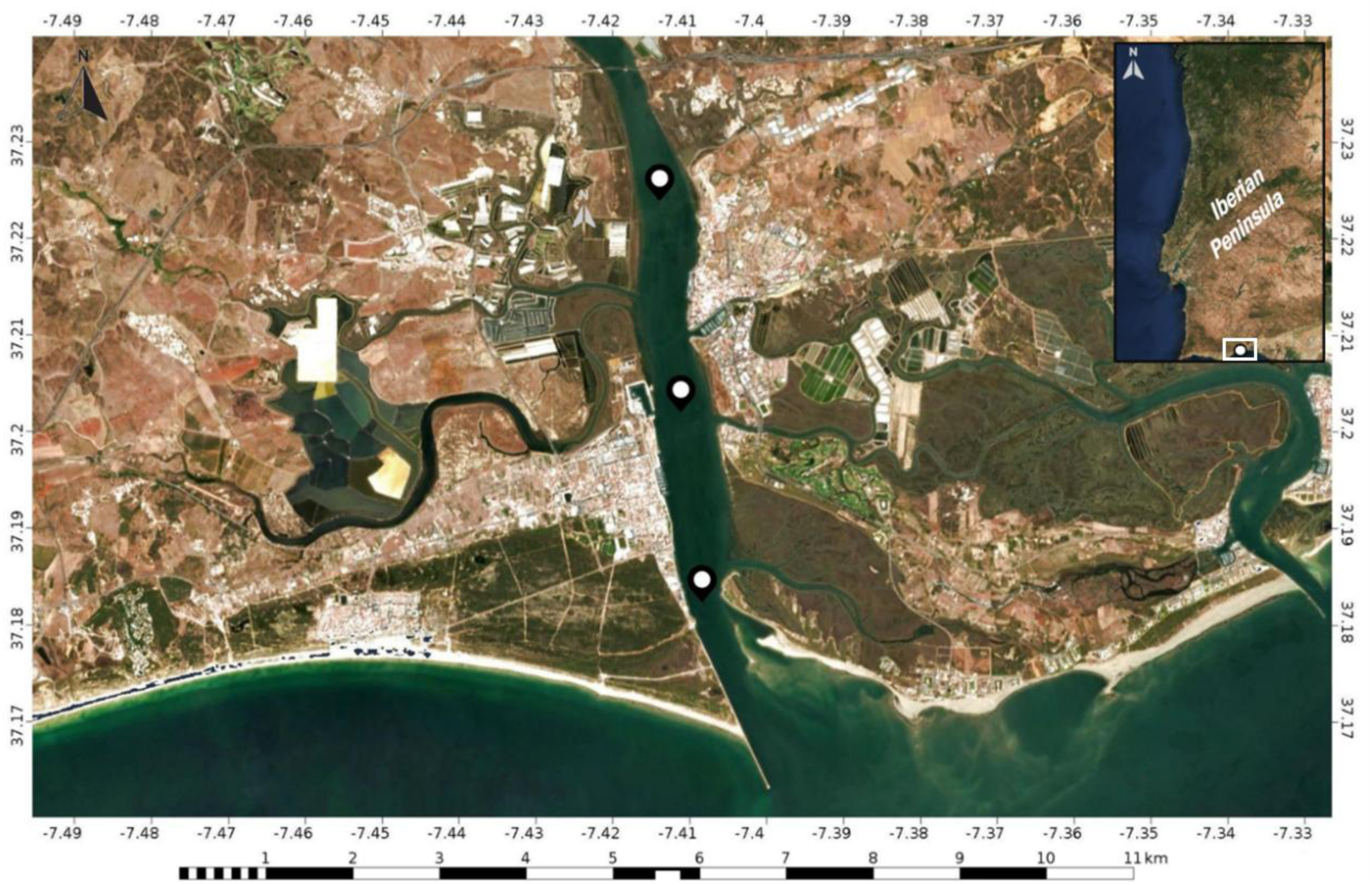
Sampling locations from the lower section of the Guadiana River Estuary, located in southeastern Portugal. Maps were created using the CalTopo application (https://caltopo.com; CalTopo LLC, 2021).

### Morphological processing

Formalin-preserved samples were inspected three days after collection to ensure the fixation process was complete. A Leica S8 APO stereomicroscope (Leica Microsystems, Wetzlar, Germany) was used for ichthyoplankton morphological identification. Samples were sorted to isolate fish eggs and larvae, and the total numbers were then counted. The identification of eggs was reliable and feasible only for sardines (*Sardina pilchardus*) and anchovies (*Engraulis encrasicolus*), as the morphology-based approach did not provide enough resolution for other taxa that lacked distinguishing characteristics. Fish larvae were identified to the lowest possible taxonomic level based on their morphological traits, following region-specific guides for identifying planktonic eggs and larval stages (Ré, 1999; Ré & Meneses, 2008; Rodriguez et al., 2017).

### Molecular processing

#### DNA Extraction

The DNA from the bulk ichthyoplankton samples was extracted following an adapted protocol described by Steinke et al. (2022) and based on Ivanova et al. (2006). After ethanol filtration using 0.45 μm pore size filters (S-Pak Filters, Millipore), samples were incubated overnight with agitation at 60 rpm and 56 °C in a lysis buffer (100 mM NaCl, 50 mM Tris-HCl pH 8.0, 10 mM EDTA pH 8.0, and 0.5% SDS) to promote cell lysis while preserving the integrity of each specimen. From the lysate, two 1 mL aliquots were transferred to microtubes for the extraction, mixed with a Binding Mix (6M GuSCN, 20 mM EDTA pH 8.0, 10 mM Tris-HCl pH 6.4 and 4% Triton X-100) and purified during 3 washing steps. In the end, the aliquots containing the eluted DNA were combined in a total volume of 60 µL (30 µL from each aliquot), and DNA concentrations were measured using the Thermo Scientific™ NanoDrop™ One/OneC spectrophotometer (ThermoFisher Scientific, Waltham, MA, USA) to assess the retention of sufficient target DNA before PCR amplification. For the eDNA samples, DNA extraction was performed from the filters used to sieved the water using the DNeasy PowerSoil Kit (Qiagen, Hilden, Germany) following the manufacturer’s protocol. Each filter was divided into two sections and processed separately from each half to maximize DNA yield. Similar to the bulk samples, the final eluted extracts were pooled to create a single sample of 60 µL (30 µL x 2) for downstream analysis. Negative controls were included throughout the DNA extraction protocols to check for possible contamination of the solutions and labware materials used. These negative controls served as templates in the subsequent PCR amplification reactions.

### PCR Amplification and High-Throughput Sequencing

Ichthyoplankton and water samples were processed using the same workflow from this step until the protocol was completed. DNA amplification targeted three mitochondrial gene regions: the cytochrome c oxidase I (COI) gene using two primer sets, the generalist primer pair mlCOIintF/LoboR1 targeting marine metazoans (Leray et al., 2013; Lobo et al., 2013), and the fish-specific primer cocktail, FishATL_Cocktail2 (Costa, 2021); the 12S rRNA gene, amplified with a primer set formed by the primer pairs miFISH-U and miFISH-E (Miya et al., 2015), and the 16S mitochondrial gene, amplified with the Fish 16S primer pair described by Berry et al. (2017) (Table 1). Each sample was submitted to two PCR replicates per primer set using the KAPA HiFi HotStart PCR Kit (Kapa Biosystems, Cape Town, South Africa), according to manufactureŕs guidelines that ensure optimal reaction conditions and high-fidelity amplification.

**Table 1.**
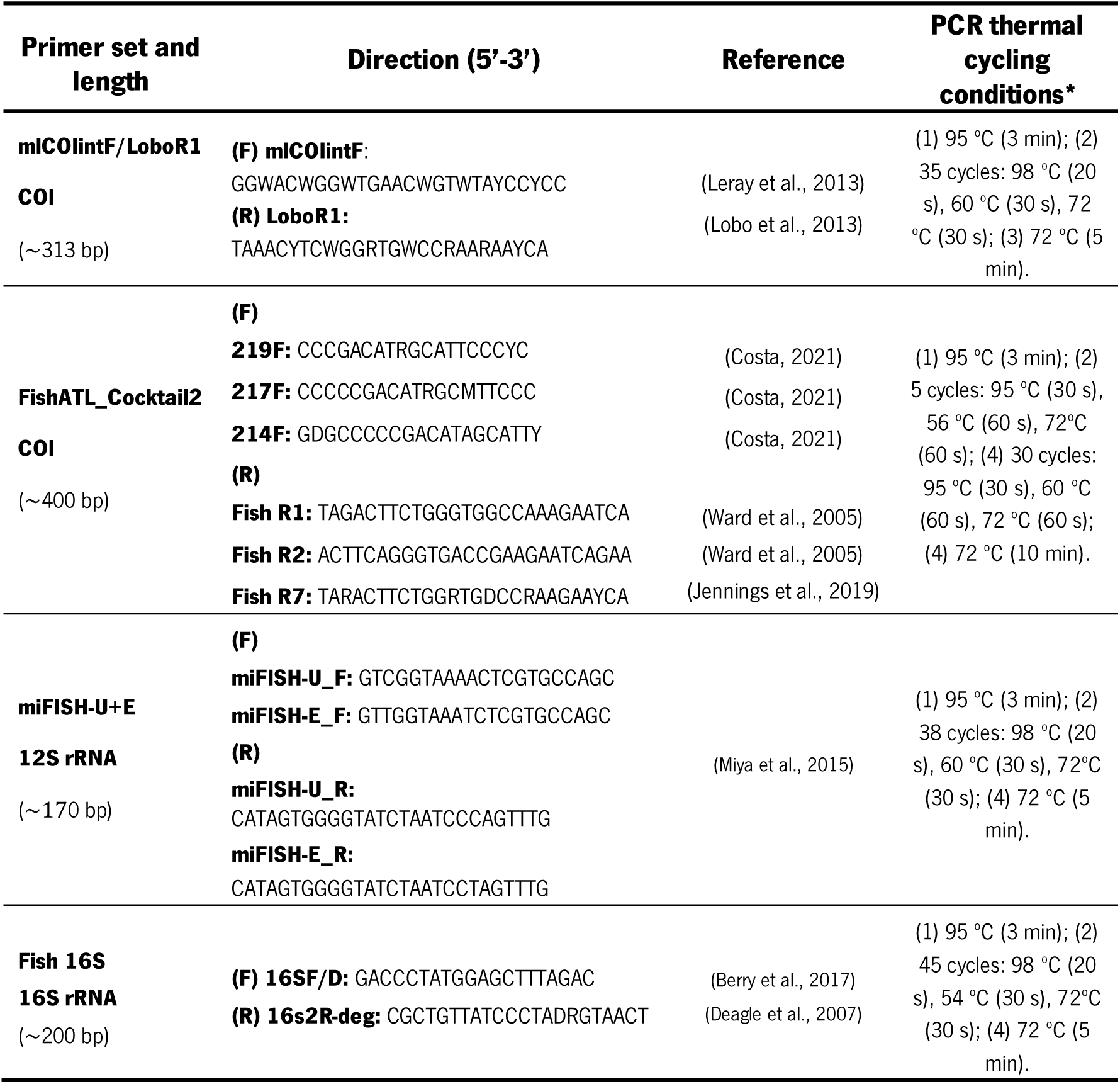
PCR primer pairs/sets used for amplifying the different mitochondrial gene regions and respective PCR thermal cycling conditions. (F) - forward; (R) - reverse; bp - base pairs; “Consensus region” refers to a conserved primer tail sequence preceding the region-specific primer sequence.

All PCR reactions were carried out in a total volume of 25 µL containing the corresponding forward and reverse primers, template DNA, and standard reaction components. Primer concentrations and template volumes varied depending on the primer set: 0.3 µM of each primer with 5 µL of template DNA for the COI (mlCOIintF/LoboR1) and 16S (16SF/D– 16s2R-deg) primers; 3 µM of each primer with 1 µL of template DNA for the FishATL_Cocktail2 (pool of three forward and three reverse primers); and 0.6 µM of each primer with 2.5 µL of template DNA for the 12S (miFISH-U_F/E_F and miFISH-U_R/E_R) primer set. The thermal cycling conditions for each set are detailed in Table 1. PCR-negative controls were included in all reactions to monitor potential contamination. The two PCR replicates for each sample and genetic region were combined before proceeding to a secondary PCR reaction, in which Nextera XT indexes and sequencing adapters were added to both ends of the amplified target region (Illumina, 2013). Finally, the PCR products underwent purification and normalization using the SequalPrep 96-well plate kit (ThermoFisher Scientific, Waltham, MA, USA) (Comeau et al., 2017), pooled, and subjected to paired-end sequencing on an Illumina MiSeq® sequencer using the MiSeq Reagent Kit v3 (600 cycles), following the manufacturer’s instructions (Illumina, San Diego, CA, USA) at Genoinseq (Cantanhede, Portugal). The raw sequence data generated from bulk ichthyoplankton samples are available in the NCBI Sequence Read Archive (SRA) under the BioProject accession number PRJNA1334324. Data corresponding to water eDNA samples will be publicly available upon subsequent publications.

### Bioinformatic processing

PMiFish version 2.4.1 (available at https://github.com/rogotoh/PMiFish.git; Miya et al., 2020) was used to process all raw FASTQ files obtained from high-throughput sequencing. This pipeline allows the analysis of data from multiple molecular markers and primer sets, enabling more standardized comparisons between the molecular techniques in the study. Based on USEARCH version 10.0.240 (Edgar, 2010), the pipeline performed several steps following a systematic workflow: (i) paired-end reads (Forward and Reverse) were merged, with reads containing more than five mismatches within the aligned region or a length shorter than 100 bp excluded, during an initial quality filtering task; (ii) primer sequences were then removed from the reads, and (iii) the reads were subjected to an additional quality filtering step to discard sequences shorter than the specified minimum length or exceeding a maximum expected error calculated from Phred quality scores; (iv) the remaining sequences were dereplicated to combine duplicates, and (v) a denoising step was performed to generate amplicon sequence variants (ASVs), eliminating chimeras and sequences containing errors; (vi) the ASVs were then taxonomically assigned to species with an identity threshold of >97%. In the end, molecular operational taxonomic units (MOTUs) were created, each represented by the ASV with the highest read count within its group of sequences. The COI reference database was built using the BAGS software (https://github.com/tadeu95/BAGS; Fontes et al., 2021) that mines sequences from BOLD (Ratnasingham & Hebert, 2007), while reference databases for the 12S and 16S markers were created by extracting available sequences from NCBI GenBank (Sayers et al., 2019). For the 12S and 16S datasets, all unverified records (those unconfirmed by GenBank staff for accuracy) and entries without species-level identifications were excluded. Additionally, all records were reviewed to correct synonyms and unaccepted names based on the accepted nomenclature from the World Register of Marine Species (WoRMS Editorial Board, 2024) in order to ensure taxonomic consistency across the entire dataset. For COI data, each sequence was assigned to a Barcode Index Number (BIN) (Ratnasingham & Hebert, 2013), which was also cross-checked against BOLD. All this curation process was carried out to improve the precision of DNA identifications and minimize potential misclassifications, while also reinforcing the reliability and accuracy of the reference databases utilized in the study. A minimum number of reads criterion was not applied to ensure more reliable comparisons, given the typically lower read counts observed in eDNA samples.

### Data analysis

Venn diagrams were generated using the InteractiVenn web tool (https://interactivenn.net/; Heberle et al., 2015) to evaluate the overlap and exclusivity of species detections between morphology and DNA metabarcoding-based identifications, the different sampling methods and among molecular markers and primer sets within the DNA-based datasets. The resulting diagrams were refined and edited using Inkscape 1.2 (Inkscape Project, 2020). Bar plots representing the monthly variation in the total number of fish species identified using different sampling approaches and species intersections across seasons of the year were generated using the ggplot2 package (Wickham, 2016) and the UpsetR package (Conway et al., 2017) in RStudio version 4.0.4 (RStudio Team, 2020). Seasonal groups were formed by assigning months based on where most of their days fell within a specific season. Furthermore, fish species recorded in the datasets were categorized based on the number of months in which they were detected, allowing for the identification of temporal patterns for each taxon. To achieve this, species were grouped into six categories: i) Very Frequent - detected in 12-13 months; ii) Frequent - detected in 8-11 months; iii) Moderately Frequent - detected in 4-7 months; iv) Rare - detected in 2-3 months; and v) Very Rare - detected in a single month. This classification enabled a clearer understanding of temporal variation and seasonal trends within the fish community, and the identification of specific spawning periods, when applied only to ichthyoplankton data. The Jaccard dissimilarity index was calculated among each sampling month to evaluate differences based on primer sets, seasons, and sampling methods using the "vegdist" function from the vegan R package (v2.6 -4) (Oksanen et al., 2022). Significant differences across these variables were assessed through Permutational Multivariate Analysis of Variance (PERMANOVA) using the "adonis2" function in the same package with 999 permutations. To visualize these patterns, a principal coordinate analysis (PCoA) was performed using R scripts with the vegan R package, highlighting trends in species recovery across seasons.

## Results

### Morphology-based identification of ichthyoplankton

Over the 13-month sampling period, 12,675 ichthyoplankton specimens were counted, consisting of 11,431 eggs and 1,244 larvae. The highest abundance of eggs was observed during April* (n = 6,831) and May was the month with the largest number of larvae registered (n = 506) (Table 2). In contrast, November recorded the lowest ichthyoplankton counts, with no eggs identified (like December) and only seven larvae captured.

**Table 2.** Monthly egg and larval counts, including taxonomic identification of fish larvae at the respective taxonomic levels. (*) - Refers to the sample collected in April of the subsequent year (2023).

| Month | Fish Eggs |  |  |  | Fish Larvae |  |  |  |
| --- | --- | --- | --- | --- | --- | --- | --- | --- |
|  | Total | Sardines | Anchovies | Other Sp. | Total | Species | Genus | Family |
| **April** | 862 | 244 | 20 | 598 | 32 | 8 | 1 | 2 |
| **May** | 527 | 354 | 6 | 167 | 506 | 12 | 3 | 3 |
| **June** | 922 | 687 | 4 | 231 | 60 | 8 | 2 | 1 |
| **July** | 346 | 272 | 18 | 56 | 94 | 10 | 3 | 1 |
| **August** | 159 | 110 | 2 | 47 | 106 | 9 | 3 | 2 |
| **September** | 18 | 11 | 4 | 3 | 12 | 4 | 0 | 2 |
| **October** | 75 | 46 | 0 | 29 | 19 | 6 | 0 | 1 |
| **November** | 0 | 0 | 0 | 0 | 7 | 5 | 0 | 0 |
| **December** | 0 | 0 | 0 | 0 | 12 | 4 | 0 | 0 |
| **January** | 16 | 6 | 0 | 10 | 8 | 3 | 1 | 0 |
| **February** | 497 | 297 | 0 | 200 | 228 | 9 | 1 | 2 |
| **March** | 1178 | 62 | 0 | 1116 | 72 | 5 | 0 | 1 |
| **April\*** | 6831 | 444 | 400 | 5987 | 88 | 6 | 2 | 1 |

The majority of the eggs (73.87%; n = 8,444) could not be identified due to the lack of distinctive morphological traits, which limited identifications to only two species, with sardine eggs (*Sardina pilchardus*) accounting for 22.16% of the total egg count, while anchovy eggs (*Engraulis encrasicolus*) represented 3.97%. Sardine egg counts outnumbered those of anchovy in all sampled months.

Looking at the larvae data, of the 1,244 individuals counted, 68% were identified to species level, 8.04% to genus level, and 20.10% to family level. Important to note that only 48 specimens (3.86%) could not be taxonomically identified, showing the overall success of larval morphological identification. In total, 23 fish species were identified, with *Sardina pilchardus* leading the records with 348 individuals counted (28% of the total larvae), followed by *Pomatoschistus pictus* (n = 115) and *Parablennius pilicornis* (n = 112). Additionally, the second highest count was larvae from the Sparidae family with 233 individuals identified (Table S.2).

### DNA metabarcoding identification of ichthyoplankton

A total of 12,712,713 reads were generated from the Illumina MiSeq high-throughput sequencing of 156 datasets (3 replicates × 4 primer sets × 13 months, plus negative controls). Following bioinformatic processing - including primer removal, demultiplexing, quality filtering, and denoising - 11,548,681 reads were retained, representing 90.84% of the initial reads (Table S.3) After taxonomic assignment, the COI primer pair mlCOIintF/LoboR1 recovered 1,799,388 reads assigned to species, accounting for 46.05% of its usable reads (Table 3). Regarding fish species assignments, both mitochondrial ribosomal primer sets performed well: the 16S set showed the highest performance, with 44.09% of usable reads (1,230,830) assigned to fish, followed closely by the 12S set with 40.07%. Within the COI markers, FishATL_Cocktail2 yielded more fish-specific assignments than mlCOIintF/LoboR1 (557,197 vs. 195,755 reads), despite lower overall read retention and total species-level assignments, underscoring its greater specificity for fish. Across all primer sets and datasets from the entire study, a total of 115 fish species were identified with DNA metabarcoding (Figure 2; Table S.4-S.7). Among the primer sets, the 12S miFISH U-E demonstrated the highest performance, detecting 80 species, followed closely by the Fish 16S primer set, which identified 78 species. In comparison, the COI primer sets recovered fewer fish species, with FishATL_Cocktail2 outperforming mlCOIintF/LoboR1 by detecting 62 versus 51 species.

**Figure 2.**
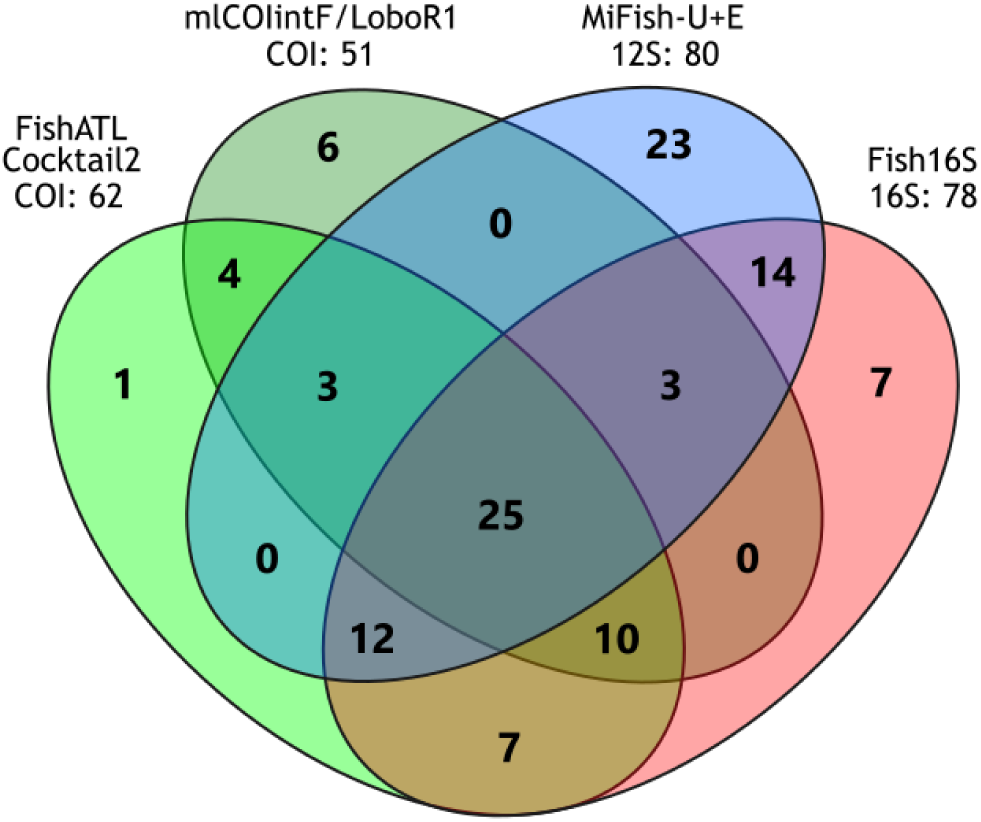
Partitioning of the total fish species identified by each primer pair/set throughout the study. The numbers next to each primer indicate the total number of fish species detected by each primer combination.

**Table 3.**
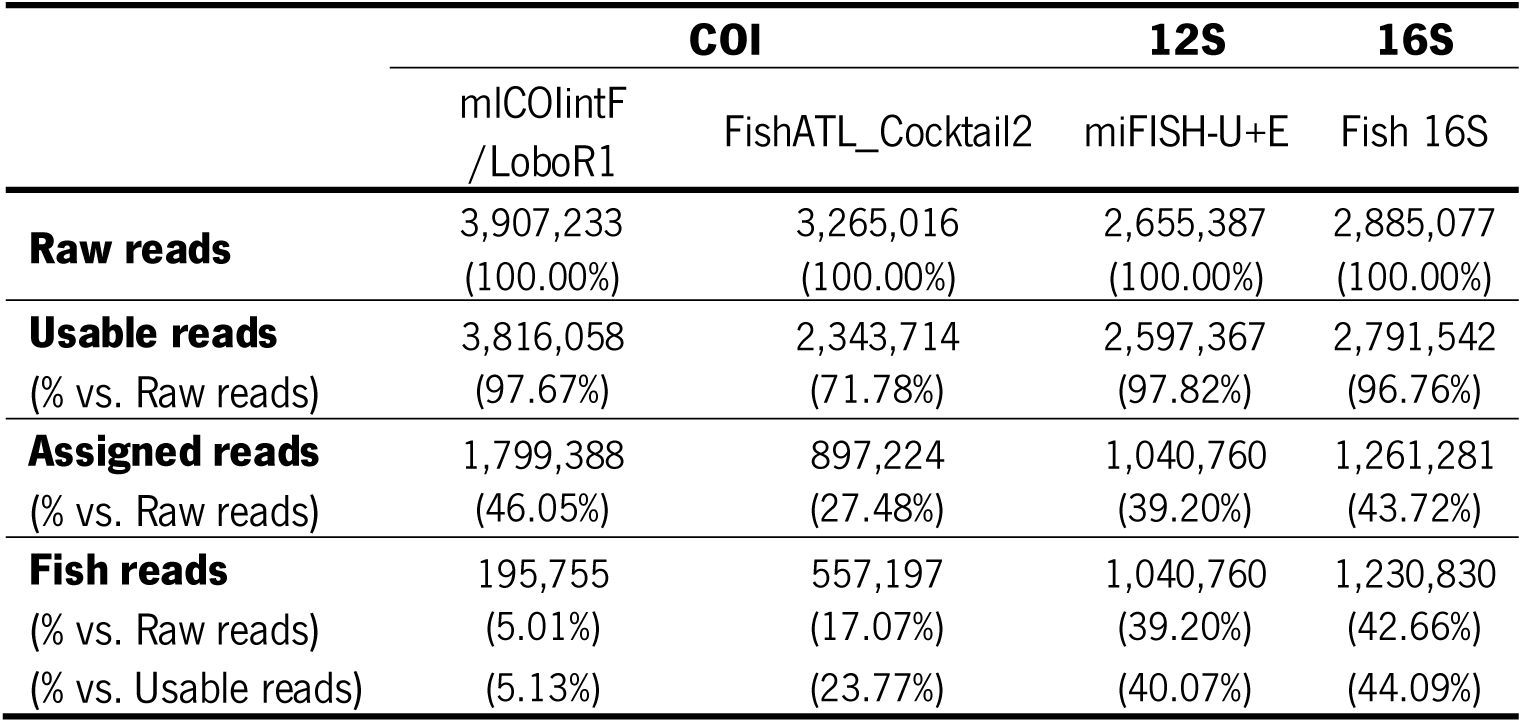
Total number of Raw reads generated through Illumina MiSeq high-throughput sequencing for DNA metabarcoding, Usable reads retained after bioinformatics processing steps (including primer removal, demultiplexing, quality filtering, and denoising), and reads taxonomically assigned to the species level (Assigned reads), and reads specifically assigned to fish species (Fish reads) for each primer set.

Together, the primer sets detected 25 fish species in common, accounting for 21.74% of the total fish diversity recovered by these molecular approaches. Notably, 34.78% of the species were detected by all three genetic markers, considering at least one of the COI primer sets in addition to the 12S and 16S primers. Regarding exclusive detections, the 12S miFISH U-E primer set exhibited a superior ability to identify unique species (n = 23), followed by Fish16S with 7 unique detections, mlCOIintF/LoboR1 with 6, and FishATL_Cocktail2 with only 1 unique species detected. The data obtained from the different primer sets was combined for comparative analysis in the following section.

### Morphology versus DNA metabarcoding: species-level identifications of ichthyoplankton

The comparison of species-level identifications between the molecular DNA metabarcoding approach and the traditional morphological identification method revealed a substantial disparity in species detected (115 *versus* 23, respectively). Over the thirteen months of sampling, 468 fish species records were obtained, reflecting the cumulative occurrences across this period. From these records, 378 were made exclusively through DNA metabarcoding, 33 via morphological identification, and 57 by both approaches.

DNA metabarcoding successfully identified 96 fish species that were not detected by the traditional morphological methods (Figure 3), highlighting its ability to retrieve hidden diversity within the multiplicity of fish eggs and larvae. In total, 19 species were identified by both approaches, while morphology exclusively detected four species (*Diplecogaster bimaculata*, *Scophthalmus rhombus*, *Solea solea*, and *Syngnathus typhle*). All the information generated from the ichthyoplankton samples, including DNA metabarcoding and morphological data, was combined to assess the efficiency of distinct sampling methods in detecting fish species in the next section.

**Figure 3.**
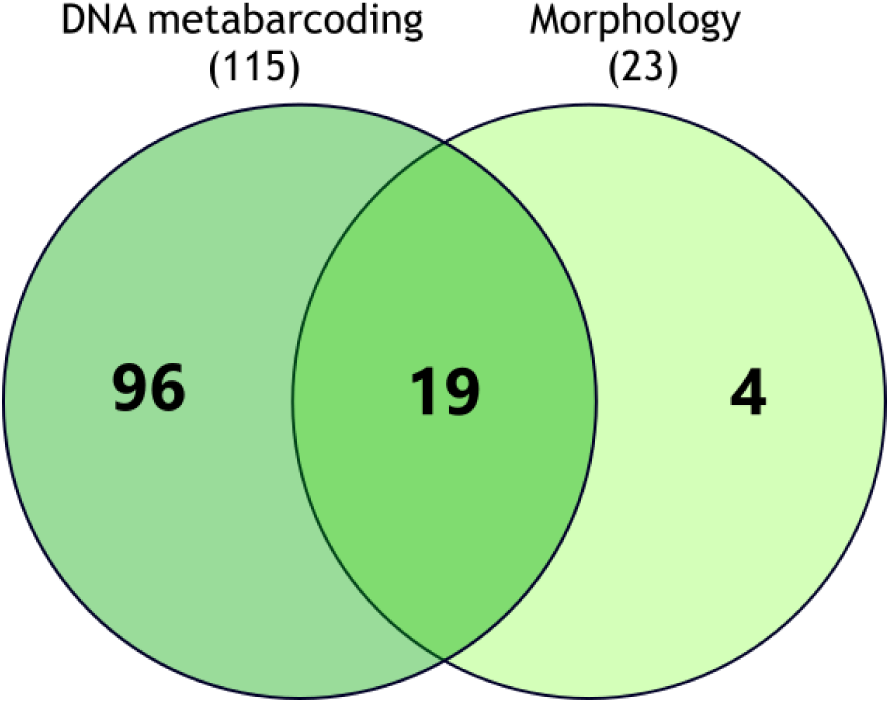
Partitioning of the total fish species identified by both methodologies - DNA metabarcoding and morphology. The numbers in parentheses represent the total number of fish species detected by each approach.

### Fish spawning insights based on ichthyoplankton data

The intersection plot (Figure 4) provides a detailed visualization of fish species detected in each season based on ichthyoplankton samples, with the objective of illustrate the importance of multi-seasonal sampling to obtain information about fish-specific spawning periods. Six Elasmobranchii species were identified by DNA but excluded from this analysis, as no encapsulated eggs were collected in our samples and their presence likely originated from external environmental DNA. From the 113 taxa identified, twenty-five species (22.12% of the fish diversity) were detected across all four seasons, indicating their broad temporal distribution and continuous reproductive activity. On the other hand, it was evident that several species have restricted reproductive periods, being found exclusively in a single season (22 species in spring, 9 species in winter, 7 species in summer, and 6 species in autumn). Species present in two seasons represented 24.78% of the total diversity, with the combination spring-summer standing out with 12 shared species during these warmer months, whereas species occurring across three seasons were less frequent (14.16% of the detections). Remarkably, the top three intersections (Figure 4) involved the spring season, the period marking the onset of optimal spawning conditions, reinforcing its dominant role to overall species richness in the ichthyoplankton samples.

**Figure 4.**
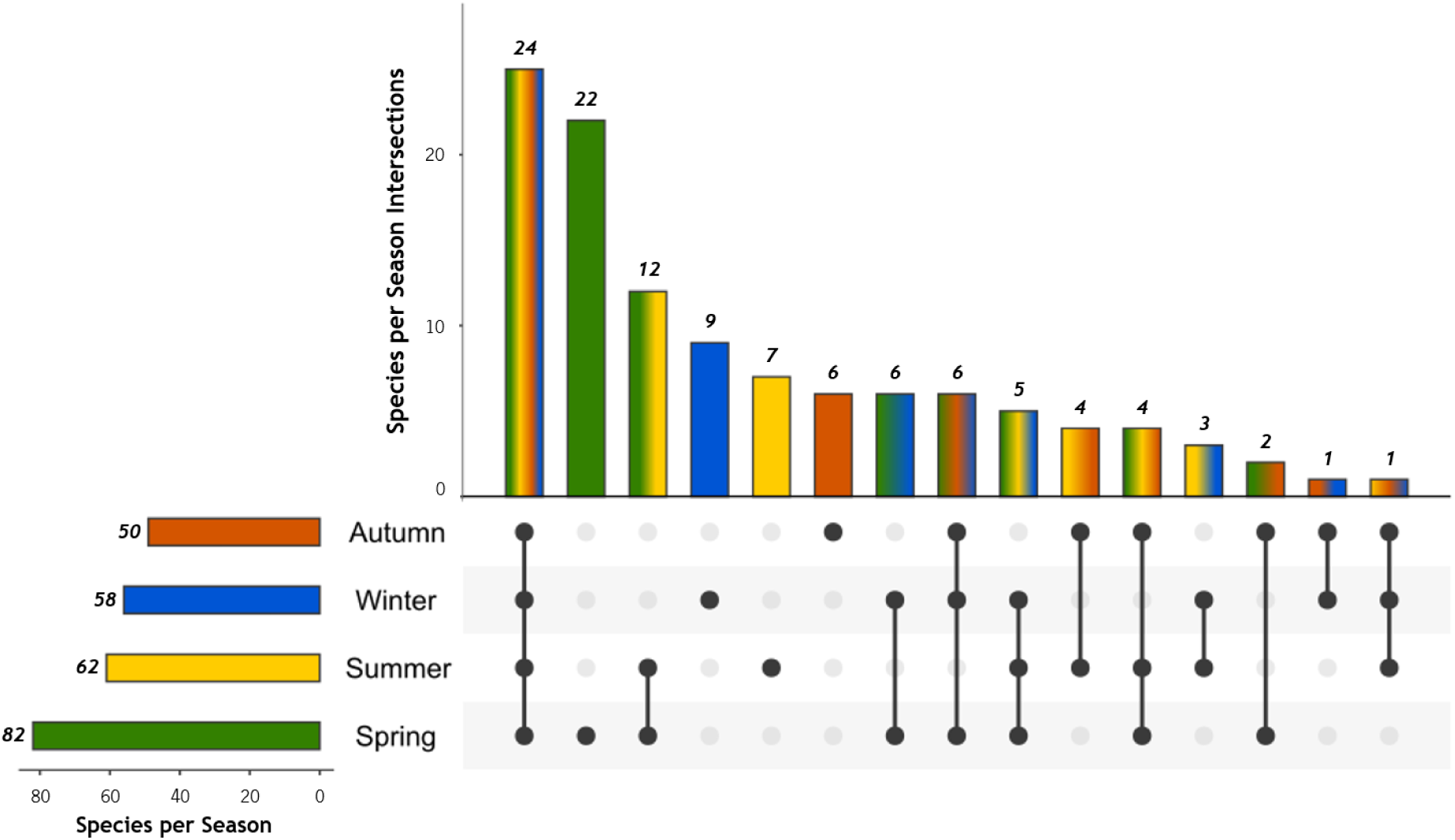
Species intersections across seasons of the year based on ichthyoplankton data. The bar plot shows the number of fish species identified exclusively or shared among seasons, with colors representing individual seasons: Autumn (orange), Winter (blue), Summer (yellow), and Spring (green). The connected dots below the bars indicate the specific seasonal combinations for shared species. The number of species detected in each season is displayed on the left as horizontal bars.

### Ichthyoplankton versus eDNA: fish species-level resolution

Different sampling methods were compared to evaluate their effectiveness in fish species identification using metabarcoding techniques: ichthyoplankton samples (combining DNA metabarcoding with additional data from morphological identification) and water samples (using eDNA metabarcoding). The eDNA data was processed and analyzed as part of a parallel study (in prep.) and subsequently integrated into this comparison analysis. By combining the data obtained from both approaches, a total of 131 fish species were identified throughout the sampling period (Table S.8). A greater fish diversity was identified through ichthyoplankton samples (119 species) compared to water eDNA samples (72 species). While both sampling types shared 60 species in common, ichthyoplankton samples yielded 59 exclusive species, whereas eDNA samples recovered only 12 unique species (Figure 5). This initial trend extended to higher taxonomic levels, with ichthyoplankton samples outperforming eDNA in the number of identified genera, families, and orders.

**Figure 5.**
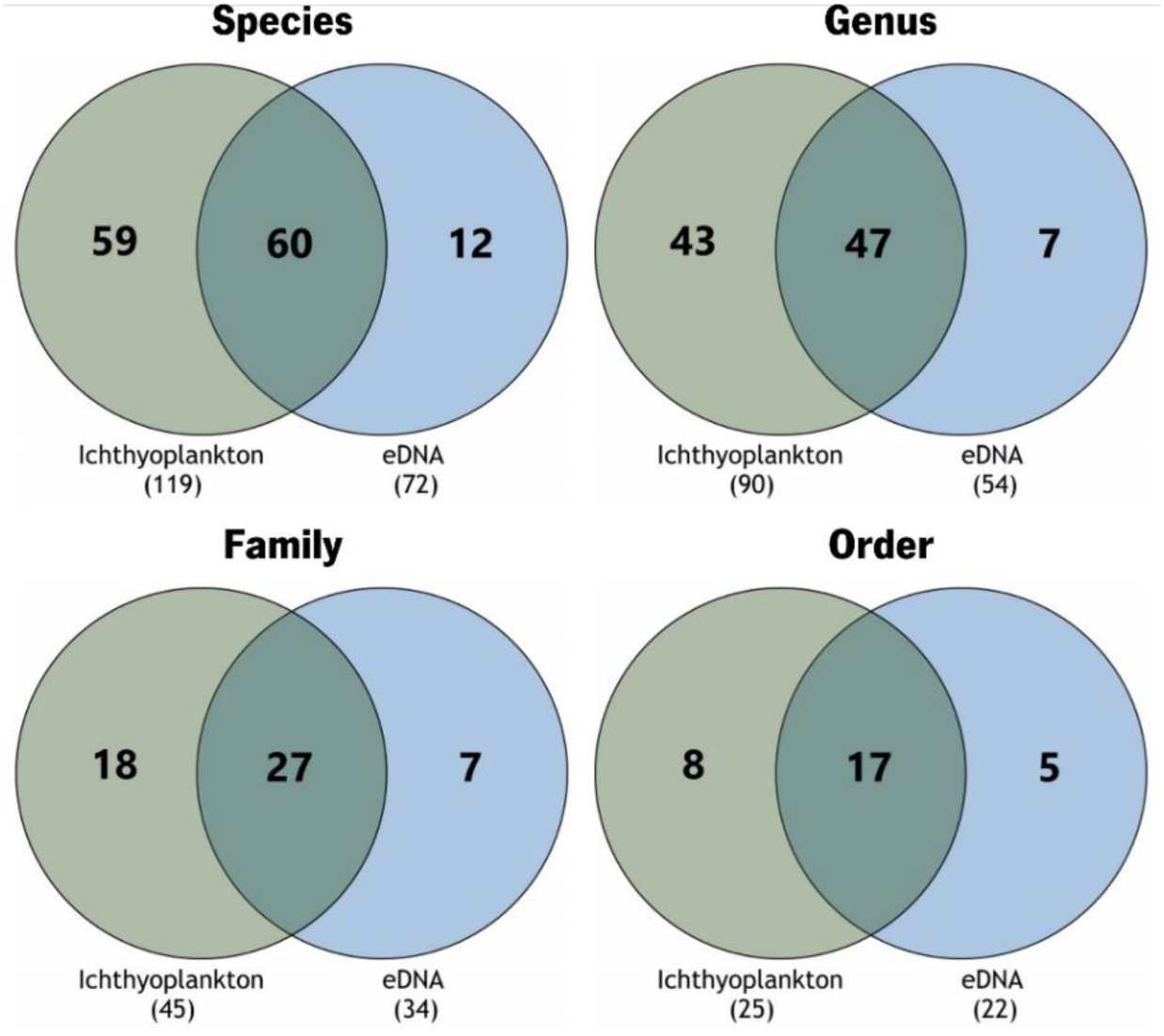
Partitioning of fish taxa detected by ichthyoplankton samples (DNA metabarcoding combined with morphology) and water samples (eDNA), displayed at the species, genus, family, and order taxonomic levels. The values within the parentheses indicate the total number of taxa identified by each method in the corresponding taxonomic category.

Based on the categorization of fish species by the number of monthly occurrences (Table S.9), 12 species - *Atherina presbyter, Engraulis encrasicolus, Sardina pilchardus, Dicentrarchus labrax, Diplodus bellottii, Diplodus sargus, Sparus aurata, Pomatoschistus microps, Chelon labrosus, Mugil cephalus, Solea senegalensis, and Syngnathus rostellatus* - were identified as “very frequent” in the Guadiana Estuary (9.16% of the total fish diversity). These species were detected in 12 or 13 months of the sampling period, indicating year-round presence. Seventeen species were classified as “frequent,” appearing in 8 to 11 months of the year, and 26 species were categorized as “moderately frequent” with detections spanning 4 to 7 months. In the “rare” category, encompassing detections in 2 to 3 months, 28 species were registered. Among these, two exceptional cases (*Chelon auratus* and *Gobius couchi*) had records spread across three different seasons, with one record in each season. Seventeen species were identified across two seasons, with spring-summer combination accounting for 47.06% of these cases. The remaining nine species were recorded only in a single season: four in spring, four in summer, and one in autumn. Lastly, the “very rare” category included 48 species (36.64% of the total fish species), each detected in only one month of the sampling period. Of these, 26 records occurred in spring, 5 in summer, 6 in autumn, and 11 in winter.

Throughout the 13-month sampling period, a total of 560 fish species identifications were recorded. Most of these identifications were obtained exclusively through ichthyoplankton samples, accounting for 303 records (54.11%). This was followed by 166 identifications (29.64%) that were shared between both ichthyoplankton and water eDNA, while water eDNA alone contributed to 91 unique records (16.25%). The fish diversity retrieved by both methods demonstrated a fluctuating detection capacity for specific species across sampling months, as illustrated in Figure 6 for the nine species consistently identified throughout all 13 months of sampling. For instance, the first listed species *Atherina presbyter*, was identified by both ichthyoplankton and eDNA in six months of the study. However, in six other months the species was detected exclusively through ichthyoplankton samples, while exclusive eDNA detection was made solely in one month. A similar pattern was observed for 28 additional species, representing 46.67% of the 60 species detected by both methods.

**Figure 6.**
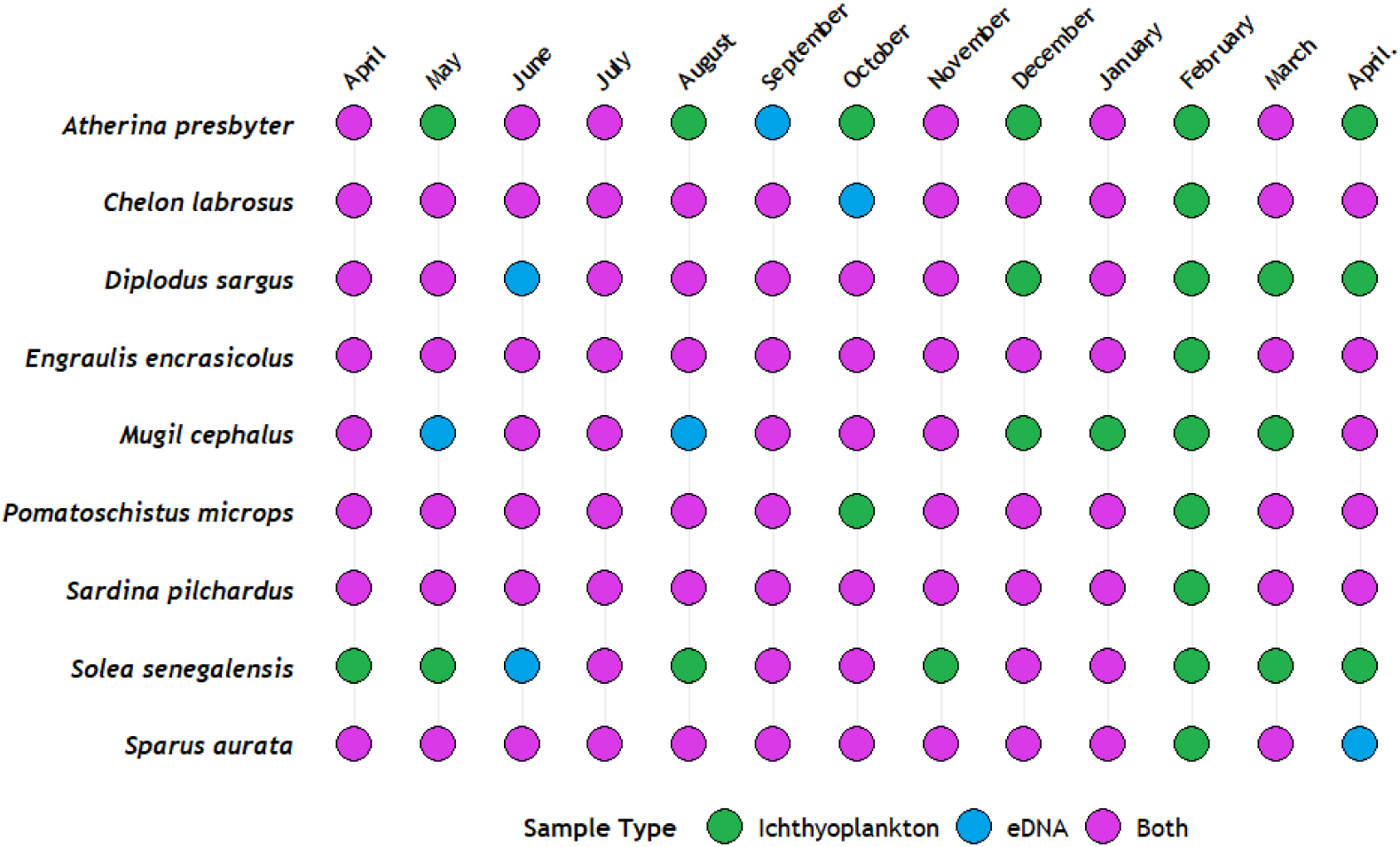
Detection of the nine fish species identified across all 13 months of sampling through bulk ichthyoplankton samples (green), water eDNA (blue), or both methods (pink). The figure highlights the variable detection capacity of each sampling method and the inherent randomness in species detection.

### Seasonal variation in fish species detected by both sampling methods

The results presented in Figure 7 demonstrate both monthly and seasonal variation in terms of fish species richness identified using ichthyoplankton samples and water eDNA. During the sampling period, species diversity peaked during spring, with April and May obtaining the highest number of species (64 and 60, respectively), followed by a slight decline in summer months. Autumn recorded the lowest overall diversity (with October registering the minimum number of species identified, 29), while the winter months obtained a modest recovery in species detections. When considering sample type performance, ichthyoplankton samples consistently surpassed water eDNA in fish identifications throughout the study period, except for the initial month of April.

**Figure 7.**
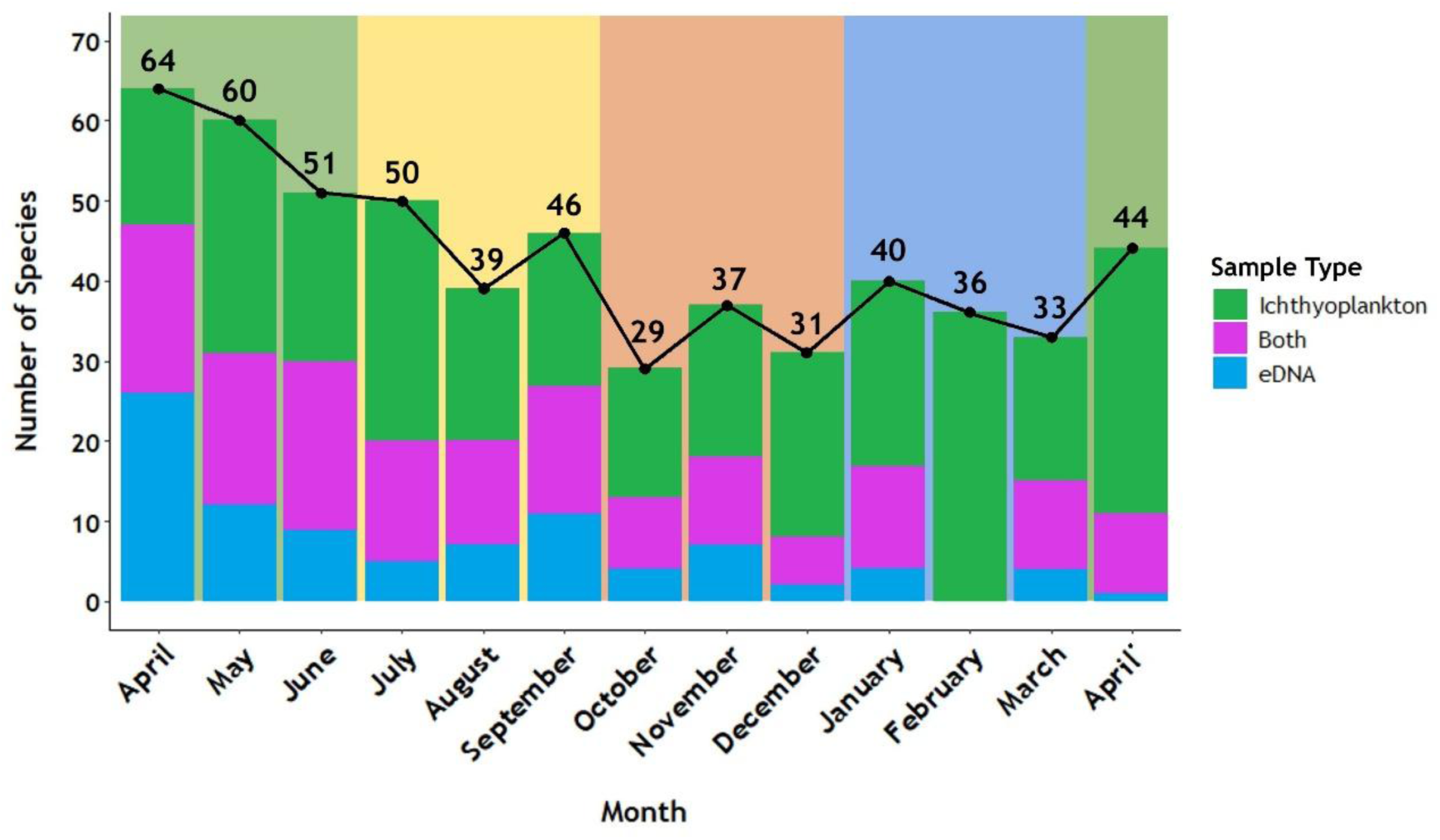
Monthly variation in the total number of fish species identified using different sampling approaches: ichthyoplankton samples (green), water samples (blue), and species detected by both methods (purple). The black line represents the total number of species identified each month, with the corresponding values displayed above each bar. Seasonal periods are indicated by the shaded background: Spring (green), Summer (yellow), Autumn (orange), and Winter (blue). The solid green bar in February reflects the sequencing failure of eDNA samples for that month.

The Venn diagram (Figure 8) presents the seasonal partitioning of fish species identified by combining data from both sampling methods (ichthyoplankton and water eDNA). The previous pattern was confirmed once again, with spring emerging as the most diverse season, showing the highest number of species detected (n = 99), followed by summer (71 species) and winter (61 species) (Table S.10). Autumn was the season that recorded the lowest number of species (n = 56). Following this tendency, warmer months (spring and summer) were characterized by higher species richness, with a total of 112 species identified, of which 53 species (40.46% of the total diversity) were exclusive to this period. In contrast, the colder months (autumn and winter) retrieved 78 species, but only 19 species (14.50%) were exclusively detected in these seasons.

**Figure 8.**
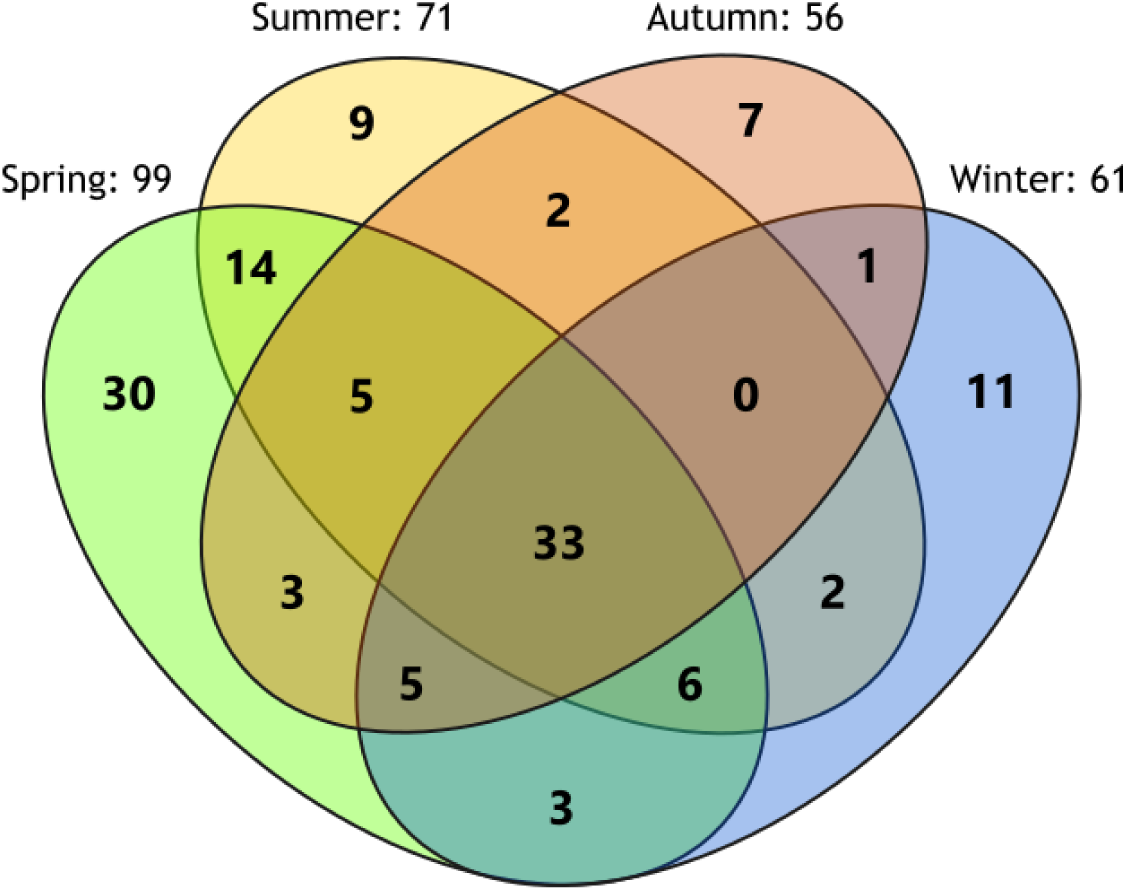
Partitioning of the total fish species identified across seasons of the year. The total number of species detected in each season is indicated next to its respective section.

Seasonal differences in the fish community structure were also evident by the Principal Coordinates Analysis (PCoA; Figure 9). The clustering patterns observed for each season suggest seasonal differentiation in the local fish community, a finding statistically supported by the PERMANOVA analysis, which confirmed significant differences in fish composition across seasons (pseudo-F = 1.6724, df = 3, p = 0.004; Table S.11). Spring samples (green) exhibited higher dispersion, reflecting greater variability within the seasonal fish community. This variability may be influenced by the additional sampling month from the subsequent year, suggesting temporal shifts in taxa composition between years. In turn, summer samples (yellow) were more closely clustered, probably indicating a more homogeneous fish composition during this season. The distinct groups formed independently by autumn (orange) and winter samples (blue) illustrate a clear shift in community structure compared to spring and summer, highlighting the unique species composition associated to these colder months.

**Figure 9.**
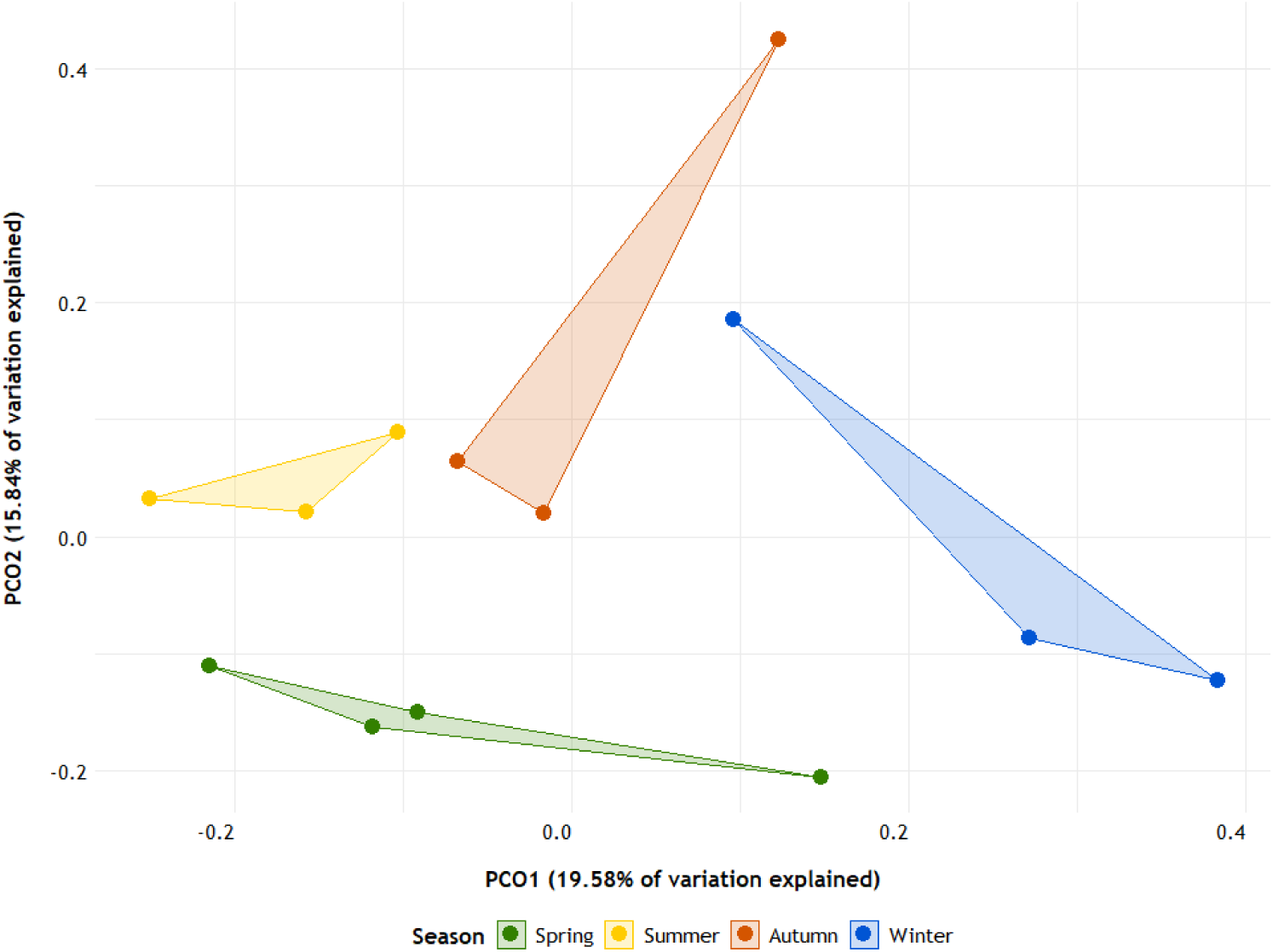
Principal Coordinates Analysis (PCoA) plot based on the Jaccard dissimilarity index, illustrating seasonal variation in fish taxa composition. Each point represents a sampling month, grouped by season (Spring: green, Summer: yellow, Autumn: orange, Winter: blue). Clusters indicate community similarity within seasons, while greater distances reflect increased dissimilarity between seasons.

## Discussion

This study aimed to employ DNA-based approaches allied with morphology to investigate fish community dynamics over an annual cycle in the Guadiana Estuary. Distinct fish communities were successfully identified across seasons, with statistical analyses confirming significant differences between them. The warmer seasons (spring and summer) obtained higher species richness in the estuary, corresponding to the main reproductive period of fish. Spring was the peak reproductive period, with 82 different species identified in ichthyoplankton samples, reflecting the optimal conditions for fish reproduction. In comparison to spring, total fish diversity showed a marked decrease during the remaining seasons, with declines ranging from 21.37% in summer to 32.82% in autumn. The comparison of sampling methods for fish detection revealed that ichthyoplankton samples, analyzed through DNA metabarcoding and morphological inspection, detected substantially more species than water eDNA samples. Environmental DNA unexpectedly detected 47 fewer fish species than bulk ichthyoplankton samples, which represents a reduction of 35.88% in the total diversity recovered.

### Detection of fish species in ichthyoplankton through DNA metabarcoding versus morphology

Globally, DNA metabarcoding demonstrated a higher capability to identify fish to the species level compared to morphology in ichthyoplankton samples. Across the 13-month study period, 115 species were identified through metabarcoding, in contrast to the 23 species identified by morphology, underscoring the superior taxonomic resolution and diversity detection offered by DNA metabarcoding (Ferreira et al., 2024). The molecular approach successfully identified all but four species across all sampling months, accounting for most species detections. Of these, 80.77% were exclusive to metabarcoding, while an additional 12.18% were identified through both metabarcoding and morphological inspection. This better performance can be associated with the use of four primer sets targeting three distinct molecular markers, which significantly enhanced the diversity recovered. This multi-marker strategy increased species detection from 30.43% to 55.46% compared to using a single primer set, aligning with similar patterns of increased detection reported in other studies (Duke & Burton, 2020; Ferreira et al., 2024; Jiang et al., 2022; Teixeira et al., 2023). Moreover, the likelihood of false positives resulting from laboratory contamination is minimal, providing greater confidence in our taxonomic assignments, as all negative controls consistently displayed no amplification throughout the study.

Morphological identification, on the other hand, faced important limitations. Among the larvae analyzed, 28.14% of the identifications were made at higher taxonomic ranks, and an additional 3.86% of larvae identifications could not be reached at all. The challenge was even more pronounced for fish eggs, with 73.87% remaining unidentified due to the absence of distinctive morphological traits. These findings expose the inherent difficulties of morphological identification, which persist even for experienced taxonomists (Ko et al., 2013; Nobile et al., 2019). Identifying such small and morphologically similar organisms is particularly challenging, being intensified by damaged specimens and the lack of comprehensive identification keys for early life stages (Hulley et al., 2018; Reynalte-Tataje et al., 2012). Our results confirm that DNA metabarcoding can be a complementary tool for addressing these challenges, revealing "hidden diversity" often overlooked by morphological methods. The molecular approach not only enhances the ability to identify species, but also provides a valuable resource for taxonomists, enabling a possible reidentification of specimens with molecular information serving as a reliable starting point (Ferreira et al., 2024).

Comparing both identification approaches also revealed the impossibility of establishing a clear connection between the number of eggs and larvae counted in a sample and the number of reads generated through metabarcoding - using *Sardina pilchardus* and *Engraulis encrasicolus* data as a case study (Table S.12). A striking randomness in read generation was observed; for instance, in months with a higher count of sardine eggs and larvae compared to anchovies, the metabarcoding reads were paradoxically higher for *Engraulis*. Similarly, within the anchovy’s data, months with higher morphological counts yielded fewer reads than months with lower counts. There were even cases where no morphologically identified eggs or larvae were recorded, yet a substantial number of reads were attributed to species not identified through morphology. This discrepancy can be attributed to the fact that the samples used for morphology and metabarcoding analyses were obtained from consecutive trawls, leading to variations in sample composition. Additionally, the existence of eggs and larvae in our samples that could not be identified to species level by morphology, probably contributed to some of these reads, as well as the potential attachment of external DNA belonging to species not physically present in the samples (Taberlet et al., 2012). These findings expose the major limitation of DNA metabarcoding, namely its inability to provide accurate quantitative data for individual species (Carvalho, 2022; Derycke et al., 2021). This challenge needs to be addressed in the near future to expand the utility of metabarcoding methods for ecological studies, with special focus on mitigating issues such as primer and amplification bias, sequencing mismatches, and improving our understanding about different DNA release rates among species (Elbrecht & Leese, 2015; Leray & Knowlton, 2017).

### Detection efficiency of fish species: bulk ichthyoplankton metabarcoding versus eDNA

Bulk ichthyoplankton metabarcoding and eDNA have different biological targets and provide distinct ecological information on fish assemblages. The first is expected to indicate the presence of early life stages (i.e., eggs and larvae), thereby serving as a proxy for fish species reproduction, whereas eDNA should, in principle, reflect the presence of fish species at any life stage, from egg to adult. Henceforth, eDNA would be expected to detect not only the same species detected in ichthyoplankton samples, but also additional ones that were present in the same location only as juveniles or adults. Contrary to these expectations, our results revealed that bulk ichthyoplankton metabarcoding identified considerably more fish species than water eDNA, which globally exhibited about a 40% lower success in species recovery and lower taxonomic diversity. These results were consistent throughout the sampled months, and the confirmation of bulk ichthyoplankton detections, at least partially, through morphology, reinforces these findings. Specifically, 119 species were identified in ichthyoplankton using both morphology and DNA metabarcoding, compared to 72 species detected via water eDNA. In terms of total records, ichthyoplankton samples accounted for 83.57% of identifications, while eDNA contributed to 45.89% (including mutual and exclusive detections). The disparity might have been slightly mitigated if the eDNA data from February had not experienced amplification issues, however, the overall conclusion remains unaffected.

Because the primers and bioinformatic pipeline were the same for bulk ichthyoplankton and eDNA, these results can possibly be explained by the impact of the sampling approach (plankton tow vs filtered water) and targeted biological material (bulk organisms vs extra-organismal DNA) on the detection of fish species in the same sampling location (see Peterson et al., 2022; Pompeu et al., 2023). In this context, several sample-related factors may be at play. Ichthyoplankton samples collected via net sampling provide higher-quality and more concentrated DNA, offering a robust template for amplification (van der Loos & Nijland, 2021) and a consistent reflection of reproductive activity in the estuary (Pritt et al., 2015). In contrast, the comparatively lower eDNA-based detection may be attributed to the smaller sample size, with only 12 L of water processed per month (two 2 L samples at three points), compared to the substantially larger volumes of water sampled during plankton tows, where eggs and larvae provide higher amounts of DNA for amplification. Furthermore, the rapid degradation and fragmentation of eDNA in seawater (Thomsen et al., 2012) likely contributed to the reduced detection success compared to bulk samples, highlighting the heightened sensitivity of eDNA to low DNA concentrations in the system (Harrison et al., 2019). Another important factor to consider is the comparatively lower amount of DNA shed from eggs and larvae compared to fully developed stages, since it has been shown that the amount of DNA shed into the water is more related to adult fish biomass (Jo et al., 2019; Maruyama et al., 2014). It is also likely that species uniquely detected by eDNA were non-reproducing individuals, such as juveniles or adults, whose genetic material was dispersed in the water. Finally, eventual differences in detection success may have originated from the DNA extraction procedures, which were tailored to each sample type, as well as from the susceptibility of eDNA to contamination, leading to heterogeneity in PCR amplification, primer biases, and failure to detect rare species (Deiner et al., 2017).

Rather than looking at these differences simply as limitations, they can be seen as indications of the complementary nature of both approaches. The ability of both sampling methods to retrieve distinct components of fish diversity is particularly relevant given the fluctuating detection of certain species across months (Figure 6). Some species detected by both methods in one month were often identified by only one approach in another, pointing out the importance of using two distinct sampling strategies in order to maximize the diversity recovered and account for this variability in the detection process. It also reinforces that neither method alone can obtain a complete representation of fish community composition in estuarine systems, so researchers need to carefully consider the implications of sampling methods and targeted biological material on taxon detection through metabarcoding. This is very important even when employing essentially the same molecular workflow, from primer selection to bioinformatics processing, highlighting the need for continuous evaluation and refinement of eDNA methodologies to enhance their reliability (Jackman et al., 2021) and to address current limitations in species identification within estuarine environments.

### Seasonal variation in fish community composition and species richness

One of the main goals of this study was to uncover seasonal variations in the local fish community using molecular techniques. The (e)DNA metabarcoding results revealed that this objective was achieved with distinct fish communities observed across the sampled months and seasons (Figure 7-9). Variations in fish community composition and the seasonal exclusivity of certain species highlight the complex ecological processes occurring in the Guadiana Estuary, including spawning cycles, environmental conditions, and habitat availability, which collectively influence fish diversity and community composition throughout the year (Faria et al., 2006; Muhling et al., 2008).

Spring emerged as the season with the highest fish diversity (n = 99) and the best reproductive period for numerous fish species and subsequent developmental stages (Kindong et al., 2020; Monteiro et al., 2021), with 84 of these species being identified in ichthyoplankton samples. To this scenario contribute several favorable environmental conditions, such as increasing water temperatures, nutrient availability, and primary productivity, that create an optimal environment for spawning and the survival of early life stages (Muhling et al., 2008; Rodriguez et al., 2009). Another factor that supports these conditions is the seasonal upwelling events that characterize the Portuguese coast, resulting in an input of nutrients and food essential for the development of ichthyoplankton and to support large ichthyofaunal communities (Santos et al., 2001; Santos et al., 2004; Wooster et al., 1976). Several species, such as *Sardina pilchardus* and *Engraulis encrasicolus*, benefit from these seasonal environmental cues by synchronizing their reproductive efforts with these periods to maximize the offspring access to abundant food resources, which enhancing their chances of survival (Katara et al., 2011; Morais, 2007; Santos et al., 2001). Summer stable conditions created a great nursery environment for early life stages and juvenile growth, supporting a diverse fish community with 71 species identified. The cooler temperatures associated with autumn and winter directly impact the reproductive activity and food availability of regional fish species, leading to the lower diversities found in these months. Notably, multiple species identified during this study originate from adjacent coastal areas where they reproduce, with the eggs and larvae drifting into the estuary by natural processes (Churchill et al., 1999), mainly influenced by tides in the lower estuary through the mechanism of "selective tidal stream transport" (Faria et al., 2006; Primo et al., 2012). This pattern aligns with historical findings for Portuguese estuaries, where most fish eggs and larvae result from coastal spawning events, with only a limited number of species actively seeking estuaries as ideal spawning grounds (Ré, 1999).

Finally, is important to note the differences observed in species composition across the two sampled April months, with only 32 species found in common out of a total of 76 (42.11%). This provided preliminary data on temporal shifts in regional taxonomic composition, likely associated with changes in environmental factors such as temperature, salinity, and nutrient availability during spawning seasons on an interannual basis (Gao et al., 2018; Sloterdijk et al., 2017). These findings reinforce the need for long-term monitoring in the Guadiana River Estuary to better understand local ecological processes and to develop conservation strategies that can sustain biodiversity and ecosystem resilience.

### Comparison of historical ichthyoplankton records in the Guadiana Estuary with DNA metabarcoding from bulk samples

The categories created based on monthly occurrences, when applied exclusively to the ichthyoplankton data, revealed valuable insights into species’ reproductive patterns across seasons. *Engraulis encrasicolus*, *Pomatoschistus microps*, and *Sardina pilchardus* were detected throughout the entire year, indicating continuous reproductive activity across all season and aligning with previous reports (Faria et al., 2006; Morais, 2007; Ré, 1999) - a trend also observed for five additional species that were detected in 12 out of 13 months. Fourteen species categorized as “frequent” indicate an extensive spawning period, while 27 “moderately frequent” species likely display reproductive activity during 4 to 7 months. The remaining ichthyofauna were detected over shorter timescales: twenty-nine species were categorized as “rare” appearing in 2 to 3 months of sampling, while 41 species were found in ichthyoplankton samples for only a single month. This limited occurrences are naturally associated with the restrict spawning period of several species, but can reflect sampling limitations, where individuals were not captured in other months despite potentially being present in the environment. To overcome this limiting factor, extending monthly monitoring across consecutive seasons can help validate these findings with additional annual records, support the patterns observed in this study, and potentially uncover new spawning periods for the local ichthyofauna (Monteiro et al., 2021; Zhang et al., 2022). Additionally, the determination of accurate spawning periods for each fish species can be conditioned by the different developmental stages of the collected larvae, which may have been spawned up to a month before collection, making fish eggs more precise indicators of recent spawning periods (Burghart et al., 2014; Lewis et al., 2016). Nevertheless, this potential temporal bias in spawning periods was minimized in our study, as most analysis were focused on seasonal data.

Regarding historical ichthyoplankton records in the Guadiana Estuary, previous studies have documented the presence of 53 fish species (Table S.13), identified as either eggs or larvae (Chícharo & Teodósio, 1991; Chícharo et al., 2006; Faria et al., 2006). By combining DNA metabarcoding with morphological analyses we were able to identify 35 of these species (66.04% of the historically recorded taxa), demonstrating the effectiveness of our approach to recover the ichthyofaunal diversity of the region. Focusing only on the sampling area of this study, the lower section of the estuary and the adjacent coastal zone, 31 species have been previously recorded, and of these, only seven species were not detected in our samples: *Ammodytes tobianus*, *Halobatrachus didactylus*, *Monochirus hispidus*, *Pomatoschistus minutus*, *Syngnathus abaster*, Syngnathus *acus*, and *Trigla lyra*. We checked our databases for the different markers, and all seven species were represented, except for *Trigla lyra* in the 12S database. All these species were identified in previous studies at relatively low abundances (Chícharo & Teodósio, 1991; Chícharo et al., 2006; Faria et al., 2006), which could explain their absence in our assignments due to limited availability during the sampling period. Another factor may explain the cases of *Pomatoschistus minutus* and both *Syngnathus* species. The lack of identification could originate from potential primer and sequencing biases favoring genetically related species such as *Pomatoschistus microps* or *Pomatoschistus pictus*, and *Syngnathus typhle* or *Syngnathus rostellatus*, species detected in our samples. This overlap likely resulted in an inability to discriminate between congeneric species (Ferreira et al., 2024; Fontes et al., 2024).

Through our molecular approach, we successfully matched the historical seasonal distribution for 69.79% of the seasons for the 24 historically documented species found in our data from the lower section of the Guadiana Estuary. We failed to record a historical seasonal presence in only 8 cases (8.33%), while also contributing with 21 additional records (21.88%) for these resident species, providing new insights into their temporal distribution. Within our data, we also identified new ichthyoplankton records for the Guadiana Estuary, being six of these records (*Chromogobius zebratus, Decapterus macrosoma, Parapristipoma trilineatum, Pseudaphya ferreri, Sicyopterus eudentatus* and *Zebrus zebrus*) potential non-indigenous species (NIS) in Portuguese waters, since they are not listed in the recent compilation of the Portuguese ichthyofauna by Carneiro et al. (2019). These records are probably some of the first documented occurrences of these species in the area, so each case should be interpreted with caution and assessed individually. The origin of three of these species is the Mediterranean Sea, making it plausible that their presence in the sampled region results from migration from adjacent waters. This includes *Pseudaphya ferreri*, which was detected in five different months using the 12S and 16S markers and emerged as one of the top records in November. Although not listed by Carneiro et al. (2019), this species has been previously reported along the Portuguese coast (Rodrigues et al., 2016). *Zebrus zebrus*, found in July and August with the 16S marker and displaying a substantial number of reads; and *Chromogobius zebratus*, detected in December with two reads using COI, which was potentially recorded for the first time along the southern coast of Portugal by Ferreira et al. (2024). The other three species, originating from distant regions, require more careful evaluation to confirm their actual presence in our samples. Two of these species have congeneric species previously described in Portuguese waters: *Decapterus macrosoma* (found in April with three reads using 16S) and *Parapristipoma trilineatum* (recorded in June with 2,368 reads using 12S). Once again, the existence of genetically similar and congeneric species in the region, combined with the limited resolution of the 12S and 16S markers for certain groups (Fontes et al., 2024) and the underrepresentation in reference databases, may have caused the assignment of these reads by the bioinformatic pipeline to the closest available match rather than the true species, as discussed in Ferreira et al. (2024). The remaining species, *Sicyopterus eudentatus*, originates from the Indo-Pacific (Lord et al., 2019) and was identified in May with only eight reads using the 12S marker. Its presence in the sample is highly debatable, pointing out the need to refine procedures to minimize false positives and the importance of cross-verifying morphological and molecular identifications to enhance accuracy and reliability in future studies. The detection of NIS reveals the importance of continuous monitoring in Guadiana Estuary, given the potential ecological impacts that these species may cause to regional biodiversity, especially in cases such as *Parapristipoma trilineatum* recorded in June, where its high read count indicates the need for further investigation, as it may already have a solid presence and a greater ecological impact on the local ecosystem.

This detailed revision is also required for the 10 Elasmobranchii species identified in our samples. Four of these records (*Cetorhinus maximus*, *Galeorhinus galeus*, *Raja undulata*, and *Torpedo marmorata*) were obtained exclusively through eDNA samples, with their presence being easily attributed to DNA shed into the environment during the sampling period. The rays *R. undulata* and *T. marmorata* and the smaller shark *G. galeus* are a regular presence in estuarine environments (Grant et al., 2019), which can explain their detections in our samples; conversely, the presence of *C. maximus*, a large shark found in pelagic open and cooler waters (Sims, 2008), is more debatable. However, this species has been recorded and sighted near the southern Portuguese coast (*pers. obs.*), so its detection could reflect an individual approaching the estuary and shedding DNA that subsequently was transported into it. The presence of six Elasmobranchii species (*Dipturus nidarosiensis*, *Himantura uarnak*, *Mobula mobular*, *Raja brachyura*, *Raja montagui*, and *Scymnodon ringens*) in our ichthyoplankton samples presents additional challenges, as the trawls collected fish eggs and larvae, while most Elasmobranchii species produce benthic egg cases (that can be easily detected) or are live-bearing. A plausible explanation for the presence of Elasmobranchii DNA in these samples is DNA shed by juveniles or adults that can be incorporated into the plankton tow - something possible to occur in the case of *H. uarnak*, *R. brachyura*, and *R. montagui*, rays that inhabit estuarine regions (Ellis et al., 2012; Grant et al., 2019). On the other hand, the presence of deep-sea species like *D. nidarosiensis* (Follesa et al., 2012) and *S. ringens* (White et al., 2015), as well as the oceanic ray *M. mobular* (Croll et al., 2012) is doubtful when looking at their natural habitats, suggesting potential contamination, sequencing errors or bioinformatic misassignments. All these findings expose the challenges of interpreting metabarcoding data, a task made even more complex when the results include species with different ecological preferences from those of the sampled environment. Overall, this study was able to detect an interesting portion of the fish species previously recorded in the Guadiana River Estuary and provided new insights into the local ichthyofauna, including potential shifts in regional biodiversity.

## Conclusions

This study features a quite comprehensive approach to DNA metabarcoding-based monitoring of fish species in a temperate estuary by: i) integrating bulk sampling of developmental stages in the plankton and water eDNA, with the former being backed-up by morphology-based identifications; ii) employing an extensive molecular approach comprising 3 genetic markers (COI, 12S, and 16S) and 4 primer sets, and iii) entailing an in-depth monthly sampling program along 13 months, comprising 3 sampling points in the lower estuary section. Such an approach provided relevant findings both on the performance of DNA-based tools and on the ecology of fish species occurring in the estuary. The aptitude of DNA metabarcoding to provide extensive and precise species-level identifications of ichthyoplankton was patent. The number and diversity of species that was possible to identify greatly expanded compared to the morphology-based approach. Novel and unique data on reproductive ecology was unraveled for many species in the area for the first time. Potential new occurrences were recorded, including possible non-indigenous species detected through DNA metabarcoding of bulk samples. Furthermore, the cross-validation of identification methodologies provided a virtuous feedback mechanism to improve both morphological and molecular approaches.

The unexpected lack of success of eDNA to detect a large fraction of the species that were detected in bulk ichthyoplankton samples is particularly significant. Although these findings require confirmation with further extensive testing, including considerations on sample size, DNA shedding and breakdown rates, their potential implications for DNA based fish monitoring strategies should be investigated. Globally our findings reinforce the added value of the employment of combined sampling of bulk plankton and water eDNA to provide in-depth monitoring of fish occurrence and reproductive ecology for whole fish assemblages, thereby leveraging the worth of such data for fisheries management and fish conservation.

## Supporting information

All suplementary tables

## Acknowledgements

We are grateful to everyone who supported our researchers throughout the 13 months of sampling in the Guadiana Estuary.

## Author contributions

A.O.F., S.D., A.M.P.S. and F.O.C. conception and design; O.M.A. sampling and morphological identification; A.O.F. filtration and DNA extraction of ichthyoplankton samples; C.M. filtration and DNA extraction of eDNA samples; C.E and C.B. amplification and sequencing; A.O.F bioinformatic and statistical analysis; A.O.F, S.D. and F.O.C. writing the article. All authors revised the manuscript.

