## Supplementary material for "From egg to adult: comprehensive (e)DNA metabarcoding monitoring of fish diversity in a temperate estuary": All suplementary tables

**Supplementary Table 1.** Sampling locations in the lower section of the Guadiana River Estuary where monthly collections were conducted, with coordinates indicating approximately the starting points of the trawls. The obtained samples were treated as replicates representing the entire section.

| Sampling Location | Coordinates (DD) |
|---|---|
| ICNF (Upstream) | 37.22390°N, 7.41388°W |
| Doca (Middle) | 37.20203°N, 7.41114°W |
| ISN (Downstream) | 37.18234°N, 7.40839°W |

**Supplementary Table 2.** Larval identifications based on morphology to the lowest possible taxonomic rank, along with corresponding counts for each sampled month in the Guadiana River Estuary.

| Morphology ID | APR | MAY | JUN | JUL | AUG | SEP | OCT | NOV | DEC | JAN | FEB | MAR | APR* | Total |
|---|---|---|---|---|---|---|---|---|---|---|---|---|---|---|
| *Atherina presbyter* | 3 | 4 | 1 | | 3 | | 4 | | 1 | | 3 | 1 | 10 | 30 |
| *Buglossidium luteum* | | | | | | | | | | | 1 | | | 1 |
| *Coryphoblennius galerita* | 1 | 2 | | 1 | 1 | | | | | | | | | 5 |
| *Diplecogaster bimaculata* | | | | 1 | | | | | | | | | | 1 |
| *Diplodus sargus* | 1 | 4 | | | | | | | | | | | | 5 |
| *Echiichthys vipera* | | 1 | | | | 1 | | | | | | | | 2 |
| *Engraulis encrasicolus* | | 44 | 3 | 8 | 30 | 2 | 4 | | 6 | | | 3 | 2 | 102 |
| *Gobius niger* | | | | | | | | | | | 1 | | | 1 |
| *Lipophrys pholis* | 1 | | | | | | | | | | | | | 1 |
| *Parablennius gattorugine* | | 1 | 2 | 3 | | | | | | | | | | 6 |
| *Parablennius pilicornis* | 9 | 23 | 18 | 35 | 18 | 2 | | 1 | | | 1 | 3 | 2 | 112 |
| *Pegusa lascaris* | | | 1 | | | | | | | | 1 | | | 2 |
| *Pomatoschistus microps* | 3 | 22 | 2 | 2 | 12 | | 1 | 3 | 1 | 2 | 10 | 3 | 28 | 89 |
| *Pomatoschistus pictus* | 1 | 68 | 12 | 3 | 20 | | 4 | 1 | | | 2 | | 4 | 115 |
| *Sardina pilchardus* | 1 | 91 | | | 4 | 3 | 4 | 1 | 2 | | 181 | 53 | 8 | 348 |
| *Scophthalmus rhombus* | | | | | | | | | | 1 | | | | 1 |
| *Serranus hepatus* | | 1 | | | | | | | | | | | | 1 |
| *Solea senegalensis* | | | | | | | | 1 | | | 2 | | | 3 |
| *Solea solea* | | | | 4 | | | | | | 4 | | | | 8 |
| *Symphodus melops* | | 1 | | | | | | | | | | | | 1 |
| *Syngnathus typhle* | | | 1 | | 1 | | | | | | | | | 2 |
| *Trachurus mediterraneus* | | | | 1 | 1 | | | | | | | | | 2 |
| *Trachurus trachurus* | | | | 6 | 1 | | 1 | | | | | | | 8 |
| Species Level | | | | | | | | | | | | | | 846 |
| *Callionymus sp.* | | 18 | 4 | 8 | | | | | | | | | | 30 |
| *Diplodus sp.* | | 16 | | | 4 | | | | | | 3 | | | 23 |
| *Parablennius sp.* | | | | 4 | 1 | | | | | | | | 1 | 6 |
| *Pomatoschistus sp.* | 6 | 28 | 1 | | 2 | | | | | | | | 1 | 38 |
| *Solea sp.* | | | | 3 | | | | | | | | | | 3 |
| Genus Level | | | | | | | | | | | | | | 100 |
| Blennidae n.id. | | 1 | | | | 1 | | | | | | | | 2 |
| Gobiidae n.id. | 1 | 5 | | | 7 | | | | | 1 | | | | 14 |
| Soleidae n.id. | | | | | | | | | | | 1 | | | 1 |
| Sparidae n.id. | 2 | 157 | 10 | 10 | 1 | 3 | 1 | | | | 15 | 8 | 26 | 233 |
| Family Level | | | | | | | | | | | | | | 250 |
| n.id. | 3 | 19 | 5 | 5 | | | | | 2 | | 7 | 1 | 6 | 48 |

**Supplementary Table 3.** Number of reads recovered from Illumina MiSeq sequencing and processed through the PMiFish bioinformatics pipeline, including reads merging, primer removal, quality filtering, denoising, taxonomic assignment, and assignment to fish reads for all samples collected over the 13-month period across all primer sets. The values in parentheses indicate the percentage of reads retained at each step relative to the initial raw reads for the corresponding sample.

| Primer Set | Month | Point | Raw reads | Merged | Strip primer | Quality filter | Denoise (Usable) | Tax Assignment | Fish reads |
|---|---|---|---|---|---|---|---|---|---|
| COI FishATL_ Cocktail2 | APR 22 | Up | 95180 (100%) | 76577 (80.45%) | 76573 (80.45%) | 62059 (65.2%) | 61849 (64.98%) | 12436 (13.07%) | 7150 (7.51%) |
| | | Mid | 125807 (100%) | 110418 (87.77%) | 110411 (87.76%) | 93177 (74.06%) | 92683 (73.67%) | 73562 (58.47%) | 24681 (19.62%) |
| | | Down | 90827 (100%) | 77295 (85.1%) | 77294 (85.1%) | 65572 (72.19%) | 64259 (70.75%) | 30630 (33.72%) | 15724 (17.31%) |
| | MAY 22 | Up | 105411 (100%) | 93696 (88.89%) | 93695 (88.89%) | 74314 (70.5%) | 73935 (70.14%) | 49029 (46.51%) | 26138 (24.80%) |
| | | Mid | 105866 (100%) | 90614 (85.59%) | 90601 (85.58%) | 75864 (71.66%) | 75292 (71.12%) | 52303 (49.4%) | 26056 (24.61%) |
| | | Down | 99268 (100%) | 80690 (81.29%) | 80690 (81.29%) | 63793 (64.26%) | 63222 (63.69%) | 16914 (17.04%) | 9024 (9.09%) |
| | JUN 22 | Up | 97386 (100%) | 82054 (84.26%) | 82053 (84.26%) | 64696 (66.43%) | 64648 (66.38%) | 22402 (23%) | 21317 (21.89%) |
| | | Mid | 133284 (100%) | 110732 (83.08%) | 110726 (83.08%) | 87827 (65.89%) | 87564 (65.7%) | 31707 (23.79%) | 27679 (20.77%) |
| | | Down | NA | NA | NA | NA | NA | NA | NA |
| | JUL 22 | Up | 82130 (100%) | 74262 (90.42%) | 74242 (90.4%) | 63601 (77.44%) | 63485 (77.3%) | 8363 (10.18%) | 7446 (9.07%) |
| | | Mid | 93632 (100%) | 77180 (82.43%) | 77171 (82.42%) | 65267 (69.71%) | 64916 (69.33%) | 11131 (11.89%) | 10267 (10.97%) |
| | | Down | 109567 (100%) | 100526 (91.75%) | 100504 (91.73%) | 90219 (82.34%) | 89687 (81.86%) | 23405 (21.36%) | 19703 (17.98%) |
| | AUG 22 | Up | 84434 (100%) | 68139 (80.7%) | 68104 (80.66%) | 57800 (68.46%) | 57629 (68.25%) | 1656 (1.96%) | 1139 (1.35%) |
| | | Mid | 109881 (100%) | 91061 (82.87%) | 91040 (82.85%) | 79749 (72.58%) | 79565 (72.41%) | 12992 (11.82%) | 10234 (9.31%) |
| | | Down | 58355 (100%) | 52283 (89.59%) | 52264 (89.56%) | 45923 (78.7%) | 45857 (78.58%) | 27462 (47.06%) | 20189 (34.60%) |
| | SEP 22 | Up | 50110 (100%) | 41258 (82.33%) | 41237 (82.29%) | 36325 (72.49%) | 36104 (72.05%) | 2967 (5.92%) | 51 (0.10%) |
| | | Mid | 58795 (100%) | 41712 (70.94%) | 41705 (70.93%) | 34163 (58.11%) | 34022 (57.87%) | 3915 (6.66%) | 592 (1.01%) |
| | | Down | 43481 (100%) | 34023 (78.25%) | 34016 (78.23%) | 29360 (67.52%) | 29216 (67.19%) | 5328 (12.25%) | 2382 (5.48%) |
| | OCT 22 | Up | 90062 (100%) | 76552 (85%) | 76539 (84.98%) | 62517 (69.42%) | 62457 (69.35%) | 15301 (16.99%) | 9971 (11.07%) |
| | | Mid | 110580 (100%) | 87238 (78.89%) | 87219 (78.87%) | 76232 (68.94%) | 75701 (68.46%) | 12988 (11.75%) | 5353 (4.84%) |
| | | Down | 63077 (100%) | 49477 (78.44%) | 49461 (78.41%) | 43292 (68.63%) | 43001 (68.17%) | 11846 (18.78%) | 8078 (12.81%) |
| | NOV 22 | Up | 69453 (100%) | 59208 (85.25%) | 59194 (85.23%) | 51257 (73.8%) | 50805 (73.15%) | 17560 (25.28%) | 15637 (22.51%) |
| | | Mid | 85549 (100%) | 75142 (87.84%) | 75134 (87.83%) | 60573 (70.81%) | 60518 (70.74%) | 4022 (4.7%) | 1861 (2.18%) |
| | | Down | 55897 (100%) | 46181 (82.62%) | 46175 (82.61%) | 41081 (73.49%) | 40742 (72.89%) | 4117 (8.91%) | 4 (0.01%) |
| | DEC 22 | Up | 80217 (100%) | 69528 (86.67%) | 69506 (86.65%) | 62658 (78.11%) | 62361 (77.74%) | 20543 (25.61%) | 10467 (13.05%) |
| | | Mid | 31174 (100%) | 23847 (76.5%) | 23843 (76.48%) | 20124 (64.55%) | 20050 (64.32%) | 6769 (21.71%) | 3904 (12.52%) |
| | | Down | 53096 (100%) | 39705 (74.78%) | 39694 (74.76%) | 35341 (66.56%) | 35188 (66.27%) | 15216 (28.66%) | 6963 (13.11%) |
| | JAN 23 | Up | 173280 (100%) | 151250 (87.29%) | 151227 (87.27%) | 133197 (76.87%) | 130598 (75.37%) | 87976 (50.77%) | 63398 (36.59%) |
| | | Mid | 72363 (100%) | 60268 (83.29%) | 60260 (83.27%) | 52802 (72.97%) | 52048 (71.93%) | 26242 (36.26%) | 16250 (22.46%) |
| | | Down | 57850 (100%) | 49924 (86.3%) | 49913 (86.28%) | 42731 (73.87%) | 41755 (72.18%) | 23659 (40.9%) | 13376 (23.12%) |
| | FEB 23 | Up | 234122 (100%) | 192124 (82.06%) | 191999 (82.01%) | 153341 (65.5%) | 151447 (64.69%) | 74804 (31.95%) | 45015 (19.23%) |

| Primer Set | Month | Point | Raw reads | Merged | Strip primer | Quality filter | Denoise (Usable) | Tax Assignment | Fish reads |
|---|---|---|---|---|---|---|---|---|---|
| | | Mid | 50920 (100%) | 41998 (82.48%) | 41980 (82.44%) | 31727 (62.31%) | 31505 (61.87%) | 11897 (23.26%) | 9814 (19.27%) |
| | | Down | 38009 (100%) | 31037 (81.66%) | 31020 (81.61%) | 24509 (64.48%) | 24411 (64.22%) | 8743 (23%) | 5790 (15.23%) |
| | **MAR 23** | Up | 94859 (100%) | 77259 (81.45%) | 77244 (81.43%) | 64018 (67.49%) | 63232 (66.66%) | 33879 (35.72%) | 33470 (35.28%) |
| | | Mid | 57456 (100%) | 46344 (80.66%) | 46336 (80.65%) | 38763 (67.47%) | 38290 (66.64%) | 18101 (31.5%) | 11068 (19.26%) |
| | | Down | 133637 (100%) | 113952 (85.27%) | 113929 (85.25%) | 95159 (71.21%) | 94715 (70.87%) | 35663 (26.69%) | 19637 (14.69%) |
| | **ABR 23** | Up | 52683 (100%) | 44036 (83.59%) | 44025 (83.57%) | 34923 (66.29%) | 34497 (65.48%) | 15744 (29.88%) | 8432 (16.01%) |
| | | Mid | 130644 (100%) | 108897 (83.35%) | 108867 (83.33%) | 86799 (66.44%) | 85770 (65.65%) | 40708 (31.16%) | 22906 (17.53%) |
| | | Down | 92085 (100%) | 75317 (81.79%) | 75292 (81.76%) | 61089 (66.34%) | 60690 (65.91%) | 25244 (27.41%) | 16031 (17.41%) |
| **COI mlCOIintF /LoboR1** | **APR 22** | Up | 123174 (100%) | 111926 (90.87%) | 111926 (90.87%) | 110596 (89.79%) | 110041 (89.34%) | 64031 (51.98%) | 2624 (2.13%) |
| | | Mid | 63422 (100%) | 58308 (91.94%) | 58308 (91.94%) | 57813 (91.16%) | 57677 (90.94%) | 38032 (59.97%) | 1835 (2.89%) |
| | | Down | 121243 (100%) | 111297 (91.8%) | 111297 (91.8%) | 109817 (90.58%) | 108420 (89.42%) | 54521 (44.97%) | 8735 (7.20%) |
| | **MAY 22** | Up | 165009 (100%) | 150186 (91.02%) | 150186 (91.02%) | 148020 (89.7%) | 147417 (89.34%) | 85085 (51.56%) | 9107 (5.52%) |
| | | Mid | 97263 (100%) | 90015 (92.55%) | 90015 (92.55%) | 89361 (91.88%) | 88966 (91.47%) | 59667 (61.35%) | 1132 (1.16%) |
| | | Down | 76370 (100%) | 69967 (91.62%) | 69967 (91.62%) | 69245 (90.67%) | 68394 (89.56%) | 41329 (41.63%) | 4976 (6.52%) |
| | **JUN 22** | Up | 43918 (100%) | 39437 (89.8%) | 39437 (89.8%) | 39148 (89.14%) | 39039 (88.89%) | 20054 (45.66%) | 1134 (2.58%) |
| | | Mid | 89923 (100%) | 80457 (89.47%) | 80456 (89.47%) | 79544 (88.46%) | 79268 (88.15%) | 34544 (38.42%) | 634 (0.71%) |
| | | Down | 115485 (100%) | 101421 (87.82%) | 101421 (87.82%) | 100403 (86.94%) | 99993 (86.59%) | 46066 (39.89%) | 411 (0.36%) |
| | **JUL 22** | Up | 67785 (100%) | 63621 (93.86%) | 63621 (93.86%) | 63079 (93.06%) | 62733 (92.55%) | 19020 (28.06%) | 2574 (3.80%) |
| | | Mid | 79156 (100%) | 75073 (94.84%) | 75073 (94.84%) | 74434 (94.03%) | 74024 (93.52%) | 31200 (39.42%) | 4083 (5.16%) |
| | | Down | 95859 (100%) | 90778 (94.7%) | 90776 (94.7%) | 90107 (94%) | 89579 (93.45%) | 24958 (26.04%) | 2239 (2.34%) |
| | **AUG 22** | Up | 124321 (100%) | 117334 (94.38%) | 117334 (94.38%) | 116480 (93.69%) | 114351 (91.98%) | 44298 (35.63%) | 3872 (3.11%) |
| | | Mid | 69079 (100%) | 65032 (94.14%) | 65032 (94.14%) | 64143 (92.85%) | 64037 (92.7%) | 24488 (35.45%) | 6181 (8.95%) |
| | | Down | NA | NA | NA | NA | NA | NA | NA |
| | **SEP 22** | Up | 74766 (100%) | 70614 (94.45%) | 70614 (94.45%) | 70060 (93.71%) | 69727 (93.26%) | 18766 (25.1%) | 48 (0.06%) |
| | | Mid | 121432 (100%) | 114838 (94.57%) | 114838 (94.57%) | 114036 (93.91%) | 113100 (93.14%) | 38448 (31.66%) | 435 (0.36%) |
| | | Down | 134230 (100%) | 126634 (94.34%) | 126634 (94.34%) | 125709 (93.65%) | 124439 (92.71%) | 64404 (47.98%) | 4904 (3.65%) |
| | **OCT 22** | Up | 176260 (100%) | 167188 (94.85%) | 167188 (94.85%) | 163310 (92.65%) | 162488 (92.19%) | 64221 (36.43%) | 10197 (5.79%) |
| | | Mid | 159170 (100%) | 150028 (94.26%) | 150027 (94.26%) | 148431 (93.25%) | 147107 (92.42%) | 52233 (32.82%) | 4757 (2.99%) |
| | | Down | 154469 (100%) | 150028 (94.26%) | 150027 (94.26%) | 148431 (93.25%) | 147107 (92.42%) | 73113 (47.33%) | 13587 (8.80%) |
| | **NOV 22** | Up | 162239 (100%) | 152479 (93.98%) | 152479 (93.98%) | 151255 (93.23%) | 150728 (92.9%) | 78112 (48.15%) | 27604 (17.01%) |
| | | Mid | 97748 (100%) | 91170 (93.27%) | 91170 (93.27%) | 90132 (92.21%) | 89803 (91.87%) | 12411 (12.7%) | 2070 (2.12%) |
| | | Down | 138903 (100%) | 130546 (93.98%) | 130545 (93.98%) | 129523 (93.25%) | 128963 (92.84%) | 58639 (42.22%) | 463 (0.33%) |
| | **DEC 22** | Up | 119029 (100%) | 110997 (93.25%) | 110996 (93.25%) | 110165 (92.55%) | 109839 (92.28%) | 18458 (15.51%) | 5994 (5.04%) |
| | | Mid | 135978 (100%) | 127367 (93.67%) | 127367 (93.67%) | 126100 (92.74%) | 123802 (91.05%) | 48269 (35.5%) | 5762 (4.24%) |
| | | Down | 150938 (100%) | 142806 (94.61%) | 142806 (94.61%) | 141711 (93.89%) | 139489 (92.41%) | 68479 (45.37%) | 8492 (5.63%) |
| | **JAN 23** | Up | 125958 (100%) | 118749 (94.28%) | 118748 (94.28%) | 117901 (93.6%) | 116547 (92.53%) | 65723 (52.18%) | 11502 (9.13%) |

| Primer Set | Month | Point | Raw reads | Merged | Strip primer | Quality filter | Denoise (Usable) | Tax Assignment | Fish reads |
|---|---|---|---|---|---|---|---|---|---|
| | | Mid | 115592 (100%) | 109147 (94.42%) | 109144 (94.42%) | 108291 (93.68%) | 105811 (91.54%) | 73882 (63.92%) | 6537 (5.66%) |
| | | Down | 146283 (100%) | 137029 (93.67%) | 137028 (93.67%) | 135871 (92.88%) | 134228 (91.76%) | 86750 (59.3%) | 11994 (8.20%) |
| | FEB 23 | Up | 54270 (100%) | 49342 (90.92%) | 49338 (90.91%) | 48339 (89.07%) | 47792 (88.06%) | 28139 (51.85%) | 1141 (2.10%) |
| | | Mid | 86685 (100%) | 78781 (90.88%) | 78781 (90.88%) | 77155 (89.01%) | 76597 (88.36%) | 56752 (65.47%) | 3499 (4.04%) |
| | | Down | 71660 (100%) | 65272 (91.09%) | 65272 (91.09%) | 64016 (89.33%) | 63641 (88.81%) | 45214 (63.1%) | 6883 (9.61%) |
| | MAR 23 | Up | 174357 (100%) | 158552 (90.94%) | 158531 (90.92%) | 155106 (88.96%) | 154259 (88.47%) | 31497 (18.06%) | 4800 (2.75%) |
| | | Mid | 77297 (100%) | 70640 (91.39%) | 70640 (91.39%) | 69357 (89.73%) | 67979 (87.95%) | 22750 (29.43%) | 1732 (2.24%) |
| | | Down | 170697 (100%) | 157618 (92.34%) | 157612 (92.33%) | 154792 (90.68%) | 151656 (88.85%) | 66939 (39.22%) | 6325 (3.71%) |
| | ABR 23 | Up | 89457 (100%) | 81821 (91.46%) | 81821 (91.46%) | 80226 (89.68%) | 78306 (87.53%) | 56947 (63.66%) | 4600 (5.14%) |
| | | Mid | 54578 (100%) | 50557 (92.63%) | 50557 (92.63%) | 49806 (91.26%) | 48406 (88.69%) | 36285 (66.48%) | 1104 (2.02%) |
| | | Down | 69482 (100%) | 63656 (91.62%) | 63655 (91.61%) | 62411 (89.82%) | 60305 (86.79%) | 45664 (65.72%) | 1788 (2.57%) |
| 12S miFISH-U+E | APR 22 | Up | 75387 (100%) | 69478 (92.16%) | 69478 (92.16%) | 69431 (92.1%) | 66275 (87.91%) | 32116 (42.6%) | 32116 (42.60%) |
| | | Mid | 62348 (100%) | 57535 (92.28%) | 57533 (92.28%) | 57475 (92.18%) | 57055 (91.51%) | 16137 (25.88%) | 16137 (25.88%) |
| | | Down | 72837 (100%) | 70902 (97.34%) | 70902 (97.34%) | 70868 (97.3%) | 67365 (92.49%) | 23819 (32.7%) | 23819 (32.72%) |
| | MAY 22 | Up | 78839 (100%) | 76320 (96.8%) | 76319 (96.8%) | 76244 (96.71%) | 73467 (93.19%) | 23708 (30.07%) | 23708 (30.07%) |
| | | Mid | 67121 (100%) | 62112 (92.54%) | 62107 (92.53%) | 62084 (92.5%) | 60829 (90.63%) | 27497 (40.97%) | 27497 (40.97%) |
| | | Down | 74595 (100%) | 68729 (92.14%) | 68727 (92.13%) | 68674 (92.06%) | 67120 (89.98%) | 24323 (32.61%) | 24323 (32.61%) |
| | JUN 22 | Up | 81911 (100%) | 75837 (92.58%) | 75836 (92.58%) | 75787 (92.52%) | 74219 (90.61%) | 8193 (10%) | 8193 (10.00%) |
| | | Mid | 58458 (100%) | 54106 (92.56%) | 54099 (92.54%) | 54042 (92.45%) | 53132 (90.89%) | 10943 (18.72%) | 10943 (18.72%) |
| | | Down | 79253 (100%) | 72400 (91.35%) | 72398 (91.35%) | 72311 (91.24%) | 71648 (90.4%) | 4 (0.01%) | 4 (0.01%) |
| | JUL 22 | Up | 56295 (100%) | 54574 (96.94%) | 54572 (96.94%) | 54551 (96.9%) | 53605 (95.22%) | 11242 (19.97%) | 11242 (19.97%) |
| | | Mid | 65465 (100%) | 63454 (96.93%) | 63454 (96.93%) | 63425 (96.88%) | 61882 (94.53%) | 12929 (19.75%) | 12929 (19.75%) |
| | | Down | 62394 (100%) | 60651 (97.21%) | 60649 (97.2%) | 60637 (97.18%) | 60108 (96.34%) | 8054 (12.91%) | 8054 (12.91%) |
| | AUG 22 | Up | 83359 (100%) | 80697 (96.81%) | 80697 (96.81%) | 80666 (96.77%) | 77968 (93.53%) | 20295 (24.35%) | 20295 (24.35%) |
| | | Mid | 95379 (100%) | 92171 (96.64%) | 92171 (96.64%) | 92150 (96.61%) | 90038 (94.4%) | 29681 (31.12%) | 29681 (31.12%) |
| | | Down | 79976 (100%) | 77676 (97.12%) | 77675 (97.12%) | 77662 (97.11%) | 76031 (95.07%) | 28244 (35.32%) | 28244 (35.32%) |
| | SEP 22 | Up | 57538 (100%) | 55606 (96.64%) | 55605 (96.64%) | 55586 (96.61%) | 54367 (94.49%) | 200 (0.35%) | 200 (0.35%) |
| | | Mid | 77420 (100%) | 75098 (97%) | 75098 (97%) | 75034 (96.92%) | 74207 (95.85%) | 21256 (27.46%) | 21256 (27.46%) |
| | | Down | 68761 (100%) | 66516 (96.74%) | 66516 (96.74%) | 66461 (96.66%) | 63693 (92.63%) | 21156 (30.77%) | 21156 (30.77%) |
| | OCT 22 | Up | 94836 (100%) | 91845 (96.85%) | 91843 (96.84%) | 91801 (96.8%) | 88590 (93.41%) | 45410 (47.88%) | 45410 (47.88%) |
| | | Mid | 114096 (100%) | 110626 (96.96%) | 110625 (96.96%) | 110592 (96.93%) | 107765 (94.45%) | 54287 (47.58%) | 54287 (47.58%) |
| | | Down | 74203 (100%) | 72034 (97.08%) | 72032 (97.07%) | 71984 (97.01%) | 70428 (94.91%) | 26210 (35.32%) | 26210 (35.32%) |
| | NOV 22 | Up | 62935 (100%) | 60558 (96.22%) | 60556 (96.22%) | 60521 (96.16%) | 58416 (92.82%) | 24062 (38.23%) | 24062 (38.23%) |
| | | Mid | 57741 (100%) | 55628 (96.34%) | 55625 (96.34%) | 55601 (96.29%) | 53379 (92.45%) | 24441 (42.33%) | 24441 (42.33%) |
| | | Down | 60146 (100%) | 57733 (95.99%) | 57732 (95.99%) | 57659 (95.87%) | 56025 (93.15%) | 24555 (40.83%) | 24555 (40.83%) |
| | DEC 22 | Up | 66181 (100%) | 63535 (96%) | 63531 (96%) | 63495 (95.94%) | 60944 (92.09%) | 24893 (37.61%) | 24893 (37.61%) |

| Primer Set | Month | Point | Raw reads | Merged | Strip primer | Quality filter | Denoise (Usable) | Tax Assignment | Fish reads |
|---|---|---|---|---|---|---|---|---|---|
| | | Mid | 68364 (100%) | 66032 (96.59%) | 66032 (96.59%) | 65994 (96.53%) | 62454 (91.36%) | 47971 (70.17%) | 47971 (70.17%) |
| | | Down | 89068 (100%) | 86278 (96.87%) | 86273 (96.86%) | 86145 (96.72%) | 84352 (94.71%) | 36646 (41.14%) | 36646 (41.14%) |
| | **JAN 23** | Up | 128958 (100%) | 124938 (96.88%) | 124936 (96.88%) | 124783 (96.76%) | 121857 (94.49%) | 95972 (74.42%) | 95972 (74.42%) |
| | | Mid | 94787 (100%) | 91846 (96.9%) | 91846 (96.9%) | 91786 (96.83%) | 88051 (92.89%) | 67619 (71.34%) | 67619 (71.34%) |
| | | Down | 73817 (100%) | 71392 (96.71%) | 71392 (96.71%) | 71342 (96.65%) | 68880 (93.31%) | 17623 (23.87%) | 17623 (23.87%) |
| | **FEB 23** | Up | 70403 (100%) | 66875 (94.99%) | 66873 (94.99%) | 66803 (94.89%) | 63208 (89.78%) | 33695 (47.86%) | 33695 (47.86%) |
| | | Mid | 73487 (100%) | 69472 (94.54%) | 69472 (94.54%) | 69423 (94.47%) | 66474 (90.46%) | 39692 (54.01%) | 39692 (54.01%) |
| | | Down | 30294 (100%) | 28824 (95.15%) | 28823 (95.14%) | 28805 (95.08%) | 27600 (91.11%) | 12593 (41.57%) | 12593 (41.57%) |
| | **MAR 23** | Up | 33048 (100%) | 31494 (95.3%) | 31494 (95.3%) | 31454 (95.18%) | 30498 (92.28%) | 7122 (21.55%) | 7122 (21.55%) |
| | | Mid | 39885 (100%) | 37647 (94.39%) | 37647 (94.39%) | 37509 (94.04%) | 35901 (90.01%) | 14563 (36.51%) | 14563 (36.51%) |
| | | Down | 27009 (100%) | 25591 (94.75%) | 25590 (94.75%) | 25551 (94.6%) | 24339 (90.11%) | 2394 (8.86%) | 2394 (8.86%) |
| | **ABR 23** | Up | 71728 (100%) | 67882 (94.64%) | 67882 (94.64%) | 67634 (94.29%) | 63922 (89.12%) | 33655 (46.92%) | 33655 (46.92%) |
| | | Mid | 88881 (100%) | 84460 (95.03%) | 84459 (95.02%) | 84361 (94.91%) | 78912 (88.78%) | 44546 (50.12%) | 44546 (50.12%) |
| | | Down | 89860 (100%) | 85927 (95.62%) | 85926 (95.62%) | 85865 (95.55%) | 82363 (91.66%) | 43014 (47.87%) | 43014 (47.87%) |
| **16S Fish16** | **APR 22** | Up | 96862 (100%) | 85172 (87.93%) | 85158 (87.92%) | 84575 (87.31%) | 81034 (83.66%) | 53303 (55.03%) | 53234 (54.96%) |
| | | Mid | 115171 (100%) | 101987 (88.55%) | 101975 (88.54%) | 101678 (88.28%) | 100642 (87.38%) | 46031 (39.97%) | 44015 (38.22%) |
| | | Down | 95595 (100%) | 91462 (95.68%) | 91461 (95.68%) | 91102 (95.3%) | 86328 (90.31%) | 43202 (45.19%) | 43201 (45.19%) |
| | **MAY 22** | Up | 101097 (100%) | 96770 (95.72%) | 96723 (95.67%) | 96631 (95.58%) | 96045 (95%) | 37806 (37.4%) | 37757 (37.35%) |
| | | Mid | 102274 (100%) | 90820 (88.8%) | 90800 (88.78%) | 90709 (88.69%) | 89650 (87.66%) | 39551 (38.67%) | 39534 (38.65%) |
| | | Down | 126908 (100%) | 112033 (88.28%) | 112025 (88.27%) | 111653 (87.98%) | 110660 (87.2%) | 57233 (45.1%) | 57233 (45.10%) |
| | **JUN 22** | Up | 99297 (100%) | 88830 (89.46%) | 88820 (89.45%) | 88323 (88.95%) | 87919 (88.54%) | 21142 (21.29%) | 21112 (21.26%) |
| | | Mid | 129376 (100%) | 115112 (88.97%) | 115074 (88.95%) | 114267 (88.32%) | 113392 (87.65%) | 42408 (32.78%) | 28818 (22.27%) |
| | | Down | NA | NA | NA | NA | NA | NA | NA |
| | **JUL 22** | Up | NA | NA | NA | NA | NA | NA | NA |
| | | Mid | 120494 (100%) | 116198 (96.43%) | 116170 (96.41%) | 116029 (96.29%) | 115074 (95.5%) | 63696 (52.86%) | 63686 (52.85%) |
| | | Down | 67969 (100%) | 66176 (97.36%) | 66150 (97.32%) | 66129 (97.29%) | 66087 (97.23%) | 13891 (20.44%) | 13805 (20.31%) |
| | **AUG 22** | Up | 90073 (100%) | 88261 (97.99%) | 88234 (97.96%) | 88213 (97.94%) | 88212 (97.93%) | 275 (0.31%) | 256 (0.28%) |
| | | Mid | 88304 (100%) | 84772 (96%) | 84722 (95.94%) | 84416 (95.6%) | 83873 (94.98%) | 37590 (42.57%) | 35195 (39.86%) |
| | | Down | 122432 (100%) | 117834 (96.24%) | 117793 (96.21%) | 117722 (96.15%) | 116114 (94.84%) | 65911 (53.83%) | 64976 (53.07%) |
| | **SEP 22** | Up | 38319 (100%) | 36600 (95.51%) | 36583 (95.47%) | 36263 (94.63%) | 35439 (92.48%) | 1860 (4.85%) | 1814 (4.73%) |
| | | Mid | 53413 (100%) | 52013 (97.38%) | 52007 (97.37%) | 51989 (97.33%) | 51935 (97.23%) | 5496 (10.29%) | 4738 (8.87%) |
| | | Down | 68004 (100%) | 65727 (96.65%) | 65721 (96.64%) | 65474 (96.28%) | 65036 (95.64%) | 12137 (17.85%) | 11577 (17.02%) |
| | **OCT 22** | Up | 80478 (100%) | 77241 (95.98%) | 77211 (95.94%) | 76905 (95.56%) | 76009 (94.45%) | 32065 (39.84%) | 31943 (39.69%) |
| | | Mid | 81581 (100%) | 78845 (96.65%) | 78843 (96.64%) | 78634 (96.39%) | 78202 (95.86%) | 24720 (30.3%) | 24697 (30.27%) |
| | | Down | 79654 (100%) | 77148 (96.85%) | 77134 (96.84%) | 76754 (96.36%) | 75804 (95.17%) | 13442 (16.88%) | 13424 (16.85%) |
| | **NOV 22** | Up | 42024 (100%) | 39598 (94.23%) | 39566 (94.15%) | 39172 (93.21%) | 37041 (88.14%) | 18479 (43.97%) | 18425 (43.84%) |

| Primer Set | Month | Point | Raw reads | Merged | Strip primer | Quality filter | Denoise (Usable) | Tax Assignment | Fish reads |
|---|---|---|---|---|---|---|---|---|---|
| | | Mid | 52060 (100%) | 49511 (95.1%) | 49502 (95.09%) | 49230 (94.56%) | 46277 (88.89%) | 33022 (63.43%) | 32766 (62.94%) |
| | | Down | 46792 (100%) | 44705 (95.54%) | 44692 (95.51%) | 44230 (94.52%) | 42786 (91.44%) | 19279 (41.2%) | 19075 (40.77%) |
| | **DEC 22** | Up | 39322 (100%) | 37758 (96.02%) | 37734 (95.96%) | 37401 (95.11%) | 36604 (93.09%) | 5280 (13.43%) | 5280 (13.43%) |
| | | Mid | 94932 (100%) | 91057 (95.92%) | 91044 (95.9%) | 90867 (95.72%) | 89872 (94.67%) | 49325 (51.96%) | 49289 (51.92%) |
| | | Down | 59226 (100%) | 57395 (96.91%) | 57376 (96.88%) | 57203 (96.58%) | 57050 (96.33%) | 15354 (25.92%) | 15343 (25.91%) |
| | **JAN 23** | Up | 46955 (100%) | 44373 (94.5%) | 44349 (94.45%) | 44000 (93.71%) | 43298 (92.21%) | 25725 (54.79%) | 25505 (54.32%) |
| | | Mid | 82301 (100%) | 78391 (95.25%) | 78373 (95.23%) | 78050 (94.83%) | 75837 (92.15%) | 41576 (50.52%) | 41535 (50.47%) |
| | | Down | 52338 (100%) | 49986 (95.51%) | 49964 (95.46%) | 49828 (95.2%) | 48798 (93.24%) | 32608 (62.3%) | 24183 (46.21%) |
| | **FEB 23** | Up | 100812 (100%) | 91415 (90.68%) | 91310 (90.57%) | 89959 (89.23%) | 88005 (87.3%) | 49520 (49.12%) | 49496 (49.10%) |
| | | Mid | 90992 (100%) | 83101 (91.33%) | 83060 (91.28%) | 82149 (90.28%) | 80852 (88.86%) | 51590 (56.7%) | 51585 (56.69%) |
| | | Down | 119678 (100%) | 109684 (91.65%) | 109672 (91.64%) | 108099 (90.32%) | 106037 (88.6%) | 57516 (48.06%) | 57471 (48.02%) |
| | **MAR 23** | Up | 44435 (100%) | 39610 (89.14%) | 39427 (88.73%) | 38673 (87.03%) | 38433 (86.49%) | 25399 (57.16%) | 25378 (57.11%) |
| | | Mid | 121477 (100%) | 109603 (90.23%) | 109501 (90.14%) | 107814 (88.75%) | 106320 (87.52%) | 71156 (58.58%) | 71142 (58.56%) |
| | | Down | 78356 (100%) | 70267 (89.68%) | 70186 (89.57%) | 67878 (86.63%) | 63668 (81.25%) | 16544 (21.11%) | 16517 (21.08%) |
| | **ABR 23** | Up | 83816 (100%) | 75266 (89.8%) | 75190 (89.71%) | 73660 (87.88%) | 70655 (84.3%) | 43372 (51.75%) | 43368 (51.74%) |
| | | Mid | 63816 (100%) | 57860 (90.67%) | 57838 (90.63%) | 56627 (88.73%) | 53921 (84.49%) | 32620 (51.12%) | 32610 (51.10%) |
| | | Down | 105136 (100%) | 91176 (86.72%) | 91050 (86.6%) | 89732 (85.35%) | 88633 (84.3%) | 61157 (58.17%) | 60847 (57.87%) |

**Supplementary Table 4.** Number of reads assigned to each fish species using the COI primer set FishATL_Cocktail2 over the 13-month sampling period.

**Primer Set: FishATL_ Cocktail2 (COI)**

| Species | APR | MAY | JUN | JUL | AUG | SEP | OCT | NOV | DEC | JAN | FEB | MAR | APR* |
|---|---|---|---|---|---|---|---|---|---|---|---|---|---|
| *Alosa fallax* | - | - | - | - | - | - | - | - | 426 | - | - | - | - |
| *Argyrosomus regius* | - | - | 3464 | 16 | - | - | - | - | - | - | - | - | 401 |
| *Arnoglossus laterna* | - | - | - | - | - | - | - | - | 1 | 2341 | 1073 | 2 | 141 |
| *Arnoglossus thori* | 31 | 93 | 5313 | 988 | 320 | 184 | - | 1 | - | - | - | 8895 | - |
| *Atherina presbyter* | 171 | 58 | - | - | - | - | - | - | 294 | - | - | - | 26 |
| *Boops boops* | - | - | - | - | - | - | - | - | - | - | 2598 | 629 | - |
| *Buglossidium luteum* | 1882 | 721 | - | - | - | - | - | - | - | - | 8 | 37 | 203 |
| *Callionymus risso* | 26 | - | - | - | - | - | - | - | - | - | 5 | 2 | - |
| *Chelon labrosus* | - | 1185 | - | - | - | - | - | - | 3 | 207 | - | - | - |
| *Chelon ramada* | - | - | - | 12 | - | 6 | - | - | 34 | 69 | 1 | - | 13 |
| *Chelon saliens* | - | - | 1044 | - | - | - | - | - | - | 18 | - | - | - |
| *Coris julis* | - | - | 7210 | 148 | - | - | - | - | - | - | - | - | - |
| *Coryphoblennius galerita* | 13 | - | - | - | - | - | - | - | - | - | - | - | - |
| *Cynoscion regalis* | - | - | - | - | - | - | - | - | - | - | - | - | 97 |
| *Dagetichthys lusitanicus* | - | - | 12045 | 16448 | 303 | 330 | - | - | - | - | - | - | - |
| *Dicentrarchus labrax* | - | - | - | - | - | - | - | - | - | 2 | - | - | - |
| *Dicentrarchus punctatus* | - | - | - | - | - | - | - | - | - | - | 5 | - | - |
| *Dicologlossa cuneata* | 656 | - | 568 | - | - | - | 106 | 1 | - | 3476 | 29 | - | 57 |
| *Diplodus annularis* | - | - | - | 1134 | - | - | - | - | - | - | - | - | - |
| *Diplodus bellottii* | 11982 | 19580 | 9520 | 3446 | 4314 | - | - | - | - | - | 11128 | 20171 | 9711 |
| *Diplodus sargus* | - | 35 | - | - | 1 | - | 4071 | - | - | - | 3301 | 2393 | 3185 |
| *Diplodus vulgaris* | - | - | - | - | - | - | - | - | 855 | 3 | - | 8959 | 1 |
| *Engraulis encrasicolus* | 1304 | 8904 | 1793 | 8091 | 20786 | 1292 | 10598 | - | 6101 | 94 | 4933 | - | 2698 |
| *Gobius couchi* | - | - | - | - | - | - | - | - | - | - | - | 2702 | 221 |
| *Gobius niger* | - | - | - | - | - | - | - | - | - | - | - | - | 1 |
| *Gobius paganellus* | - | - | - | - | - | - | - | - | - | - | 4848 | - | - |
| *Hypleurochilus bananensis* | - | - | - | - | - | - | - | - | 1 | - | - | - | - |
| *Lepidotrigla cavillone* | - | - | - | - | - | - | - | - | - | - | - | - | 57 |
| *Lipophrys pholis* | 25 | - | - | - | - | - | - | - | - | - | - | - | - |
| *Lithognathus mormyrus* | - | - | - | - | 587 | 436 | - | - | - | - | - | - | - |
| *Macroramphosus scolopax* | - | - | - | - | - | - | - | - | - | - | 61 | - | - |
| *Merluccius merluccius* | - | - | 249 | - | - | - | - | - | - | - | - | - | - |
| *Microchirus boscanion* | - | 858 | - | - | - | - | - | - | - | - | - | - | 728 |
| *Mullus barbatus* | - | 603 | - | 498 | - | - | - | - | - | - | - | - | - |
| *Opeatogenys gracilis* | - | - | - | 1087 | - | - | - | - | - | - | - | - | - |
| *Pagellus acarne* | - | - | - | - | - | - | 1 | - | - | - | - | - | - |
| *Pagellus erythrinus* | - | - | - | 79 | - | 693 | - | - | - | - | - | - | 4 |
| *Parablennius gattorugine* | - | - | - | - | - | - | - | - | - | - | 1 | - | 1 |
| *Parablennius incognitus* | - | 1319 | - | 86 | - | - | - | - | - | - | - | - | - |
| *Parablennius pilicornis* | 5653 | - | - | - | - | - | - | - | - | - | 1 | 12961 | 787 |
| *Parablennius sanguinolentus* | - | 1999 | - | - | - | - | - | - | - | - | - | - | - |
| *Pegusa lascaris* | 1017 | 387 | - | - | - | 2 | - | 1992 | - | 1852 | 66 | - | 8 |
| *Pomadasys incisus* | - | - | - | 1434 | - | - | - | - | - | - | - | - | - |

**Primer Set: FishATL_ Cocktail2 (COI)**

| Species | APR | MAY | JUN | JUL | AUG | SEP | OCT | NOV | DEC | JAN | FEB | MAR | APR* |
|---|---|---|---|---|---|---|---|---|---|---|---|---|---|
| *Pomatomus saltatrix* | - | - | - | 784 | - | - | - | - | - | - | - | - | - |
| *Pomatoschistus microps* | 14795 | 13594 | 228 | 2289 | 567 | 1 | 6563 | 4658 | 132 | 24082 | 10935 | 10 | 11912 |
| *Pomatoschistus norvegicus* | - | - | - | - | - | - | - | - | - | - | 1 | - | - |
| *Salaria pavo* | - | - | 1028 | 4 | 1 | - | - | - | 1 | - | - | - | - |
| *Sardina pilchardus* | 586 | 10733 | 216 | 392 | 1 | 10 | 727 | 1244 | 13409 | 1216 | 17936 | 4028 | 16367 |
| *Sardinella aurita* | - | - | - | - | 4672 | - | - | - | 47 | - | - | - | - |
| *Sarpa salpa* | - | 653 | - | 172 | - | - | - | 145 | 25 | 27 | - | - | - |
| *Scomber colias* | - | - | - | - | - | - | - | - | - | - | - | - | 202 |
| *Scorpaena notata* | 350 | - | - | - | - | - | 1 | - | - | - | - | - | - |
| *Serranus hepatus* | - | - | - | - | - | - | - | - | - | - | - | - | 5 |
| *Solea senegalensis* | 8541 | 96 | - | - | 1 | 21 | 1334 | 9293 | 1 | 13303 | 529 | 3385 | 4 |
| *Sparus aurata* | 497 | 376 | 6299 | - | 4 | 50 | - | - | - | - | - | - | - |
| *Spicara maena* | 1 | - | - | - | - | - | - | - | - | - | - | - | - |
| *Spondyliosoma cantharus* | - | - | - | - | - | - | - | - | - | - | - | - | 1 |
| *Symphodus bailloni* | - | - | - | - | - | - | - | - | - | - | - | - | 538 |
| *Syngnathus rostellatus* | - | - | - | - | 5 | - | - | 168 | 4 | 46334 | 2537 | 1 | - |
| *Trachurus trachurus* | - | - | - | - | - | - | 1 | - | - | - | 623 | - | - |
| *Umbrina canariensis* | - | - | - | 308 | - | - | - | - | - | - | - | - | - |
| *Zeus faber* | - | 24 | - | - | - | - | - | - | - | - | - | - | - |

**Supplementary Table 5.** Number of reads assigned to each fish species using the COI primer pair mICOIintF/LoboR1 for the 13-month sampling period.

**Primer Set: mICOIintF/LoboR1 (COI)**

| Sample | APR | MAY | JUN | JUL | AUG | SEP | OCT | NOV | DEC | JAN | FEB | MAR | APR* |
|---|---|---|---|---|---|---|---|---|---|---|---|---|---|
| *Anthias anthias* | - | - | - | - | - | - | - | 55 | 16 | - | - | - | - |
| *Argyrosomus regius* | - | - | 432 | - | - | - | - | - | - | - | - | - | 19 |
| *Arnoglossus laterna* | - | - | - | - | - | - | - | - | - | 337 | 49 | - | 2 |
| *Arnoglossus thori* | - | 1 | - | 229 | 33 | 17 | 110 | - | - | - | - | 380 | - |
| *Atherina presbyter* | 935 | 133 | - | - | - | - | - | 3 | 5602 | - | - | - | 74 |
| *Belone belone* | - | - | - | - | - | 1 | - | - | - | - | - | - | - |
| *Boops boops* | 2 | - | - | - | - | - | - | - | - | 62 | - | - | - |
| *Buglossidium luteum* | 154 | - | - | - | - | - | - | - | - | - | - | 7 | 3 |
| *Callionymus risso* | 34 | - | 24 | 1 | - | - | - | - | - | - | 63 | 106 | 1 |
| *Chelidonichthys obscurus* | - | 219 | - | - | - | - | - | - | - | - | - | - | - |
| *Chelon labrosus* | - | 55 | 5 | - | - | - | - | - | 12 | 217 | - | 1069 | - |
| *Chelon saliens* | - | 1 | - | - | - | 29 | - | - | - | - | - | - | - |
| *Chromogobius zebratus* | - | - | - | - | - | - | - | - | 2 | - | - | - | - |
| *Coryphoblennius galerita* | 5530 | 1 | - | - | - | - | - | - | - | - | - | - | - |
| *Dagetichthys lusitanicus* | - | - | 82 | 278 | 5 | - | - | - | - | - | - | - | - |
| *Dicentrarchus labrax* | 39 | - | - | - | - | - | - | - | - | 2813 | - | - | - |
| *Dicologlossa cuneata* | 34 | - | - | - | - | - | 1 | - | - | 3756 | 16 | - | 6 |
| *Diplodus annularis* | - | - | - | 27 | - | - | - | - | - | - | - | - | - |
| *Diplodus bellottii* | 1554 | 2620 | 261 | 842 | 82 | - | - | - | - | - | 2058 | 7025 | 1629 |
| *Diplodus sargus* | - | - | - | - | - | 1 | 1360 | - | - | - | 10 | - | 360 |

**Primer Set: mICOlintF/LoboR1 (COI)**

| Sample | APR | MAY | JUN | JUL | AUG | SEP | OCT | NOV | DEC | JAN | FEB | MAR | APR* |
|---|---|---|---|---|---|---|---|---|---|---|---|---|---|
| *Diplodus vulgaris* | - | - | - | - | - | - | - | - | 1 | - | - | 507 | - |
| *Engraulis encrasicolus* | 643 | 10531 | 713 | 3993 | 9111 | 4888 | 23414 | 2695 | 10441 | 793 | 2154 | 1 | 3947 |
| *Gaidropsarus mediterraneus* | - | - | - | - | - | - | - | - | - | - | 5 | - | - |
| *Gobius couchi* | - | - | - | - | - | - | - | - | - | - | - | 2994 | 307 |
| *Gobius niger* | - | - | - | - | - | - | - | - | - | - | - | - | 22 |
| *Lipophrys pholis* | 114 | - | 410 | - | - | - | - | - | - | - | - | - | - |
| *Lithognathus mormyrus* | - | - | - | - | - | 107 | - | - | - | - | - | - | - |
| *Microchirus boscanion* | - | 58 | - | - | - | - | - | - | - | - | - | - | 763 |
| *Mullus barbatus* | - | 19 | - | - | - | 70 | - | - | - | - | - | - | - |
| *Oncorhynchus mykiss* | - | - | - | - | - | - | - | - | - | - | 1 | - | - |
| *Pagellus acarne* | 1 | - | 1 | - | - | 109 | - | 182 | - | 385 | - | - | - |
| *Pagellus erythrinus* | - | 1 | 9 | - | 505 | 86 | - | 48 | - | 112 | - | - | - |
| *Parablennius gattorugine* | - | - | - | - | - | - | - | - | 4 | - | - | - | - |
| *Parablennius incognitus* | - | 1208 | 89 | 1191 | - | - | - | - | - | - | - | - | - |
| *Parablennius sanguinolentus* | - | 11 | - | - | - | - | - | - | - | - | - | - | - |
| *Pegusa lascaris* | 276 | 59 | - | - | - | - | - | 3330 | - | 2401 | 80 | - | 17 |
| *Pomadasys incisus* | - | - | - | 15 | - | - | - | - | - | - | - | - | - |
| *Pomatomus saltatrix* | - | 3 | - | 10 | - | - | - | - | - | - | - | - | - |
| *Pomatoschistus microps* | 1998 | 277 | 38 | 1 | 1 | - | 411 | 116 | - | 685 | 110 | - | 112 |
| *Sardina pilchardus* | 1 | 9 | - | - | - | - | - | - | 132 | 2 | 111 | 215 | 220 |
| *Scomber colias* | - | - | - | - | - | 5 | - | - | - | - | - | - | 1 |
| *Scomberesox saurus* | - | - | - | - | - | - | - | - | 2208 | - | - | - | - |
| *Scorpaena notata* | - | - | - | - | - | - | 438 | - | - | - | - | - | - |
| *Serranus cabrilla* | - | - | - | 91 | - | 29 | - | 444 | - | - | - | - | - |
| *Serranus hepatus* | - | - | - | 2210 | 315 | 1 | 25 | 79 | 1829 | 38 | - | - | 3 |
| *Solea senegalensis* | 1179 | 9 | - | - | - | 3 | 821 | 22875 | 1 | 7810 | 651 | 552 | 6 |
| *Sparus aurata* | 676 | - | 94 | - | - | 41 | 1960 | 293 | - | - | - | - | - |
| *Syngnathus rostellatus* | - | - | 5 | - | 1 | - | 1 | 17 | - | 10622 | 664 | - | - |
| *Trachurus mediterraneus* | 1 | - | - | - | - | - | - | - | - | - | - | - | - |
| *Trachurus trachurus* | - | - | - | - | - | - | - | - | - | - | 5551 | 1 | - |
| *Umbrina canariensis* | - | - | - | 8 | - | - | - | - | - | - | - | - | - |

**Supplementary Table 6.** Number of reads assigned to each fish species using the 12S primer set miFISH U-E for the 13-month sampling period.

**Primer Set: miFISH U-E (12S)**

| Sample | APR | MAY | JUN | JUL | AUG | SEP | OCT | NOV | DEC | JAN | FEB | MAR | APR* |
|---|---|---|---|---|---|---|---|---|---|---|---|---|---|
| *Abudefduf saxatilis* | - | - | - | - | - | - | - | - | 43 | - | - | - | - |
| *Acantholabrus palloni* | - | - | - | - | - | - | - | - | - | - | - | - | 3 |
| *Alosa fallax* | - | - | - | 10 | - | - | - | - | 54 | 5 | - | - | - |
| *Argyrosomus regius* | - | - | 4200 | 23 | - | - | - | - | - | - | - | 1 | 690 |
| *Arnoglossus imperialis* | - | 48 | 1055 | 394 | 545 | - | - | - | - | - | - | 579 | 15 |
| *Arnoglossus laterna* | - | - | - | - | - | - | - | - | - | 605 | 825 | 1 | 30 |
| *Arnoglossus thori* | 13 | - | - | 330 | 111 | 1588 | 1 | - | - | - | - | - | - |
| *Atherina presbyter* | 14765 | 5798 | 102 | 1961 | 852 | - | 9 | 8882 | 24779 | 9 | - | - | 6188 |

**Primer Set: miFISH U-E (12S)**

| Sample | APR | MAY | JUN | JUL | AUG | SEP | OCT | NOV | DEC | JAN | FEB | MAR | APR* |
|---|---|---|---|---|---|---|---|---|---|---|---|---|---|
| *Auxis rochei* | 3 | - | - | - | - | - | - | - | - | - | - | - | - |
| *Balistes capriscus* | - | - | - | - | 5 | - | - | - | - | - | - | - | - |
| *Belone belone* | 2 | - | 13 | - | - | 160 | 4 | - | - | - | - | - | 4 |
| *Boops boops* | 2 | 15 | 1 | - | - | - | 8 | - | - | - | 1911 | 200 | 1 |
| *Buglossidium luteum* | 4094 | 850 | - | - | - | - | - | - | - | - | 27 | 77 | 1103 |
| *Callionymus reticulatus* | - | - | - | - | - | - | - | - | - | - | - | - | 3 |
| *Caranx hippos* | - | - | - | - | - | 3 | - | - | - | - | - | - | - |
| *Cepola macrophthalma* | - | 1 | - | - | - | - | - | - | - | - | - | - | - |
| *Chelidonichthys lucerna* | - | - | - | - | - | - | - | - | - | - | - | - | 2 |
| *Chelidonichthys obscurus* | - | 232 | - | - | - | - | - | - | - | - | - | - | - |
| *Chelon auratus* | - | 2 | - | - | - | - | - | - | 17 | - | 2 | - | - |
| *Chelon labrosus* | 7 | 264 | 133 | 52 | 18 | 38 | - | - | 133 | 650 | 1 | 3 | 34 |
| *Chelon ramada* | - | - | - | 3 | - | 6 | - | - | - | - | - | - | - |
| *Chelon saliens* | - | - | 375 | - | - | 224 | - | - | - | - | - | - | - |
| *Coris julis* | - | - | 2765 | 1081 | 183 | - | 18 | 57 | - | - | - | - | - |
| *Coryphoblennius galerita* | 6969 | - | - | - | - | - | - | - | - | - | - | - | - |
| *Cynoscion regalis* | - | 5 | - | - | 4 | - | - | - | - | - | - | - | 60 |
| *Dentex gibbosus* | - | - | - | - | - | - | - | - | - | 11 | - | - | - |
| *Dicentrarchus labrax* | - | - | - | - | - | 93 | 4 | - | - | 7842 | - | 1 | 6 |
| *Dicentrarchus punctatus* | - | 1347 | - | - | - | - | - | 28 | - | - | 2311 | - | 377 |
| *Diplodus bellottii* | - | 374 | 6571 | - | - | - | 1 | - | - | - | - | - | - |
| *Diplodus sargus* | - | 114 | - | 7651 | 1927 | 34 | 16902 | 120 | 34 | - | 2797 | 18 | 14472 |
| *Diplodus vulgaris* | - | - | - | - | - | - | - | - | 991 | - | - | 4206 | 1 |
| *Dipturus nidarosiensis* | 1 | - | - | - | - | - | - | - | - | - | - | - | - |
| *Echiichthys vipera* | - | - | - | - | - | - | - | - | - | - | - | - | 2 |
| *Engraulis encrasicolus* | 2312 | 13584 | 850 | 13199 | 37001 | 24403 | 43894 | 3 | 17392 | 34 | 3883 | 14 | 9896 |
| *Euthynnus alletteratus* | - | - | - | - | - | - | - | - | 36 | - | - | - | - |
| *Gadiculus argenteus* | - | - | - | - | - | - | 1 | - | - | - | - | - | - |
| *Gobius niger* | 1521 | 54 | 17 | - | 1520 | 12 | - | - | - | - | - | 6 | 1508 |
| *Gobius paganellus* | 17 | - | - | - | - | - | - | - | - | 57 | 3786 | - | - |
| *Helicolenus dactylopterus* | - | - | - | - | 34 | - | - | - | - | - | - | - | - |
| *Himantura uarnak* | - | - | - | - | - | - | - | - | 35 | - | - | - | - |
| *Katsuwonus pelamis* | 3 | - | - | - | - | - | - | - | - | - | - | - | - |
| *Labrus bergylta* | - | - | - | - | - | - | - | - | - | - | - | 1 | - |
| *Lepidotrigla cavillone* | - | - | - | - | - | - | - | - | - | - | - | - | 29 |
| *Lophius budegassa* | 1 | - | - | - | - | - | - | - | - | - | - | - | - |
| *Macroramphosus scolopax* | - | - | - | - | - | - | - | - | - | - | 18 | - | - |
| *Merluccius merluccius* | - | - | - | - | - | - | - | - | - | 1 | - | - | - |
| *Micromesistius poutassou* | 1 | - | - | - | - | - | - | - | - | - | - | - | - |
| *Mobula mobular* | - | - | - | 3 | - | - | - | - | - | - | - | - | - |
| *Mugil cephalus* | - | - | - | 9 | - | - | 23 | - | - | - | - | - | - |
| *Mullus surmuletus* | - | - | 2 | - | - | 8 | 1 | 6 | - | - | - | - | - |
| *Myctophum punctatum* | - | 1 | - | - | - | - | - | 32 | - | - | - | - | - |
| *Nerophis ophidion* | - | 286 | - | - | - | - | - | - | - | - | - | - | - |
| *Pagellus acarne* | - | 2 | 3 | - | - | 7 | 1 | - | - | - | - | - | - |
| *Pagellus erythrinus* | - | 239 | 2 | 23 | 4 | 8007 | - | - | 1 | - | - | - | 2 |

**Primer Set: miFISH U-E (12S)**

| Sample | APR | MAY | JUN | JUL | AUG | SEP | OCT | NOV | DEC | JAN | FEB | MAR | APR* |
|---|---|---|---|---|---|---|---|---|---|---|---|---|---|
| *Pagrus pagrus* | - | 1 | - | - | - | 2 | - | - | - | - | - | - | - |
| *Parablennius gattorugine* | - | - | - | - | - | - | - | - | - | - | 2 | 1 | 6 |
| *Parapristipoma trilineatum* | - | 2368 | - | - | - | - | - | - | - | - | - | - | - |
| *Pegusa lascaris* | - | - | - | - | - | 70 | - | - | - | - | - | - | - |
| *Pomatomus saltatrix* | - | 408 | 250 | 2137 | 2256 | - | - | - | - | - | - | - | - |
| *Pomatoschistus microps* | 34044 | 30547 | 1686 | 3760 | 19882 | 4 | 38876 | 14119 | 608 | 60520 | 17181 | 10 | 31016 |
| *Pomatoschistus pictus* | - | - | - | - | - | - | - | - | - | - | 7 | - | - |
| *Pseudaphya ferreri* | - | 8 | - | 1 | - | 2 | - | 25201 | - | - | - | - | - |
| *Raja montagui* | - | - | - | - | - | - | - | - | - | 2 | - | - | - |
| *Salmo trutta* | - | - | - | - | - | - | - | - | - | - | - | 1 | - |
| *Sardina pilchardus* | 1081 | 18657 | 52 | 142 | 239 | 5690 | 15730 | 10609 | 65381 | 7698 | 40712 | 15623 | 54159 |
| *Sardinella aurita* | - | - | - | 3 | 13480 | 7 | - | - | - | - | - | - | - |
| *Scomber colias* | 18 | - | - | 12 | - | 56 | - | - | - | - | - | - | 159 |
| *Scymnodon ringens* | 23 | - | - | - | - | - | - | - | - | - | - | - | - |
| *Sicyopterus eudentatus* | - | 8 | - | - | - | - | - | - | - | - | - | - | - |
| *Solea senegalensis* | 7440 | 167 | - | 4 | - | 2119 | 10171 | 13928 | 4 | 36794 | 3389 | 3337 | - |
| *Sparus aurata* | 953 | 148 | 353 | 1 | 102 | 54 | 255 | 10 | 2 | 6 | 2 | - | - |
| *Spondyliosoma cantharus* | 2 | - | 1 | - | - | - | - | - | - | - | - | - | - |
| *Symphodus bailloni* | - | - | 106 | - | - | - | - | - | - | - | - | - | 1447 |
| *Symphodus melops* | - | - | - | - | - | - | - | - | - | - | 1 | - | - |
| *Symphodus ocellatus* | - | - | 9 | - | - | - | - | - | - | - | - | - | - |
| *Syngnathus rostellatus* | 320 | - | 593 | - | 1 | - | 1 | 48 | - | 66980 | 7652 | - | 1 |
| *Trachinotus ovatus* | - | - | - | - | - | 2 | 7 | - | - | - | - | - | - |
| *Trachurus mediterraneus* | - | - | - | - | 56 | - | - | - | - | - | - | - | - |
| *Trachurus trachurus* | - | - | 1 | - | - | - | - | 15 | - | - | 1473 | - | 1 |
| *Umbrina canariensis* | - | - | - | 1426 | - | 23 | - | - | - | - | - | - | - |

**Supplementary Table 7.** Number of reads assigned to each fish species using the 16S primer pair Fish 16S for the 13-month sampling period.

**Primer Set: Fish 16S (16S)**

| Sample | APR | MAY | JUN | JUL | AUG | SEP | OCT | NOV | DEC | JAN | FEB | MAR | APR* |
|---|---|---|---|---|---|---|---|---|---|---|---|---|---|
| *Alosa fallax* | - | - | - | 6 | - | - | - | - | 5 | - | - | 6 | - |
| *Argyrosomus regius* | - | - | 12348 | 40 | - | - | - | - | - | - | - | - | 720 |
| *Arnoglossus imperialis* | - | - | - | - | - | - | - | - | - | - | 775 | - | 5 |
| *Arnoglossus laterna* | - | - | - | - | - | - | - | - | - | 718 | 923 | - | 3 |
| *Arnoglossus thori* | 34 | 7 | 2619 | 717 | 2 | 56 | - | - | - | - | - | 908 | 6 |
| *Atherina presbyter* | 18181 | 4247 | 85 | - | 232 | - | - | 6333 | 24742 | 7 | - | - | 2545 |
| *Belone belone* | - | - | - | - | - | - | 3 | - | - | - | - | - | - |
| *Belone svetovidovi* | - | 2 | - | - | - | - | - | - | - | - | - | - | - |
| *Boops boops* | - | 16 | - | - | - | - | - | - | - | - | 1480 | 224 | - |
| *Buglossidium luteum* | 4093 | 362 | 2 | - | - | - | - | - | - | - | 30 | 65 | 423 |
| *Chelidonichthys lucerna* | - | - | - | - | - | - | - | - | - | - | 1 | - | - |
| *Chelidonichthys obscurus* | - | 154 | - | - | - | - | - | - | - | - | - | - | - |
| *Chelon auratus* | - | 3 | - | - | - | - | - | - | - | - | 5 | - | - |

**Primer Set: Fish 16S (16S)**

| Sample | APR | MAY | JUN | JUL | AUG | SEP | OCT | NOV | DEC | JAN | FEB | MAR | APR* |
|---|---|---|---|---|---|---|---|---|---|---|---|---|---|
| *Chelon labrosus* | - | 175 | - | - | - | 1 | - | 1 | - | 109 | - | 1 | - |
| *Chelon ramada* | - | - | - | 8 | - | 2 | - | 6 | 34 | - | - | - | 14 |
| *Chelon saliens* | - | - | 13 | - | - | - | - | - | - | - | - | - | - |
| *Coris julis* | - | - | 11482 | 2916 | 753 | - | - | - | - | - | - | - | - |
| *Coryphoblennius galerita* | 26 | - | - | - | - | - | - | - | - | - | - | - | - |
| *Cynoscion regalis* | - | 11 | - | 5331 | - | - | - | - | - | - | - | - | 102 |
| *Dagetichthys lusitanicus* | - | - | 178 | 24 | - | - | - | - | - | - | - | - | - |
| *Decapterus macrosoma* | 3 | - | - | - | - | - | - | - | - | - | - | - | - |
| *Dentex gibbosus* | - | - | - | - | - | - | - | - | - | 1 | - | - | - |
| *Dicentrarchus labrax* | - | 4 | 2 | - | - | 3 | 2 | - | 1 | 3719 | - | 2 | 6 |
| *Dicentrarchus punctatus* | - | 139 | - | - | - | - | - | - | - | - | 4339 | - | 72 |
| *Dicologlossa cuneata* | 620 | - | 509 | - | - | - | 232 | 1 | - | 9277 | 320 | - | 39 |
| *Diplodus annularis* | - | - | - | - | - | - | - | - | - | - | - | 9 | - |
| *Diplodus bellottii* | 11531 | 17163 | 9446 | 8092 | 12626 | 6 | 1 | - | - | 1 | 26394 | 25556 | 19128 |
| *Diplodus sargus* | - | 230 | - | 208 | 851 | - | 15757 | 210 | 1 | 4 | 4229 | 29 | 22022 |
| *Diplodus vulgaris* | - | - | - | - | - | 1 | - | - | 1333 | - | 2 | 9356 | - |
| *Engraulis encrasicolus* | 2966 | 19641 | 2364 | 22179 | 37147 | 5050 | 26287 | 26 | 5467 | 75 | 4994 | 3 | 11048 |
| *Gobius couchi* | - | - | - | - | - | - | - | - | - | - | - | 3963 | 944 |
| *Gobius niger* | 2 | 461 | 37 | - | 5310 | 18 | - | - | - | - | - | - | 1123 |
| *Gobius paganellus* | 23 | - | - | - | - | - | - | - | - | 36 | 3929 | - | - |
| *Helicolenus dactylopterus* | - | - | - | - | - | - | - | - | - | 4 | - | - | - |
| *Hypleurochilus bananensis* | - | - | 449 | - | - | - | - | 7 | 5 | 51 | - | - | - |
| *Katsuwonus pelamis* | 1 | - | - | - | - | - | - | - | - | - | - | - | - |
| *Lepidotrigla cavillone* | - | - | - | - | - | - | - | - | - | - | - | - | 21 |
| *Lithognathus mormyrus* | - | - | - | - | 4454 | 4804 | 1 | - | - | - | - | - | - |
| *Macroramphosus scolopax* | - | - | - | - | - | - | - | - | - | - | 18 | - | - |
| *Merluccius merluccius* | - | 1 | - | - | - | - | - | - | - | - | - | - | - |
| *Microchirus boscanion* | - | - | - | - | - | - | - | - | - | - | - | - | 363 |
| *Microlipophrys canevae* | - | - | 37 | - | - | - | - | - | - | - | - | - | - |
| *Mugil cephalus* | 4 | - | 308 | 1 | - | 7 | 1 | 1 | 14 | 262 | 1 | 6 | 1 |
| *Mullus barbatus* | - | 114 | - | - | - | - | - | - | - | - | - | - | - |
| *Mullus surmuletus* | - | - | - | - | - | - | 4 | 34 | - | - | - | - | - |
| *Myctophum punctatum* | - | 1 | - | - | - | - | - | - | - | - | - | - | - |
| *Opeatogenys gracilis* | - | - | - | 40 | 17 | - | - | - | - | - | - | - | - |
| *Pagellus erythrinus* | - | 142 | 3 | 4 | - | 2078 | - | - | - | - | - | - | 1 |
| *Parablennius incognitus* | - | 272 | 41 | 189 | - | - | - | - | - | - | - | - | - |
| *Parablennius pilicornis* | 13660 | 3 | - | - | 4 | 1761 | - | 2383 | - | - | - | 8393 | 2095 |
| *Parablennius sanguinolentus* | - | 565 | - | - | - | - | - | - | - | - | - | - | - |
| *Pegusa impar* | - | - | - | - | - | - | - | 4 | - | 7 | 3 | - | - |
| *Pegusa lascaris* | 4009 | 116 | - | - | - | 1 | - | 4584 | - | 6094 | 705 | - | 25 |
| *Pomadasys incisus* | - | - | - | 165 | - | - | - | - | - | - | - | - | - |
| *Pomatomus saltatrix* | - | 202 | 2 | 2978 | 281 | - | - | - | - | - | - | - | - |
| *Pomatoschistus microps* | 65629 | 46924 | 3142 | 3862 | 6745 | 2 | 16104 | 12289 | 20 | 35538 | 20325 | 2 | 21867 |
| *Pomatoschistus norvegicus* | - | - | - | - | - | - | - | - | - | - | 16 | - | - |
| *Pomatoschistus pictus* | - | - | - | - | - | - | - | - | - | - | 2 | - | - |
| *Pseudaphya ferreri* | - | 18 | - | 1 | - | 1 | - | 20152 | - | 1 | - | - | - |

**Primer Set: Fish 16S (16S)**

| Sample | APR | MAY | JUN | JUL | AUG | SEP | OCT | NOV | DEC | JAN | FEB | MAR | APR* |
|---|---|---|---|---|---|---|---|---|---|---|---|---|---|
| *Raja brachyura* | - | - | - | - | - | - | - | - | - | 7 | - | - | - |
| *Raja montagui* | - | - | - | - | - | - | - | - | - | 5 | - | - | - |
| *Salaria pavo* | - | 30 | 43 | 13234 | 1453 | - | - | - | - | - | - | - | - |
| *Sardina pilchardus* | 2910 | 33937 | 27 | 240 | 65 | 4321 | 10347 | 15101 | 38288 | 6757 | 80979 | 56533 | 53292 |
| *Sardinella aurita* | - | - | - | - | 25683 | 1 | - | - | - | - | - | - | - |
| *Sarpa salpa* | - | 136 | - | 429 | - | 1 | - | 27 | 1 | 13 | - | - | - |
| *Scomber colias* | 19 | - | - | - | - | 8 | - | - | - | - | - | - | 114 |
| *Scymnodon ringens* | 15 | - | - | - | - | - | - | - | - | - | - | - | - |
| *Solea senegalensis* | 12418 | 27 | - | - | - | 2 | 1168 | 9106 | - | 21682 | 2193 | 7978 | - |
| *Sparus aurata* | 813 | 77 | 4142 | - | - | 5 | 156 | 1 | 1 | 3 | - | 1 | - |
| *Spicara maena* | 3448 | 9344 | 2279 | 16510 | - | - | - | - | - | - | - | - | 49 |
| *Spondyliosoma cantharus* | 5 | - | - | - | - | - | - | - | - | - | - | - | - |
| *Symphodus bailloni* | - | - | 307 | - | - | - | - | - | - | - | - | - | 797 |
| *Symphodus ocellatus* | - | - | 57 | - | - | - | - | - | - | - | - | - | - |
| *Syngnathus rostellatus* | 4 | - | 8 | - | - | - | - | - | - | 6851 | 646 | - | - |
| *Trachurus mediterraneus* | 1 | - | - | - | 221 | - | - | - | - | - | - | - | - |
| *Trachurus trachurus* | - | - | - | - | - | - | 1 | - | - | 1 | 6239 | 2 | - |
| *Trisopterus luscus* | - | - | - | - | - | - | - | - | - | - | 4 | - | - |
| *Zebrus zebrus* | - | - | - | 317 | 4583 | - | - | - | - | - | - | - | - |

**Supplementary Table 8.** Taxonomic classification of the identified fish species, listed at the species level along with their respective authority and habitat classification, as referenced in WoRMS (accessed 6th November 2024). Species are organized alphabetically by order name (↓). Habitat categories: M – Marine; B – Brackish; F – Freshwater.

| Scientific Name | Author | Class | Order ↓ | Family | Genus | Habitat |
|---|---|---|---|---|---|---|
| *Capros aper* | Linnaeus, 1758 | Teleostei | Acanthuriformes | Caproidae | Capros | M |
| *Conger conger* | Linnaeus, 1758 | Teleostei | Anguilliformes | Congridae | Conger | M |
| *Atherina presbyter* | Cuvier, 1829 | Teleostei | Atheriniformes | Atherinidae | Atherina | M / B |
| *Belone belone* | Linnaeus, 1760 | Teleostei | Beloniformes | Belonidae | Belone | M / B |
| *Belone svetovidovi* | Collette & Parin, 1970 | Teleostei | Beloniformes | Belonidae | Belone | M |
| *Scomberesox saurus* | Walbaum, 1792 | Teleostei | Beloniformes | Scomberesocidae | Scomberesox | M |
| *Coryphoblennius galerita* | Linnaeus, 1758 | Teleostei | Blenniiformes | Blenniidae | Coryphoblennius | M |
| *Hypleurochilus bananensis* | Poll, 1959 | Teleostei | Blenniiformes | Blenniidae | Hypleurochilus | M |
| *Lipophrys pholis* | Linnaeus, 1758 | Teleostei | Blenniiformes | Blenniidae | Lipophrys | M |
| *Microlipophrys canevae* | Vinciguerra, 1880 | Teleostei | Blenniiformes | Blenniidae | Microlipophrys | M |
| *Parablennius gattorugine* | Linnaeus, 1758 | Teleostei | Blenniiformes | Blenniidae | Parablennius | M |
| *Parablennius incognitus* | Bath, 1968 | Teleostei | Blenniiformes | Blenniidae | Parablennius | M |
| *Parablennius pilicornis* | Cuvier, 1829 | Teleostei | Blenniiformes | Blenniidae | Parablennius | M |
| *Parablennius sanguinolentus* | Pallas, 1814 | Teleostei | Blenniiformes | Blenniidae | Parablennius | M |
| *Salaria pavo* | Risso, 1810 | Teleostei | Blenniiformes | Blenniidae | Salaria | M / B |
| *Callionymus reticulatus* | Valenciennes, 1837 | Teleostei | Callionymiformes | Callionymidae | Callionymus | M / B |
| *Callionymus risso* | Lesueur, 1814 | Teleostei | Callionymiformes | Callionymidae | Callionymus | M |
| *Caranx hippos* | Linnaeus, 1766 | Teleostei | Carangiformes | Carangidae | Caranx | M / B |
| *Decapterus macrosoma* | Bleeker, 1851 | Teleostei | Carangiformes | Carangidae | Decapterus | M |
| *Trachinotus ovatus* | Linnaeus, 1758 | Teleostei | Carangiformes | Carangidae | Trachinotus | M / B |
| *Trachurus mediterraneus* | Steindachner, 1868 | Teleostei | Carangiformes | Carangidae | Trachurus | M / B |
| *Trachurus picturatus* | Bowdich, 1825 | Teleostei | Carangiformes | Carangidae | Trachurus | M |
| *Trachurus trachurus* | Linnaeus, 1758 | Teleostei | Carangiformes | Carangidae | Trachurus | M |
| *Alosa fallax* | Lacepède, 1803 | Teleostei | Clupeiformes | Alosidae | Alosa | M / B / F |

| Scientific Name | Author | Class | Order ↓ | Family | Genus | Habitat |
|---|---|---|---|---|---|---|
| *Engraulis encrasicolus* | Linnaeus, 1758 | Teleostei | Clupeiformes | Engraulidae | Engraulis | M / B |
| *Sardina pilchardus* | Walbaum, 1792 | Teleostei | Clupeiformes | Alosidae | Sardina | M / B / F |
| *Sardinella aurita* | Valenciennes, 1847 | Teleostei | Clupeiformes | Dorosomatidae | Sardinella | M / B |
| *Acantholabrus palloni* | Risso, 1810 | Teleostei | Eupercaria incertae sedis | Labridae | Acantholabrus | M |
| *Argyrosomus regius* | Asso, 1801 | Teleostei | Eupercaria incertae sedis | Sciaenidae | Argyrosomus | M / B |
| *Boops boops* | Linnaeus, 1758 | Teleostei | Eupercaria incertae sedis | Sparidae | Boops | M |
| *Cepola macrophthalma* | Linnaeus, 1758 | Teleostei | Eupercaria incertae sedis | Cepolidae | Cepola | M |
| *Coris julis* | Linnaeus, 1758 | Teleostei | Eupercaria incertae sedis | Labridae | Coris | M |
| *Cynoscion regalis* | Bloch & Schneider, 1801 | Teleostei | Eupercaria incertae sedis | Sciaenidae | Cynoscion | M / B |
| *Dentex gibbosus* | Rafinesque, 1810 | Teleostei | Eupercaria incertae sedis | Sparidae | Dentex | M |
| *Dicentrarchus labrax* | Linnaeus, 1758 | Teleostei | Eupercaria incertae sedis | Moronidae | Dicentrarchus | M / B / F |
| *Dicentrarchus punctatus* | Bloch, 1792 | Teleostei | Eupercaria incertae sedis | Moronidae | Dicentrarchus | M / B |
| *Diplodus annularis* | Linnaeus, 1758 | Teleostei | Eupercaria incertae sedis | Sparidae | Diplodus | M / B |
| *Diplodus bellottii* | Steindachner, 1882 | Teleostei | Eupercaria incertae sedis | Sparidae | Diplodus | M |
| *Diplodus sargus* | Linnaeus, 1758 | Teleostei | Eupercaria incertae sedis | Sparidae | Diplodus | M / B |
| *Diplodus vulgaris* | Geoffroy Saint-Hilaire, 1817 | Teleostei | Eupercaria incertae sedis | Sparidae | Diplodus | M |
| *Labrus bergylta* | Ascanius, 1767 | Teleostei | Eupercaria incertae sedis | Labridae | Labrus | M |
| *Lithognathus mormyrus* | Linnaeus, 1758 | Teleostei | Eupercaria incertae sedis | Sparidae | Lithognathus | M / B |
| *Pagellus acarne* | Risso, 1827 | Teleostei | Eupercaria incertae sedis | Sparidae | Pagellus | M |
| *Pagellus bogaraveo* | Brünnich, 1768 | Teleostei | Eupercaria incertae sedis | Sparidae | Pagellus | M |
| *Pagellus erythrinus* | Linnaeus, 1758 | Teleostei | Eupercaria incertae sedis | Sparidae | Pagellus | M |
| *Pagrus pagrus* | Linnaeus, 1758 | Teleostei | Eupercaria incertae sedis | Sparidae | Pagrus | M |
| *Parapristipoma trilineatum* | Thunberg, 1793 | Teleostei | Eupercaria incertae sedis | Haemulidae | Parapristipoma | M |
| *Pomadasys incisus* | Bowdich, 1825 | Teleostei | Eupercaria incertae sedis | Haemulidae | Pomadasys | M / B |
| *Sarpa salpa* | Linnaeus, 1758 | Teleostei | Eupercaria incertae sedis | Sparidae | Sarpa | M / B |
| *Sparus aurata* | Linnaeus, 1758 | Teleostei | Eupercaria incertae sedis | Sparidae | Sparus | M / B |
| *Spicara maena* | Linnaeus, 1758 | Teleostei | Eupercaria incertae sedis | Sparidae | Spicara | M |
| *Spondyliosoma cantharus* | Linnaeus, 1758 | Teleostei | Eupercaria incertae sedis | Sparidae | Spondyliosoma | M |
| *Symphodus bailloni* | Valenciennes, 1839 | Teleostei | Eupercaria incertae sedis | Labridae | Symphodus | M |
| *Symphodus melops* | Linnaeus, 1758 | Teleostei | Eupercaria incertae sedis | Labridae | Symphodus | M |

| Scientific Name | Author | Class | Order ↓ | Family | Genus | Habitat |
|---|---|---|---|---|---|---|
| *Symphodus ocellatus* | Linnaeus, 1758 | Teleostei | Eupercaria incertae sedis | Labridae | Symphodus | M |
| *Umbrina canariensis* | Valenciennes, 1843 | Teleostei | Eupercaria incertae sedis | Sciaenidae | Umbrina | M |
| *Gadiculus argenteus* | Guichenot, 1850 | Teleostei | Gadiformes | Gadidae | Gadiculus | M |
| *Gaidropsarus mediterraneus* | Linnaeus, 1758 | Teleostei | Gadiformes | Gaidropsaridae | Gaidropsarus | M / B |
| *Merluccius merluccius* | Linnaeus, 1758 | Teleostei | Gadiformes | Merlucciidae | Merluccius | M |
| *Micromesistius poutassou* | Risso, 1827 | Teleostei | Gadiformes | Gadidae | Micromesistius | M |
| *Trisopterus luscus* | Linnaeus, 1758 | Teleostei | Gadiformes | Gadidae | Trisopterus | M / B |
| *Diplecogaster bimaculata* | Bonnaterre, 1788 | Teleostei | Gobiesociformes | Gobiesocidae | Diplecogaster | M |
| *Opeatogenys gracilis* | Canestrini, 1864 | Teleostei | Gobiesociformes | Gobiesocidae | Opeatogenys | M |
| *Chromogobius zebratus* | Kolombatovic, 1891 | Teleostei | Gobiiformes | Gobiidae | Chromogobius | M |
| *Gobius couchi* | Miller & El-Tawil, 1974 | Teleostei | Gobiiformes | Gobiidae | Gobius | M / B |
| *Gobius niger* | Linnaeus, 1758 | Teleostei | Gobiiformes | Gobiidae | Gobius | M / B |
| *Gobius paganellus* | Linnaeus, 1758 | Teleostei | Gobiiformes | Gobiidae | Gobius | M / B / F |
| *Pomatoschistus microps* | Krøyer, 1838 | Teleostei | Gobiiformes | Gobiidae | Pomatoschistus | M / B / F |
| *Pomatoschistus norvegicus* | Collett, 1902 | Teleostei | Gobiiformes | Gobiidae | Pomatoschistus | M |
| *Pomatoschistus pictus* | Malm, 1865 | Teleostei | Gobiiformes | Gobiidae | Pomatoschistus | M |
| *Pseudaphya ferreri* | de Buen & Fage, 1908 | Teleostei | Gobiiformes | Gobiidae | Pseudaphya | M |
| *Sicyopterus eudentatus* | Parenti & Maciolek, 1993 | Teleostei | Gobiiformes | Gobiidae | Sicyopterus | M / B / F |
| *Zebrus zebrus* | Risso, 1827 | Teleostei | Gobiiformes | Gobiidae | Zebrus | M |
| *Lophius budegassa* | Spinola, 1807 | Teleostei | Lophiiformes | Lophiidae | Lophius | M |
| *Lophius piscatorius* | Linnaeus, 1758 | Teleostei | Lophiiformes | Lophiidae | Lophius | M |
| *Chelon auratus* | Risso, 1810 | Teleostei | Mugiliformes | Mugilidae | Chelon | M / B / F |
| *Chelon labrosus* | Risso, 1827 | Teleostei | Mugiliformes | Mugilidae | Chelon | M / B / F |
| *Chelon ramada* | Risso, 1827 | Teleostei | Mugiliformes | Mugilidae | Chelon | M / B / F |
| *Chelon saliens* | Risso, 1810 | Teleostei | Mugiliformes | Mugilidae | Chelon | M / B |
| *Mugil cephalus* | Linnaeus, 1758 | Teleostei | Mugiliformes | Mugilidae | Mugil | M / B / F |
| *Mullus barbatus* | Linnaeus, 1758 | Teleostei | Mulliformes | Mullidae | Mullus | M |
| *Mullus surmuletus* | Linnaeus, 1758 | Teleostei | Mulliformes | Mullidae | Mullus | M |
| *Myctophum punctatum* | Rafinesque, 1810 | Teleostei | Myctophiformes | Myctophidae | Myctophum | M |

| Scientific Name | Author | Class | Order ↓ | Family | Genus | Habitat |
|---|---|---|---|---|---|---|
| *Abudefduf saxatilis* | Linnaeus, 1758 | Teleostei | Ovalentaria incertae sedis | Pomacentridae | Abudefduf | M |
| *Anthias anthias* | Linnaeus, 1758 | Teleostei | Perciformes | Anthiadidae | Anthias | M |
| *Chelidonichthys lucerna* | Linnaeus, 1758 | Teleostei | Perciformes | Triglidae | Chelidonichthys | M |
| *Chelidonichthys obscurus* | Walbaum, 1792 | Teleostei | Perciformes | Triglidae | Chelidonichthys | M |
| *Echiichthys vipera* | Cuvier, 1829 | Teleostei | Perciformes | Trachinidae | Echiichthys | M |
| *Helicolenus dactylopterus* | Delaroche, 1809 | Teleostei | Perciformes | Sebastidae | Helicolenus | M |
| *Lepidotrigla cavillone* | Lacepède, 1801 | Teleostei | Perciformes | Triglidae | Lepidotrigla | M |
| *Scorpaena notata* | Rafinesque, 1810 | Teleostei | Perciformes | Scorpaenidae | Scorpaena | M |
| *Serranus cabrilla* | Linnaeus, 1758 | Teleostei | Perciformes | Serranidae | Serranus | M |
| *Serranus hepatus* | Linnaeus, 1758 | Teleostei | Perciformes | Serranidae | Serranus | M |
| *Arnoglossus imperialis* | Rafinesque, 1810 | Teleostei | Pleuronectiformes | Bothidae | Arnoglossus | M |
| *Arnoglossus laterna* | Walbaum, 1792 | Teleostei | Pleuronectiformes | Bothidae | Arnoglossus | M |
| *Arnoglossus thori* | Kyle, 1913 | Teleostei | Pleuronectiformes | Bothidae | Arnoglossus | M |
| *Buglossidium luteum* | Risso, 1810 | Teleostei | Pleuronectiformes | Soleidae | Buglossidium | M |
| *Citharus linguatula* | Linnaeus, 1758 | Teleostei | Pleuronectiformes | Citharidae | Citharus | M |
| *Dagetichthys lusitanicus* | de Brito Capello, 1868 | Teleostei | Pleuronectiformes | Soleidae | Dagetichthys | M / B |
| *Dicologlossa cuneata* | Moreau, 1881 | Teleostei | Pleuronectiformes | Soleidae | Dicologlossa | M / B |
| *Microchirus boscanion* | Chabanaud, 1926 | Teleostei | Pleuronectiformes | Soleidae | Microchirus | M |
| *Pegusa impar* | Bennett, 1831 | Teleostei | Pleuronectiformes | Soleidae | Pegusa | M |
| *Pegusa lascaris* | Risso, 1810 | Teleostei | Pleuronectiformes | Soleidae | Pegusa | M / B |
| *Scophthalmus rhombus* | Linnaeus, 1758 | Teleostei | Pleuronectiformes | Scophthalmidae | Scophthalmus | M |
| *Solea senegalensis* | Kaup, 1858 | Teleostei | Pleuronectiformes | Soleidae | Solea | M |
| *Solea solea* | Linnaeus, 1758 | Teleostei | Pleuronectiformes | Soleidae | Solea | M / B |
| *Oncorhynchus mykiss* | Walbaum, 1792 | Teleostei | Salmoniformes | Salmonidae | Oncorhynchus | M / B / F |
| *Salmo trutta* | Linnaeus, 1758 | Teleostei | Salmoniformes | Salmonidae | Salmo | M / B / F |
| *Auxis rochei* | Risso, 1810 | Teleostei | Scombriformes | Scombridae | Auxis | M / B |
| *Euthynnus alletteratus* | Rafinesque, 1810 | Teleostei | Scombriformes | Scombridae | Euthynnus | M / B |
| *Katsuwonus pelamis* | Linnaeus, 1758 | Teleostei | Scombriformes | Scombridae | Katsuwonus | M |
| *Pomatomus saltatrix* | Linnaeus, 1766 | Teleostei | Scombriformes | Pomatomidae | Pomatomus | M / B |
| *Scomber colias* | Gmelin, 1789 | Teleostei | Scombriformes | Scombridae | Scomber | M / B |

| Scientific Name | Author | Class | Order ↓ | Family | Genus | Habitat |
|---|---|---|---|---|---|---|
| *Scomber scombrus* | Linnaeus, 1758 | Teleostei | Scombriformes | Scombridae | Scomber | M / B |
| *Stromateus fiatola* | Linnaeus, 1758 | Teleostei | Scombriformes | Stromateidae | Stromateus | M |
| *Macroramphosus scolopax* | Linnaeus, 1758 | Teleostei | Syngnathiformes | Centriscidae | Macroramphosus | M |
| *Nerophis ophidion* | Linnaeus, 1758 | Teleostei | Syngnathiformes | Syngnathidae | Nerophis | M / B / F |
| *Syngnathus rostellatus* | Nilsson, 1855 | Teleostei | Syngnathiformes | Syngnathidae | Syngnathus | M / B |
| *Syngnathus typhle* | Linnaeus, 1758 | Teleostei | Syngnathiformes | Syngnathidae | Syngnathus | M / B |
| *Balistes capriscus* | Gmelin, 1789 | Teleostei | Tetraodontiformes | Balistidae | Balistes | M |
| *Zeus faber* | Linnaeus, 1758 | Teleostei | Zeiformes | Zeidae | Zeus | M / B |
| *Galeorhinus galeus* | Linnaeus, 1758 | Elasmobranchii | Carcharhiniformes | Triakidae | Galeorhinus | M |
| *Cetorhinus maximus* | Gunnerus, 1765 | Elasmobranchii | Lamniformes | Cetorhinidae | Cetorhinus | M |
| *Himantura uarnak* | Gmelin, 1789 | Elasmobranchii | Myliobatiformes | Dasyatidae | Himantura | M / B |
| *Mobula mobular* | Bonnaterre, 1788 | Elasmobranchii | Myliobatiformes | Mobulidae | Mobula | M |
| *Dipturus nidarosiensis* | Storm, 1881 | Elasmobranchii | Rajiformes | Rajidae | Dipturus | M |
| *Raja brachyura* | Lafont, 1871 | Elasmobranchii | Rajiformes | Rajidae | Raja | M |
| *Raja montagui* | Fowler, 1910 | Elasmobranchii | Rajiformes | Rajidae | Raja | M |
| *Raja undulata* | Lacepède, 1802 | Elasmobranchii | Rajiformes | Rajidae | Raja | M |
| *Scymnodon ringens* | Bocage & Capello, 1864 | Elasmobranchii | Squaliformes | Somniosidae | Scymnodon | M |
| *Torpedo marmorata* | Risso, 1810 | Elasmobranchii | Torpediniformes | Torpedinidae | Torpedo | M / B |

**Supplementary Table 9.** Fish species identified through ichthyoplankton samples (DNA metabarcoding and morphology – green pattern), water samples (eDNA – blue pattern), and both sampling methods combined ( purple pattern) over the 13-month study period. Table is subdivided for each category associated with the number of monthly occurrences registered for each species: **a)** Very Frequent – present in 12-13 months; **b)** Frequent – present in 8-11 months; **c)** Moderately Frequent – present in 4-7 months; **d)** Rare – present in 2-3 months; **e)** Very Rare – present in 1 month, and **f)** specific table for the Elasmobranchii records. Species are organized alphabetically by order name (↓). (Spr.) denotes Spring; (*) indicates *incertae sedis*.

| **Order (↓)** | **Species** | **Spring** | | | **Summer** | | | **Autumn** | | | **Winter** | | | **Spr.** |
|---|---|---|---|---|---|---|---|---|---|---|---|---|---|---|
| | | **APR** | **MAY** | **JUN** | **JUL** | **AUG** | **SEP** | **OCT** | **NOV** | **DEC** | **JAN** | **FEB** | **MAR** | **APR*** |
| **a)** *Monthly records for "Very Frequent" fish species (present in 12 - 13 months)* | | | | | | | | | | | | | | |
| Atheriniformes | *Atherina presbyter* | Purple | Green | Purple | Purple | Green | Blue | Green | Purple | Green | Purple | Green | Purple | Green |
| Clupeiformes | *Engraulis encrasicolus* | Purple | Purple | Purple | Purple | Purple | Purple | Purple | Purple | Purple | Purple | Green | Purple | Purple |
| Clupeiformes | *Sardina pilchardus* | Purple | Purple | Purple | Purple | Purple | Purple | Purple | Purple | Purple | Purple | Green | Purple | Purple |
| Eupercaria* | *Dicentrarchus labrax* | Purple | Purple | Purple | Blue | Blue | Purple | Purple | Blue | Green | Purple | | Purple | Purple |
| Eupercaria* | *Diplodus bellottii* | Purple | Purple | Purple | Purple | Purple | Purple | Green | Blue | | Purple | Green | Purple | Purple |
| Eupercaria* | *Diplodus sargus* | Purple | Purple | Blue | Purple | Purple | Purple | Purple | Purple | Green | Purple | Green | Green | Green |
| Eupercaria* | *Sparus aurata* | Purple | Purple | Purple | Purple | Purple | Purple | Purple | Purple | Purple | Purple | Green | Purple | Blue |
| Gobiiformes | *Pomatoschistus microps* | Purple | Purple | Purple | Purple | Purple | Purple | Green | Purple | Purple | Purple | Green | Purple | Purple |
| Mugiliformes | *Chelon labrosus* | Purple | Purple | Purple | Purple | Purple | Purple | Blue | Purple | Purple | Purple | Green | Purple | Purple |
| Mugiliformes | *Mugil cephalus* | Purple | Blue | Purple | Purple | Blue | Purple | Purple | Purple | Green | Green | Green | Green | Purple |
| Pleuronectiformes | *Solea senegalensis* | Green | Green | Blue | Purple | Green | Purple | Purple | Green | Purple | Purple | Green | Green | Green |
| Syngnathiformes | *Syngnathus rostellatus* | Green | | Green | Blue | Purple | Blue | Green | Green | Green | Purple | Green | Green | Green |
| **b)** *Monthly records for "Frequent" fish species (present in 8 - 11 months)* | | | | | | | | | | | | | | |
| Beloniformes | *Belone belone* | Purple | Blue | Purple | Blue | | Purple | Green | Blue | | Blue | | | Purple |
| Blenniiformes | *Parablennius pilicornis* | Purple | Green | Green | Purple | Green | Green | | Purple | | | Green | Purple | Green |
| Carangiformes | *Trachurus trachurus* | Blue | Blue | Purple | Green | Green | | Green | Green | | Green | Green | Green | Green |
| Eupercaria* | *Boops boops* | Green | Purple | Purple | | | | Green | | | Green | Green | Green | Purple |
| Eupercaria* | *Coris julis* | | Blue | Purple | Purple | Purple | | Green | Green | Blue | | | | |
| Eupercaria* | *Dicentrarchus punctatus* | Blue | Green | | Blue | | | | Purple | | Blue | Green | Blue | Purple |
| Eupercaria* | *Diplodus vulgaris* | Blue | | | | Blue | Green | | | Green | Green | Green | Green | Green |
| Eupercaria* | *Pagellus erythrinus* | | Green | Purple | Green | Purple | Green | | Green | Green | Purple | | | Green |
| Gobiiformes | *Gobius niger* | Purple | Purple | Green | | Purple | Purple | Blue | Blue | | | Green | Green | Green |
| Gobiiformes | *Pomatoschistus pictus* | Purple | Green | Green | Green | Green | | Green | Green | | | Green | | Green |
| Mugiliformes | *Chelon ramada* | Blue | Blue | | Green | | Green | | Purple | Green | Green | Green | Blue | Green |
| Mugiliformes | *Chelon saliens* | Blue | Purple | Purple | | Blue | Purple | Blue | Blue | | Green | | Blue | |
| Perciformes | *Serranus hepatus* | Blue | Purple | Blue | Green | Green | Green | Green | Green | Green | Green | | | Green |
| Pleuronectiformes | *Arnoglossus imperialis* | | Purple | Green | Green | Green | | | | | | Green | Green | Green |
| Pleuronectiformes | *Arnoglossus thori* | Green | Green | Purple | Green | Purple | Green | Green | Green | | | | Green | Green |
| Pleuronectiformes | *Dicologlossa cuneata* | Green | | Green | Blue | | | Green | Green | | Green | Green | | Green |

| Order (↓) | Species | Spring | | | Summer | | | Autumn | | | Winter | | | Spr. |
|---|---|---|---|---|---|---|---|---|---|---|---|---|---|---|
| | | APR | MAY | JUN | JUL | AUG | SEP | OCT | NOV | DEC | JAN | FEB | MAR | APR* |
| Pleuronectiformes | *Pegusa lascaris* | | | | | | | | | | | | | |
| **c)** *Monthly records for "Moderately Frequent " fish species (present in 4 - 7 months)* | | | | | | | | | | | | | | |
| Blenniiformes | *Coryphoblennius galerita* | | | | | | | | | | | | | |
| Blenniiformes | *Hypleurochilus bananensis* | | | | | | | | | | | | | |
| Blenniiformes | *Parablennius gattorugine* | | | | | | | | | | | | | |
| Blenniiformes | *Salaria pavo* | | | | | | | | | | | | | |
| Callionymiformes | *Callionymus risso* | | | | | | | | | | | | | |
| Clupeiformes | *Alosa fallax* | | | | | | | | | | | | | |
| Clupeiformes | *Sardinella aurita* | | | | | | | | | | | | | |
| Eupercaria* | *Argyrosomus regius* | | | | | | | | | | | | | |
| Eupercaria* | *Cynoscion regalis* | | | | | | | | | | | | | |
| Eupercaria* | *Diplodus annularis* | | | | | | | | | | | | | |
| Eupercaria* | *Lithognathus mormyrus* | | | | | | | | | | | | | |
| Eupercaria* | *Pagellus acarne* | | | | | | | | | | | | | |
| Eupercaria* | *Sarpa salpa* | | | | | | | | | | | | | |
| Eupercaria* | *Spicara maena* | | | | | | | | | | | | | |
| Gadiformes | *Merluccius merluccius* | | | | | | | | | | | | | |
| Gobiiformes | *Gobius paganellus* | | | | | | | | | | | | | |
| Gobiiformes | *Pseudaphya ferreri* | | | | | | | | | | | | | |
| Mulliformes | *Mullus barbatus* | | | | | | | | | | | | | |
| Mulliformes | *Mullus surmuletus* | | | | | | | | | | | | | |
| Perciformes | *Serranus cabrilla* | | | | | | | | | | | | | |
| Pleuronectiformes | *Arnoglossus laterna* | | | | | | | | | | | | | |
| Pleuronectiformes | *Buglossidium luteum* | | | | | | | | | | | | | |
| Pleuronectiformes | *Dagetichthys lusitanicus* | | | | | | | | | | | | | |
| Scombriformes | *Pomatomus saltatrix* | | | | | | | | | | | | | |
| Scombriformes | *Scomber colias* | | | | | | | | | | | | | |
| **d)** *Monthly records for "Rare" fish species (present in 2 - 3 months)* | | | | | | | | | | | | | | |
| Anguilliformes | *Conger conger* | | | | | | | | | | | | | |
| Beloniformes | *Belone svetovidovi* | | | | | | | | | | | | | |
| Blenniiformes | *Lipophrys pholis* | | | | | | | | | | | | | |
| Blenniiformes | *Parablennius incognitus* | | | | | | | | | | | | | |
| Carangiformes | *Trachinotus ovatus* | | | | | | | | | | | | | |
| Carangiformes | *Trachurus mediterraneus* | | | | | | | | | | | | | |
| Eupercaria* | *Pagrus pagrus* | | | | | | | | | | | | | |
| Eupercaria* | *Spondyliosoma cantharus* | | | | | | | | | | | | | |
| Eupercaria* | *Symphodus bailloni* | | | | | | | | | | | | | |
| Eupercaria* | *Symphodus melops* | | | | | | | | | | | | | |
| Eupercaria* | *Umbrina canariensis* | | | | | | | | | | | | | |
| Gadiformes | *Gadiculus argenteus* | | | | | | | | | | | | | |
| Gadiformes | *Micromesistius poutassou* | | | | | | | | | | | | | |

| Order (↓) | Species | Spring | | | Summer | | | Autumn | | | Winter | | | Spr. |
|---|---|---|---|---|---|---|---|---|---|---|---|---|---|---|
| | | APR | MAY | JUN | JUL | AUG | SEP | OCT | NOV | DEC | JAN | FEB | MAR | APR* |
| Gobiesociformes | *Opeatogenys gracilis* | | | | ■ | ■ | | | | | | | | |
| Gobiiformes | *Gobius couchi* | | | | | | ▥ | | | | | | ■ | ■ |
| Gobiiformes | *Zebrus zebrus* | | | | ■ | ■ | | | | | | | | |
| Mugiliformes | *Chelon auratus* | | ■ | | | | | | | ■ | | ■ | | |
| Myctophiformes | *Myctophum punctatum* | | ▨ | ▥ | | | | | ■ | | | | | |
| Perciformes | *Anthias anthias* | | | | | | | | ■ | ■ | | | | |
| Perciformes | *Chelidonichthys lucerna* | | | | | | | | | | | ■ | | ■ |
| Perciformes | *Echiichthys vipera* | | ■ | | | | ■ | | | | | | | ■ |
| Perciformes | *Helicolenus dactylopterus* | | | | | ■ | | | | | ■ | | | |
| Perciformes | *Scorpaena notata* | ■ | ▥ | | | | | ■ | | | | | | |
| Pleuronectiformes | *Microchirus boscanion* | | ■ | | | | | | | | | | | ■ |
| Pleuronectiformes | *Pegusa impar* | | | | | | | | ■ | | ■ | ■ | | |
| Pleuronectiformes | *Solea solea* | | | | ■ | | | | | | ■ | | | |
| Syngnathiformes | *Syngnathus typhle* | | | ■ | | ■ | | | | | | | | |
| Tetraodontiformes | *Balistes capriscus* | | | | | ■ | | | | | | | | |

**e)** *Monthly records for "Very Rare" fish species (present in 1 month)*

| Order | Species | APR | MAY | JUN | JUL | AUG | SEP | OCT | NOV | DEC | JAN | FEB | MAR | APR* |
|---|---|---|---|---|---|---|---|---|---|---|---|---|---|---|
| Acanthuriformes | *Capros aper* | ▥ | | | | | | | | | | | | |
| Beloniformes | *Scomberesox saurus* | | | | | | | | | ■ | | | | |
| Blenniiformes | *Microlipophrys canevae* | | | ■ | | | | | | | | | | |
| Blenniiformes | *Parablennius sanguinolentus* | | ■ | | | | | | | | | | | |
| Callionymiformes | *Callionymus reticulatus* | | | | | | | | | | | | | ■ |
| Carangiformes | *Caranx hippos* | | | | | | ■ | | | | | | | |
| Carangiformes | *Decapterus macrosoma* | ■ | | | | | | | | | | | | |
| Carangiformes | *Trachurus picturatus* | ▥ | | | | | | | | | | | | |
| Eupercaria* | *Acantholabrus palloni* | | | | | | | | | | | | | ■ |
| Eupercaria* | *Cepola macrophthalma* | | ■ | | | | | | | | | | | |
| Eupercaria* | *Dentex gibbosus* | | | | | | | | | | ■ | | | |
| Eupercaria* | *Labrus bergylta* | | | | | | | | | | | | ■ | |
| Eupercaria* | *Pagellus bogaraveo* | | | ▥ | | | | | | | | | | |
| Eupercaria* | *Parapristipoma trilineatum* | | ■ | | | | | | | | | | | |
| Eupercaria* | *Pomadasys incisus* | | | | ■ | | | | | | | | | |
| Eupercaria* | *Symphodus ocellatus* | | | ■ | | | | | | | | | | |
| Gadiformes | *Gaidropsarus mediterraneus* | | | | | | | | | | | ■ | | |
| Gadiformes | *Trisopterus luscus* | | | | | | | | | | | ■ | | |
| Gobiesociformes | *Diplecogaster bimaculata* | | | | ■ | | | | | | | | | |
| Gobiiformes | *Chromogobius zebratus* | | | | | | | | | ■ | | | | |
| Gobiiformes | *Pomatoschistus norvegicus* | | | | | | | | | | | ■ | | |
| Gobiiformes | *Sicyopterus eudentatus* | | ■ | | | | | | | | | | | |
| Lophiiformes | *Lophius budegassa* | ■ | | | | | | | | | | | | |
| Lophiiformes | *Lophius piscatorius* | | | | | | | ▥ | | | | | | |
| Ovalentaria* | *Abudefduf saxatilis* | | | | | | | | | ■ | | | | |
| Perciformes | *Chelidonichthys obscurus* | | ▨ | | | | | | | | | | | |

| Order (↓) | Species | Spring | | | Summer | | | Autumn | | | Winter | | | Spr. |
|---|---|---|---|---|---|---|---|---|---|---|---|---|---|---|
| | | APR | MAY | JUN | JUL | AUG | SEP | OCT | NOV | DEC | JAN | FEB | MAR | APR* |
| Perciformes | *Lepidotrigla cavillone* | | | | | | | | | | | | | ■ |
| Pleuronectiformes | *Citharus linguatula* | ▥ | | | | | | | | | | | | |
| Pleuronectiformes | *Scophthalmus rhombus* | | | | | | | | | | ■ | | | |
| Salmoniformes | *Oncorhynchus mykiss* | | | | | | | | | | | ■ | | |
| Salmoniformes | *Salmo trutta* | | | | | | | | | | | | ■ | |
| Scombriformes | *Auxis rochei* | ■ | | | | | | | | | | | | |
| Scombriformes | *Euthynnus alletteratus* | | | | | | | | | ■ | | | | |
| Scombriformes | *Katsuwonus pelamis* | ■ | | | | | | | | | | | | |
| Scombriformes | *Scomber scombrus* | ▥ | | | | | | | | | | | | |
| Scombriformes | *Stromateus fiatola* | | | | | | ▥ | | | | | | | |
| Syngnathiformes | *Macroramphosus scolopax* | | | | | | | | | | | ■ | | |
| Syngnathiformes | *Nerophis ophidion* | | ■ | | | | | | | | | | | |
| Zeiformes | *Zeus faber* | | ■ | | | | | | | | | | | |
| **f)** *Monthly records for Elasmobranchii class* | | | | | | | | | | | | | | |
| Carcharhiniformes | *Galeorhinus galeus* | ▥ | | | | | | | | | | | | |
| Lamniformes | *Cetorhinus maximus* | ▥ | | | | | | | | | | | | |
| Myliobatiformes | *Himantura uarnak* | | | | | | | | | ■ | | | | |
| Myliobatiformes | *Mobula mobular* | | | | ■ | | | | | | | | | |
| Rajiformes | *Dipturus nidarosiensis* | ■ | | | | | | | | | | | | |
| Rajiformes | *Raja brachyura* | | | | | | | | | | ■ | | | |
| Rajiformes | *Raja montagui* | | | | | | | | | | ■ | | | |
| Rajiformes | *Raja undulata* | ▥ | ▥ | ▥ | | | ▥ | | | | | | | |
| Squaliformes | *Scymnodon ringens* | ■ | | | | | | | | | | | | |
| Torpediniformes | *Torpedo marmorata* | ▥ | | | | | | | | | | | | |

**Supplementary Table 10**. List of fish species identified in each season during the 13-month study in the lower section of the Guadiana River Estuary.

| Spring (n=99) | Summer (n=71) | Autumn (n=56) | Winter (n=61) |
|---|---|---|---|
| *Acantholabrus palloni* | *Alosa fallax* | *Atherina presbyter* | *Arnoglossus laterna* |
| *Alosa fallax* | *Argyrosomus regius* | *Abudefduf saxatilis* | *Alosa fallax* |
| *Argyrosomus regius* | *Arnoglossus imperialis* | *Alosa fallax* | *Argyrosomus regius* |
| *Arnoglossus imperialis* | *Arnoglossus thori* | *Anthias anthias* | *Arnoglossus imperialis* |
| *Arnoglossus laterna* | *Atherina presbyter* | *Arnoglossus laterna* | *Arnoglossus thori* |
| *Arnoglossus thori* | *Balistes capriscus* | *Arnoglossus thori* | *Atherina presbyter* |
| *Atherina presbyter* | *Belone belone* | *Belone belone* | *Belone belone* |
| *Auxis rochei* | *Belone svetovidovi* | *Boops boops* | *Boops boops* |
| *Belone belone* | *Callionymus risso* | *Chelon auratus* | *Buglossidium luteum* |
| *Belone svetovidovi* | *Caranx hippos* | *Chelon labrosus* | *Callionymus risso* |
| *Boops boops* | *Chelon labrosus* | *Chelon ramada* | *Chelidonichthys lucerna* |
| *Buglossidium luteum* | *Chelon ramada* | *Chelon saliens* | *Chelon auratus* |
| *Callionymus reticulatus* | *Chelon saliens* | *Chromogobius zebratus* | *Chelon labrosus* |
| *Callionymus risso* | *Conger conger* | *Coris julis* | *Chelon ramada* |
| *Capros aper* | *Coris julis* | *Coryphoblennius galerita* | *Chelon saliens* |
| *Cepola macrophthalma* | *Coryphoblennius galerita* | *Dicentrarchus labrax* | *Coryphoblennius galerita* |
| *Cetorhinus maximus* | *Cynoscion regalis* | *Dicentrarchus punctatus* | *Dagetichthys lusitanicus* |
| *Chelidonichthys lucerna* | *Dagetichthys lusitanicus* | *Dicologlossa cuneata* | *Dentex gibbosus* |
| *Chelidonichthys obscurus* | *Dicentrarchus labrax* | *Diplodus annularis* | *Dicentrarchus labrax* |
| *Chelon auratus* | *Dicentrarchus punctatus* | *Diplodus bellottii* | *Dicentrarchus punctatus* |
| *Chelon labrosus* | *Dicologlossa cuneata* | *Diplodus sargus* | *Dicologlossa cuneata* |
| *Chelon ramada* | *Diplecogaster bimaculata* | *Diplodus vulgaris* | *Diplodus annularis* |
| *Chelon saliens* | *Diplodus annularis* | *Engraulis encrasicolus* | *Diplodus bellottii* |
| *Citharus linguatula* | *Diplodus bellottii* | *Euthynnus alletteratus* | *Diplodus sargus* |
| *Conger conger* | *Diplodus sargus* | *Gadiculus argenteus* | *Diplodus vulgaris* |
| *Coris julis* | *Diplodus vulgaris* | *Gobius niger* | *Engraulis encrasicolus* |
| *Coryphoblennius galerita* | *Echiichthys vipera* | *Himantura uarnak* | *Gaidropsarus mediterraneus* |
| *Cynoscion regalis* | *Engraulis encrasicolus* | *Hypleurochilus bananensis* | *Gobius couchi* |
| *Dagetichthys lusitanicus* | *Gobius couchi* | *Lithognathus mormyrus* | *Gobius niger* |
| *Decapterus macrosoma* | *Gobius niger* | *Lophius piscatorius* | *Gobius paganellus* |
| *Dicentrarchus labrax* | *Gobius paganellus* | *Merluccius merluccius* | *Helicolenus dactylopterus* |
| *Dicentrarchus punctatus* | *Helicolenus dactylopterus* | *Mugil cephalus* | *Hypleurochilus bananensis* |
| *Dicologlossa cuneata* | *Lithognathus mormyrus* | *Mullus surmuletus* | *Labrus bergylta* |
| *Diplodus annularis* | *Micromesistius poutassou* | *Myctophum punctatum* | *Macroramphosus scolopax* |
| *Diplodus bellottii* | *Mobula mobular* | *Pagellus acarne* | *Merluccius merluccius* |
| *Diplodus sargus* | *Mugil cephalus* | *Pagellus erythrinus* | *Mugil cephalus* |
| *Diplodus vulgaris* | *Mullus barbatus* | *Parablennius gattorugine* | *Oncorhynchus mykiss* |
| *Dipturus nidarosiensis* | *Mullus surmuletus* | *Parablennius pilicornis* | *Pagellus acarne* |
| *Echiichthys vipera* | *Opeatogenys gracilis* | *Pegusa impar* | *Pagellus erythrinus* |
| *Engraulis encrasicolus* | *Pagellus acarne* | *Pegusa lascaris* | *Parablennius gattorugine* |
| *Gadiculus argenteus* | *Pagellus erythrinus* | *Pomatoschistus microps* | *Parablennius pilicornis* |
| *Galeorhinus galeus* | *Pagrus pagrus* | *Pomatoschistus pictus* | *Pegusa impar* |

| Spring (n=99) | Summer (n=71) | Autumn (n=56) | Winter (n=61) |
|---|---|---|---|
| *Gobius couchi* | *Parablennius gattorugine* | *Pseudaphya ferreri* | *Pegusa lascaris* |
| *Gobius niger* | *Parablennius incognitus* | *Salaria pavo* | *Pomatoschistus microps* |
| *Gobius paganellus* | *Parablennius pilicornis* | *Sardina pilchardus* | *Pomatoschistus norvegicus* |
| *Hypleurochilus bananensis* | *Pegusa lascaris* | *Sardinella aurita* | *Pomatoschistus pictus* |
| *Katsuwonus pelamis* | *Pomadasys incisus* | *Sarpa salpa* | *Pseudaphya ferreri* |
| *Lepidotrigla cavillone* | *Pomatomus saltatrix* | *Scomberesox saurus* | *Raja brachyura* |
| *Lipophrys pholis* | *Pomatoschistus microps* | *Scorpaena notata* | *Raja montagui* |
| *Lophius budegassa* | *Pomatoschistus pictus* | *Serranus cabrilla* | *Salmo trutta* |
| *Merluccius merluccius* | *Pseudaphya ferreri* | *Serranus hepatus* | *Sardina pilchardus* |
| *Microchirus boscanion* | *Raja undulata* | *Solea senegalensis* | *Sarpa salpa* |
| *Microlipophrys canevae* | *Salaria pavo* | *Sparus aurata* | *Scophthalmus rhombus* |
| *Micromesistius poutassou* | *Sardina pilchardus* | *Syngnathus rostellatus* | *Serranus hepatus* |
| *Mugil cephalus* | *Sardinella aurita* | *Trachinotus ovatus* | *Solea senegalensis* |
| *Mullus barbatus* | *Sarpa salpa* | *Trachurus trachurus* | *Solea solea* |
| *Mullus surmuletus* | *Scomber colias* | | *Sparus aurata* |
| *Myctophum punctatum* | *Serranus cabrilla* | | *Symphodus melops* |
| *Nerophis ophidion* | *Serranus hepatus* | | *Syngnathus rostellatus* |
| *Pagellus acarne* | *Solea senegalensis* | | *Trachurus trachurus* |
| *Pagellus bogaraveo* | *Solea solea* | | *Trisopterus luscus* |
| *Pagellus erythrinus* | *Sparus aurata* | | |
| *Pagrus pagrus* | *Spicara maena* | | |
| *Parablennius gattorugine* | *Stromateus fiatola* | | |
| *Parablennius incognitus* | *Syngnathus rostellatus* | | |
| *Parablennius pilicornis* | *Syngnathus typhle* | | |
| *Parablennius sanguinolentus* | *Trachinotus ovatus* | | |
| *Parapristipoma trilineatum* | *Trachurus mediterraneus* | | |
| *Pegusa lascaris* | *Trachurus trachurus* | | |
| *Pomatomus saltatrix* | *Umbrina canariensis* | | |
| *Pomatoschistus microps* | *Zebrus zebrus* | | |
| *Pomatoschistus pictus* | | | |
| *Pseudaphya ferreri* | | | |
| *Raja undulata* | | | |
| *Salaria pavo* | | | |
| *Sardina pilchardus* | | | |
| *Sardinella aurita* | | | |
| *Sarpa salpa* | | | |
| *Scomber colias* | | | |
| *Scomber scombrus* | | | |
| *Scorpaena notata* | | | |
| *Scymnodon ringens* | | | |
| *Serranus cabrilla* | | | |
| *Serranus hepatus* | | | |
| *Sicyopterus eudentatus* | | | |
| *Solea senegalensis* | | | |
| *Sparus aurata* | | | |

| Spring (n=99) | Summer (n=71) | Autumn (n=56) | Winter (n=61) |
|---|---|---|---|
| *Spicara maena* | | | |
| *Spondyliosoma cantharus* | | | |
| *Symphodus bailloni* | | | |
| *Symphodus melops* | | | |
| *Symphodus ocellatus* | | | |
| *Syngnathus rostellatus* | | | |
| *Syngnathus typhle* | | | |
| *Torpedo marmorata* | | | |
| *Trachurus mediterraneus* | | | |
| *Trachurus picturatus* | | | |
| *Trachurus trachurus* | | | |
| *Zeus faber* | | | |

**Supplementary Table 11.** Summary of PERMANOVA results (based on 999 permutations) assessing seasonal differences in species composition. The table includes Df (degrees of freedom), SumOfSqs (sums of squares), $R^2$ (explained variation), F (F-test value), and p (p-value of the F score). Significant values are indicated with "**".

| Variable | Df | SumOfSqs | $R^2$ | *F* | *p* |
|---|---|---|---|---|---|
| Season | 3 | 0.7972 | 0.3579 | 1.6724 | 0.004** |
| Residual | 9 | 1.43 | 0.6421 | | |
| Total | 12 | 2.2272 | 1 | | |

**Supplementary Table 12**. Number of Sardina pilchardus and Engraulis encrasicolus eggs and their corresponding read counts for each primer set used in high-throughput sequencing across all sampled months. % Total Ich. represents the percentage of eggs from each species relative to the total ichthyoplankton (eggs and larvae) collected in that month. Read counts are provided for each primer set: C2 reads (FishATL_Cocktail2), MiCOI reads (mICOIintF/LoboR1), 12S reads (miFISH U+E), and 16S reads (Fish 16S).

| Month | ***Sardina pilchardus* Data** | | | | | | ***Engraulis encrasicolus* Data** | | | | | |
|---|---|---|---|---|---|---|---|---|---|---|---|---|
| | **Sardine Eggs** | **% Total Ich.** | **C2 Reads** | **MiCOI Reads** | **12S Reads** | **16S Reads** | **Engraulis Eggs** | **% Total Ich.** | **C2 Reads** | **MiCOI Reads** | **12S Reads** | **16S Reads** |
| April | 244 | 27.29 | 586 | 1 | 1081 | 2910 | 20 | 2.24 | 1304 | 643 | 2312 | 2966 |
| May | 354 | 34.27 | 10733 | 9 | 18657 | 33937 | 6 | 0.58 | 8904 | 10531 | 13584 | 19641 |
| June | 687 | 69.96 | 216 | 0 | 52 | 27 | 4 | 0.41 | 1793 | 713 | 850 | 2364 |
| July | 272 | 61.82 | 392 | 0 | 142 | 240 | 18 | 4.09 | 8091 | 3993 | 13199 | 22179 |
| August | 110 | 41.51 | 1 | 0 | 239 | 65 | 2 | 0.75 | 20786 | 9111 | 37001 | 37147 |
| September | 11 | 36.67 | 10 | 0 | 5690 | 4319 | 4 | 13.33 | 1292 | 4888 | 24403 | 5050 |
| October | 46 | 48.94 | 727 | 0 | 15728 | 10346 | 0 | 0.00 | 10598 | 23414 | 43894 | 26287 |
| November | 0 | 0.00 | 1244 | 0 | 10608 | 15098 | 0 | 0.00 | 0 | 2695 | 3 | 26 |
| December | 0 | 0.00 | 13409 | 132 | 65381 | 38288 | 0 | 0.00 | 6101 | 10441 | 17392 | 5467 |
| January | 6 | 25.00 | 1216 | 2 | 7698 | 6757 | 0 | 0.00 | 94 | 793 | 34 | 75 |
| February | 159 | 21.93 | 17936 | 111 | 40712 | 80979 | 0 | 0.00 | 4933 | 2154 | 3883 | 4994 |
| March | 62 | 4.96 | 4028 | 215 | 15623 | 56533 | 0 | 0.00 | 0 | 1 | 14 | 3 |
| April * | 444 | 6.42 | 16367 | 220 | 54159 | 53292 | 400 | 5.78 | 2698 | 3947 | 9896 | 11048 |

**Supplementary Table 13.** Historical records of fish species identified in ichthyoplankton from the Guadiana River Estuary, considering the season of collection and estuarine section, based on findings from previous studies (Chícharo & Teodósio, 1991; Chícharo et al., 2006; Faria et al., 2006).

| **Species** | **Family** | **Spring** | **Summer** | **Autumn** | **Winter** | **Coastal** | **Low Est.** | **Mid Est.** | **High Est.** |
|---|---|---|---|---|---|---|---|---|---|
| *Ammodytes tobianus* | Ammodytidae | | | | X | X | | | |
| *Anguilla anguilla* | Anguillidae | X | | | X | | | X | |
| *Argyrosomus regius* | Sciaenidae | | | | | | | | |
| *Atherina presbyter* | Atherinidae | X | X | | | X | X | X | X |
| *Belone belone* | Belonidae | X | X | | | | | X | |
| *Blennius ocellaris* | Blenniidae | | | | | | | | |
| *Buglossidium luteum* | Soleidae | X | | X | X | | X | | |
| *Callionymus reticulatus* | Callionymidae | X | X | | | X | X | | |
| *Centrolabrus exoletus* | Labridae | X | X | | | | | | |
| *Chelon auratus* | Mugilidae | | | | X | | | X | |
| *Chelon ramada* | Mugilidae | | | | X | | | X | |
| *Coryphoblennius galerita* | Blenniidae | X | | | | X | X | | |
| *Ctenolabrus rupestris* | Labridae | X | X | | | | | | |
| *Cyprinus carpio* | Cyprinidae | | | | | | | | |
| *Dicentrarchus labrax* | Moronidae | X | X | | X | | X | X | |
| *Diplodus bellottii* | Sparidae | | | | | | | | |
| *Diplodus sargus* | Sparidae | X | X | | X | X | X | | |

| Species | Family | Spring | Summer | Autumn | Winter | Coastal | Low Est. | Mid Est. | High Est. |
|---|---|---|---|---|---|---|---|---|---|
| *Diplodus vulgaris* | Sparidae | | X | | | | X | | |
| *Echiichthys vipera* | Trachinidae | X | X | | X | X | | | |
| *Engraulis encrasicolus* | Engraulidae | X | X | | X | X | X | X | X |
| *Entelurus aequoreus* | Syngnathidae | X | | | | | | | |
| *Gobius niger* | Gobiidae | X | X | | | X | X | X | X |
| *Halobatrachus didactylus* | Batrachoididae | X | X | | X | X | X | X | |
| *Hippocampus guttulatus* | Syngnathidae | X | | | | | | | |
| *Hippocampus hippocampus* | Syngnathidae | X | X | | | | | X | |
| *Labrus bergylta* | Labridae | | | | | | | | |
| *Lipophrys pholis* | Blenniidae | X | | | | | | | |
| *Luciobarbus comizo* | Cyprinidae | X | X | | X | | | | X |
| *Luciobarbus microcephalus* | Cyprinidae | | X | | | | | | X |
| *Luciobarbus sclateri* | Cyprinidae | X | X | | X | | | X | X |
| *Monochirus hispidus* | Soleidae | X | | X | X | X | | | |
| *Mullus surmuletus* | Mullidae | | X | | | X | | | |
| *Nerophis ophidion* | Syngnathidae | X | X | | | | X | X | X |
| *Parablennius gattorugine* | Blenniidae | X | X | | X | X | X | X | |
| *Parablennius pilicornis* | Blenniidae | X | X | | X | X | X | X | |
| *Pegusa lascaris* | Soleidae | X | X | | X | X | X | X | |
| *Pomatoschistus microps* | Gobiidae | X | X | X | X | X | X | X | X |
| *Pomatoschistus minutus* | Gobiidae | X | X | X | X | X | X | X | X |
| *Pomatoschistus pictus* | Gobiidae | X | X | X | X | X | X | X | X |
| *Sardina pilchardus* | Alosidae | X | X | X | X | X | X | X | X |
| *Sarpa salpa* | Sparidae | | X | | | X | X | | |
| *Serranus hepatus* | Serranidae | | X | | | | | | |
| *Solea senegalensis* | Soleidae | X | X | | X | X | X | | |
| *Solea solea* | Soleidae | X | | | X | X | | | |
| *Sparus aurata* | Sparidae | | | | | | | | |
| *Spondyliosoma cantharus* | Sparidae | X | | | X | | | | |
| *Symphodus bailloni* | Labridae | X | X | X | | X | X | | |
| *Symphodus melops* | Labridae | X | X | | | | X | | |
| *Syngnathus abaster* | Syngnathidae | X | X | X | X | | X | X | X |
| *Syngnathus acus* | Syngnathidae | X | X | | | | X | X | X |
| *Syngnathus rostellatus* | Syngnathidae | | | | | | | | |
| *Syngnathus typhle* | Syngnathidae | X | X | | | X | | X | |
| *Trigla lyra* | Triglidae | X | | | | | X | | |
